# Genome-Wide Asymptomatic B-Cell, CD4^+^ and CD8^+^ T-Cell Epitopes, that are Highly Conserved Between Human and Animal Coronaviruses, Identified from SARS-CoV-2 as Immune Targets for Pre-Emptive Pan-Coronavirus Vaccines

**DOI:** 10.1101/2020.09.27.316018

**Authors:** Swayam Prakash, Ruchi Srivastava, Pierre-Gregoire Coulon, Nisha R. Dhanushkodi, Aziz A. Chentoufi, Delia F. Tifrea, Robert A. Edwards, Cesar J. Figueroa, Sebastian D. Schubl, Lanny Hsieh, Michael J. Buchmeier, Mohammed Bouziane, Anthony B. Nesburn, Baruch D. Kuppermann, Lbachir BenMohamed

**Affiliations:** Laboratory of Cellular and Molecular Immunology, Gavin Herbert Eye Institute, University of California Irvine, School of Medicine, Irvine, CA 92697; Department of Pathology and Laboratory Medicine, School of Medicine, Irvine, CA 92697; Department of Surgery, Divisions of Trauma, Burns & Critical Care, School of Medicine, Irvine, CA 92697; Department of Medicine, Division of Infectious Diseases and Hospitalist Program, School of Medicine, Irvine, CA 92697; Center for Virus Research, and Division of Infectious Disease,, University of California School of Medicine, Irvine, CA, 92697; Sunomix Therapeutics, Inc., San Diego, CA 92121; Institute for Immunology; University of California Irvine, School of Medicine, Irvine, CA 92697, USA

**Author notes:** Corresponding Author: Professor Lbachir BenMohamed, Laboratory of Cellular and Molecular Immunology, Gavin Herbert Eye Institute, University of California Irvine, School of Medicine, Hewitt Hall, Room 2032; 843 Health Sciences Rd.; Irvine, CA 92697-4390. Contributed equally to this study (Swayam Prakash, Ruchi Srivastava, Pierre-Gregoire Coulon, Nisha R. Dhanushkodi).

**Keywords:** SARS-CoV-2, SL-CoVs, COVID-19, Pan-Coronavirus, Vaccine, Epitopes, Antibodies, CD4^+^ T cells, CD8^+^ T cells

## Abstract

Over the last two decades, there have been three deadly human outbreaks of Coronaviruses (CoVs) caused by emerging zoonotic CoVs: SARS-CoV, MERS-CoV, and the latest highly transmissible and deadly SARS-CoV-2, which has caused the current COVID-19 global pandemic. All three deadly CoVs originated from bats, the natural hosts, and transmitted to humans *via* various intermediate animal reservoirs. Because there is currently no universal pan-Coronavirus vaccine available, two worst-case scenarios remain highly possible: (1) SARS-CoV-2 mutates and transforms into a seasonal “flu-like” global pandemic; and/or (2) Other global COVID-like pandemics will emerge in the coming years, caused by yet another spillover of an unknown zoonotic bat-derived SARS-like Coronavirus (SL-CoV) into an unvaccinated human population. Determining the antigen and epitope landscapes that are conserved among human and animal Coronaviruses as well as the repertoire, phenotype and function of B cells and CD4^+^ and CD8^+^ T cells that correlate with resistance seen in asymptomatic COVID-19 patients should inform in the development of pan-Coronavirus vaccines ^1^. In the present study, using several immuno-informatics and sequence alignment approaches, we identified several human B-cell, CD4^+^ and CD8^+^ T cell epitopes that are highly conserved in: (*i*) greater than 81,000 SARS-CoV-2 human strains identified to date in 190 countries on six continents; (*ii*) six circulating CoVs that caused previous human outbreaks of the “Common Cold”; (*iii*) five SL-CoVs isolated from bats; (*iv*) five SL-CoV isolated from pangolins; (*v*) three SL-CoVs isolated from Civet Cats; and (*vi*) four MERS strains isolated from camels. Furthermore, we identified cross-reactive asymptomatic epitopes that: (*i*) recalled B cell, CD4^+^ and CD8^+^ T cell responses from both asymptomatic COVID-19 patients and healthy individuals who were never exposed to SARS-CoV-2; and (*ii*) induced strong B cell and T cell responses in “humanized” Human Leukocyte Antigen (HLA)-DR/HLA-A*02:01 double transgenic mice. The findings herein pave the way to develop a pre-emptive multi-epitope pan-Coronavirus vaccine to protect against past, current, and potential future outbreaks.

## INTRODUCTION

As deforestation continues to expand and humans progressively conquer wildlife habitats around the globe, the wildlife “fights back” by spilling over many zoonotic viruses into human populations ^2, 3^. Among these, is the large family of RNA viruses called Coronaviruses (a name alluding to the small pointy spikes that give them the appearance of a corona). Since the first human Coronavirus was identified in 1965, many additional strains of Coronaviruses have continued to emerge ^4–6^. These set the stage for several major human disease outbreaks within the last two decades (i.e., from 2002 to 2019): SARS-CoV ^7^; CoV-NL63 ^8^; CoV-HKU1 ^9^; CoV-229E ^9^; CoV-OC43^10^; MERS-CoV ^11^; and the highly contagious and deadly SARS-CoV-2 ^12, 13^ The many deadly Coronavirus outbreaks that occurred in the past 20 years should have been the impetus for urgently developing a pre-emptive pan-Coronavirus vaccine.

The first two deadly Coronaviruses, the MERS-CoV and the SARS-CoV, originated from bats, as natural hosts and reservoirs, and were transmitted to humans from intermediate animals namely camels and civet-cats, respectively ^11, 14–17^. The third highly contagious and deadly SARS-CoV-2, appears to be 96% identical to a bat SARS-like Coronavirus (SL-CoV) strain, termed Bat-CoV-RaTG13, and transmitted to humans from a yet-to-be determined intermediate animal ^18, 19^. Whether SARS-CoV-2 virus is actually a man-made “artificial” recombinant virus is still currently being debated^20^. While animal-to-human spread of ‘‘common cold’’ Coronaviruses occurs frequently, only rarely do human-to-human Coronavirus transmissions occur. However, the highly contagious SARS-CoV-2 successfully produces both animal-to-human spread ^21^ and human-to-human transmission ^22^. As the first known human-to-human transmission of SARS-CoV-2, which causes Coronavirus disease-2019 (COVID-19), was reported in late January 2020, the WHO and US authorities declared a public health emergency ^23^.

Within 2-14 days after SARS-CoV-2 exposure, newly infected individuals may develop fever, fatigue, myalgia and respiratory symptoms including cough and shortness of breath ^24, 25^. While the majority (80-85%) of newly infected individuals are asymptomatic, a minority of individuals are symptomatic, especially the elderly and those with comorbidities ^24, 25^. They develop severe pulmonary inflammatory disease and may need a rapid medical intervention to prevent acute respiratory distress syndrome and death ^24, 25^. Except for Remdesivir, a viral RNA nucleoside analog, none of the existing anti-viral drugs or repositioned drugs appear to have a significantly positive effect on COVID-19, especially during the late stages of infection and illness, prompting an urgent need for alternative measures, such as vaccines to produce a long-lasting immunity. The SARS-CoV-2 infection induces antiviral CD4^+^ T cells, helping the production of neutralizing/blocking antibodies and the formation of effector IFN-γ-producing CD4^+^ T cells and cytotoxic CD8^+^ T cells, all arms of immunity critical in reducing viral load in the majority of asymptomatic and convalescence patients ^26–31^. While SARS-CoV-2-specific antibody and CD4^+^ and CD8^+^ T cells are critical to reducing viral infection in a majority of asymptomatic individuals ^32^, an excessive proinflammatory cytokine storm appears to lead to acute respiratory distress syndrome and death in many symptomatic individuals ^33–39^. Thus, it is crucial to determine the B cell and T cell epitopes landscape that is conserved among past, present and future human and animal Coronavirus strains and isolates along with the B cell and T cell-epitope-specificities in asymptomatic COVID-19 patients, the repertoire, the phenotype and function of B cells and CD4^+^ and CD8^+^ T cells that are associated with natural resistance seen in asymptomatic patients. This information should guide the development of pan-Coronavirus vaccines ^1^.

In the present study, we identified several asymptomatic human B, CD4^+^ and CD8^+^ T cell epitopes that are highly conserved among many strains of the six Coronaviruses previously reported to infect humans and over 81,000 strains of SARS-CoV-2 that currently circulate in 190 countries on six continents. Moreover, as immune targets for pre-emptive pan-coronavirus vaccines, we identified the epitopes that are common among the above human Coronaviruses and twenty-five animal strains isolated from bats, pangolins, civet cats; and camels. We demonstrated the antigenicity of these epitopes in both asymptomatic SARS-CoV-2 patients and unexposed healthy individuals; and their immunogenicity in “humanized” Human Leukocyte Antigen (HLA)-DR/HLA-A*02:01 double transgenic mice. Our findings pave the way for incorporating these highly conserved asymptomatic B-cell and T-cell epitopes in pre-emptive multi-epitope pan-Coronavirus vaccines that would be expected to, not only protect against COVID-19, but also provide a measure of defense against the next global outbreak.

## RESULTS

### 1. Evolutionary convergence of human SARS-CoV-2 into bat and pangolin-derived SARS-like Coronaviruses

Understanding the animal origins of SARS-CoV-2 is critical for the development of a pre-emptive pan-Coronavirus vaccine to protect from future human outbreaks and deter future zoonosis.

We first screened for the evolutionary relationship among human SARS-CoV-2 and SARS-CoV/MERS-CoV strains from previous outbreaks (i.e. Urbani, MERS-CoV, OC43, NL63, 229E, HKU1-genotype-B) along with 25 SARS-like Coronaviruses genome sequence (SL-CoVs) obtained from different animal species: Bats (*Rhinolophus affinis* and *Rhinolophus malayanus*), civet cats (*Paguma larvata*) and pangolins (*Manis javanica*), and MERS-CoVs from camels (*Camelus dromedarius* and *Camelus bactrianus*) (**Fig. 1**). This sequence alignments revealed similarity of the original human-SARS-CoV-2 strain found in Wuhan to four bat SL-CoV strains: hCoV-19-bat-Yunnan-RmYN02, bat-CoV-19-ZXC21, and hCoV-19-bat-Yunnan-RaTG13 obtained from the Yunnan and Zhejiang provinces of China (**Fig. 1A**). With further genetic distance analysis we discovered the least evolutionary divergence between SARS-CoV-2 isolate Wuhan-Hu-1 and the above mentioned three SL-CoVs isolates from bats, namely: (1) the Bat-CoV-RaTG13 (0.1), (2) bat-CoV-19-ZXC21 (0.1) and (3) the Bat-CoV-YN02 (0.2) (**Fig. 1B** and **1C**). Moreover, the phylogenetic analysis performed among the whole genome sequences of a total of 81,963 SARS-CoV-2 strains for which sequences have been reported in circulation in 190 countries suggest an evolutionary convergence of bat and pangolin SL-CoVs into the human SARS-CoV-2 strains (**Figs. 1D** and **1E**). Furthermore, through a complete genome tree derived from the 81,963 SARS-CoV-2 genome sequences submitted from Asian, African, North American, South American, European, and Oceanian regions, we confirmed that the least evolutionary divergence for SARS-CoV-2 strains are in SL-CoVs isolated from bats and pangolins (**Figs. 1D, 1E** and **1F**).

**Figure 1.**
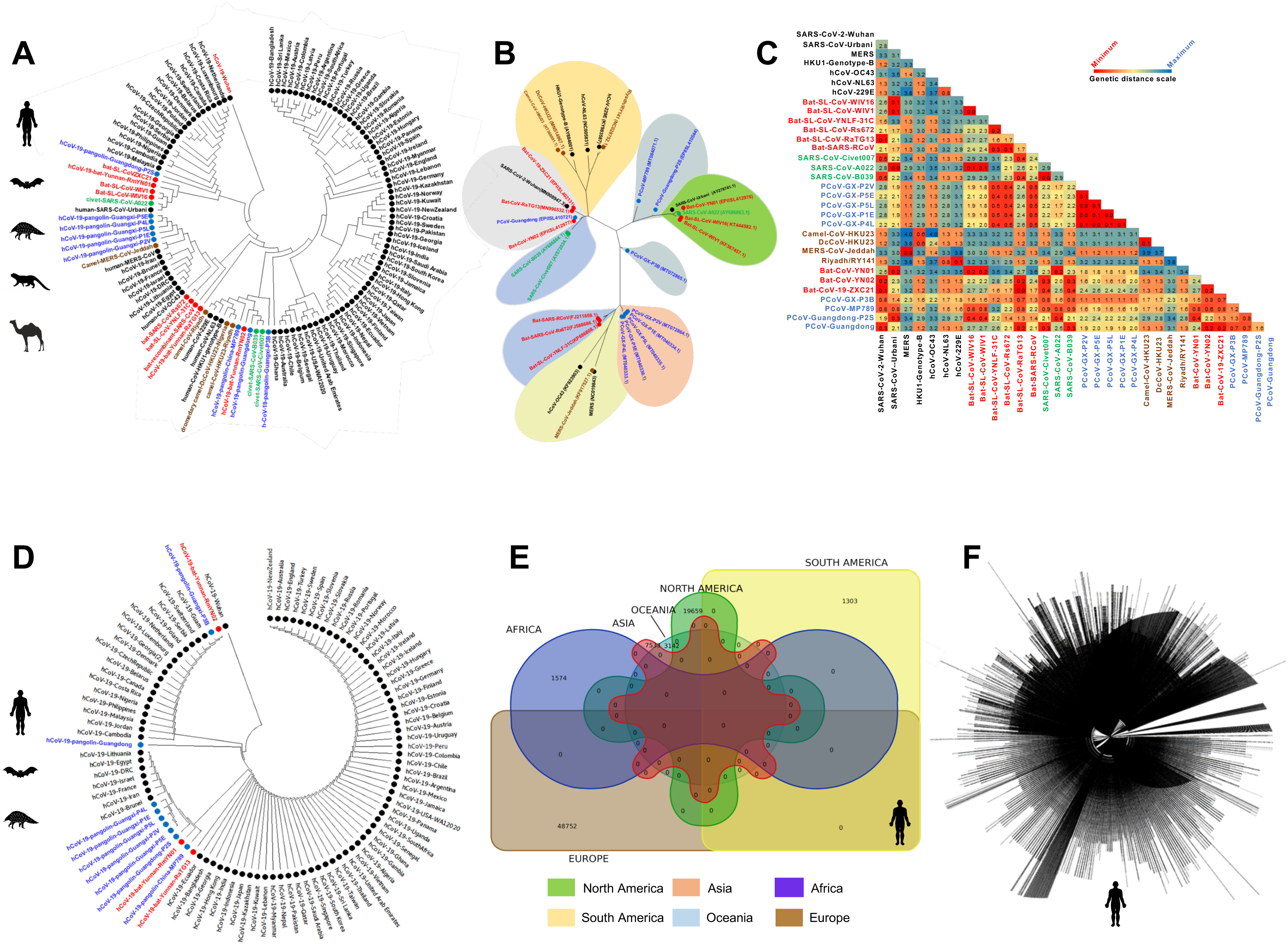
Evolutionary comparison of genome sequences among beta-Coronavirus strains isolated from humans and animals: (**A**) *Left panel:* Phylogenetic analysis performed between SARS-CoV-2 strains (obtained from humans (*Homo Sapiens (black)*), along with the animal’s SARS-like Coronaviruses genome sequence (SL-CoVs) sequences obtained from bats (*Rhinolophus affinis*, *Rhinolophus malayanus (red)*), pangolins (*Manis javanica (blue)*), civet cats (*Paguma larvata (green)),* and camels (*Camelus dromedaries (Brown)*). The included SARS-CoV/MERS-CoV strains are from previous outbreaks (obtained from humans (Urbani, MERS-CoV, OC43, NL63, 229E, HKU1-genotype-B), bats (WIV16, WIV1, YNLF-31C, Rs672, recombinant strains), camel (*Camelus dromedaries,* (KT368891.1, MN514967.1, KF917527.1, NC_028752.1), and civet (Civet007, A022, B039)). The human SARS-CoV-2 genome sequences are represented from six continents. (**B**) Phylogenetic analysis performed among SARS-CoV-2 strains from human and other species with previous strains of SARS/MERS-CoV showed minimum genetic distance between the first SARS-CoV-2 isolate Wuhan-Hu-1 reported from the Wuhan Seafood market with bat strains hCoV-19-bat-Yunnan-RmYN02, bat-CoV-19-ZXC21, and hCoV-19-bat-Yunnan-RaTG13. This makes the bat strains nearest precursor to the human-SARS-CoV-2 strain. (**C**) Genetic distances based on Maximum Composite Likelihood model among the human, bat, pangolin, civet cat and camel genome sequences. Results indicate least genetic distance among SARS-CoV-2 isolate Wuhan-Hu-1 and bat strains bat-CoV-19-ZXC21 (0.1), hCoV-19-bat-Yunnan-RaTG13 (0.1), and hCoV-19-bat-Yunnan-RmYN02 (0.2). (**D**) Evolutionary analysis performed among the human-SARS-CoV-2 genome sequences reported from six continents and SARS-CoV-2 genome sequences obtained from bats (*Rhinolophus affinis*, *Rhinolophus malayanus*), and pangolins (*Manis javanica*)) (**E**) Venn diagram showing the number of SARS-CoV-2 genome sequences reported from Africa (*n* = 1574), Asia (*n* = 7533), North America (*n* = 19659), South America (*n* = 1303), Europe (*n* = 48752), and Oceania region (*n* = 3142) as on August 18, 2020. (**F**) Complete genome tree derived from 81963 outbreak SARS-CoV-2 genome sequences submitted from Asian, African, North American, South American, European, and Oceanian regions.

Altogether, the phylogenetic analysis and genetic distance suggest that the highly contagious and deadly human-SARS-CoV-2 strain originated from bats, likely from either the Bat-CoV-19-ZXC21 or Bat-CoV-RaTG13 strains, that spilled over into humans after further mutations and/or recombination with a yet-to-be determined CoV strain from the pangolin, as the likely intermediate animal reservoir. These mutations and/or recombination possibly contributed to the rapid global expansion of the highly contagious and deadly SARS-CoV-2 ^40, 41^.

### 2. Genome-wide identification of SARS-CoV-2 CD8^+^ T cell epitopes that are highly conserved between human and bat/pangolin Coronaviruses

We first predicted potential CD8^+^ T cell epitopes from the entire genome sequence of the first SARS-CoV-2-Wuhan-Hu-1 strain (NCBI GeneBank accession number MN908947.3) ^42–48^. For this, we used multiple databases and algorithms including the SYFPEITHI, MHC-I processing predictions, MHC-I binding predictions, MHC-I immunogenicity and Immune Epitope Database (IEDB) ^45, 49^. We focused on epitopes that are restricted to the 5 most frequent human leukocyte antigen (HLA) class I alleles with large coverage in worldwide human populations, regardless of race and ethnicity (i.e. HLA-A*01:01, HLA-A*02:01, HLA-A*03:01, HLA-A*11:01, HLA-A*23:01) ^50–52^ (**Supplemental Figs. S1A** and **S1C**). Using the above criteria, we originally identified a total of 9,660 potential CD8^+^ T cell epitopes derived from 12 SARS-CoV-2 open-reading-frames (ORFs) of SARS-CoV-2-Wuhan-Hu-1 strain (**Supplemental Table S1**). Subsequently, this large pool of epitopes was narrowed to 91 epitopes, that are highly conserved among: (*i*) over 81,000 SARS-CoV-2 strains (that currently circulate in 190 countries on 6 continents); (*ii*) the 4 major ‘‘common cold’’ Coronaviruses that caused previous outbreaks (i.e. CoV-OC43, CoV-229E, CoV-HKU1 genotype B, and CoV-NL63); and (*iii*) the SL-CoVs that are isolated from bats, civet cats, pangolins and camels (**Fig. 2A**). While the highest degree of similarity (expressed as % of resemblance) was identified among 81,963 SARS-CoV-2 strains, six strains of previous human SARS-CoVs and animal SL-CoVs strains isolated from bats and pangolins, only a small percentage of similarity was found between strains of SARS-CoV-2 and MERS-CoV strains (**Fig. 2D**). However, a significantly lower degree of similarity was recorded among SARS-CoV-2 strains and the SL-CoVs isolated from civet cats’ and camels’ CoVs (**Fig. 2D**).

**Figure 2:**
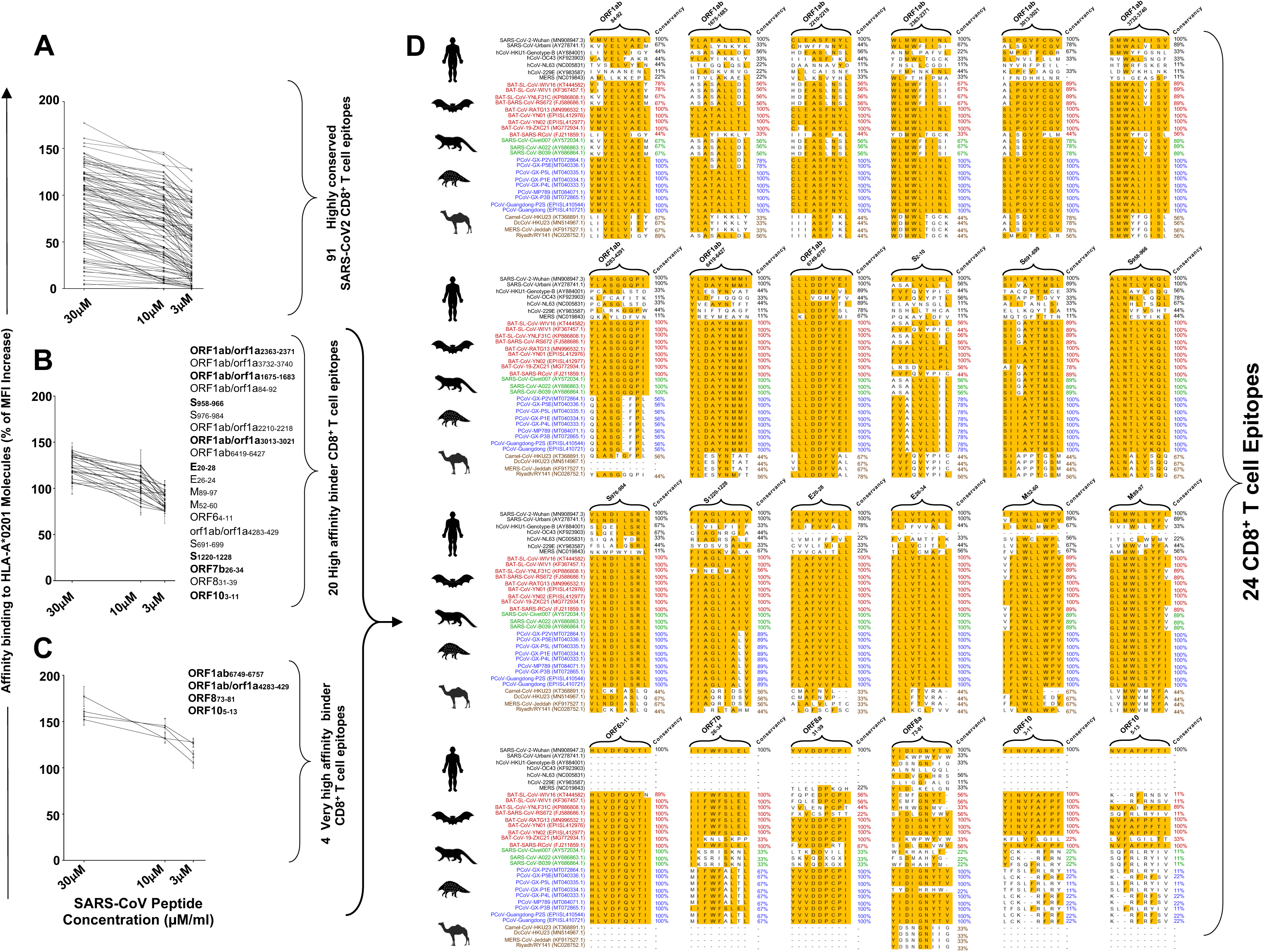
Identification of highly conserved potential SARS-CoV-2-derived human CD8^+^ T cell epitopes that bind with high affinity to HLA-A*02:01 molecules: (**A**) Ninety one, genome-wide *In-silico* predicted, and highly conserved SARS-CoV-2-derived CD8^+^ T cell epitope peptides were synthetized and were tested for their binding affinity *in vitro* to HLA-A*02:01 molecules expressed on the surface of T_2_ cells. (**B**) Out of the 91 CD8^+^ T cell epitopes, 20 epitopes bind with high affinity and stabilized the expression of HLA-A*0201 molecules on the surface of the T_2_ cells. (**C**) Four epitopes were subsequently selected with the highest affinity binding to HLA-A*02:01 molecules even at the lowest molarity of 3 uM. The levels of HLA-A*02:01 surface expression was determined by mean fluorescence intensity (MFI), measured by flow cytometry on T_2_ cells following an overnight incubation of T_2_ cells at 26°C with decreasing peptide epitopes molarity (30, 15 and 5μM) as shown in graphs. Percent MFI increase was calculated as follows: Percent MFI increase = (MFI with the given peptide - MFI without peptide) / (MFI without peptide) X 100. (**D**) Conservancy of the 24 high binder CD8^+^ T cell epitopes among human and animal Coronaviruses. Shown are the comparison of sequence homology for the 24 CD8^+^ T cell epitopes among 81,963 SARS-CoV-2 strains (that currently circulate in 190 countries on 6 continents), the 4 major ‘‘common cold’’ Coronaviruses that cased previous outbreaks (i.e. CoV-OC43, CoV-229E, CoV-HKU1, and CoV-NL63), and the SL-CoVs that were isolated from bats, civet cats, pangolins and camels. Epitope sequences highlighted in yellow present high degree of homology among the currently circulating 81,963 SARS-CoV-2 strains and at least a 50% conservancy among two or more humans SARS-CoV strains from previous outbreaks, and the SL-CoV strains isolated from bats, civet cats, pangolins and camels, as described in *Materials* and *Methods*. *Homo Sapiens-black*, bats (*Rhinolophus affinis*, *Rhinolophus malayanus - red*), pangolins (*Manis javanica-blue*), civet cats (*Paguma larvata-green*), and camels (*Camelus dromedaries-brown*).

We further identified 24 SARS-CoV-2 human CD8^+^ T cell epitopes, out of the 91 epitopes, that bind with high affinity with HLA-A*02:01 molecules, using *in vitro* peptide-HLA binding assay (**Fig. 2B**). Four epitopes were found to be very high affinity binders (**Fig. 2C**). The 24 epitopes with high binding affinity were later confirmed *in silico* using molecular docking models across 5 major HLA-A*01:01, HLA-A*02:01, HLA-A*03:01, HLA-A*11:01, HLA-A*23:01 haplotypes (**Supplemental Fig. S2**) ^53^. The highest binding affinity to HLA-A*02:01 molecules, with the highest interaction similarity (S_inter_) scores (*blue squares*), were recorded for ORF1ab_6749-6757_, S_2-10_, S_958-966_, S_1220-1228_, E_26-34_, ORF8_83-91_, ORF10_3-11_ and ORF10_5-13_ whereas minimum S_inter_ score was observed for ORF1ab_3732-3740_, S_691-699_ and M_89-97_. Other CD8^+^ T cell epitopes like ORF1ab_1675-1683_, ORF1ab_2363-2371_, ORF1ab_3013-3021_ and ORF7b_26-34_ were also found with intermediate S_inter_ scores (**Supplemental Figs. S2A** and **S2C**). While the identified highly conserved CD8^+^ T cell epitopes were distributed within 8 of the 12 structural and non-structural ORFs (i.e. ORF1ab, S, E, M, ORF6, ORF7b, ORF8, and ORF10), the highest numbers of epitopes were localized in the replicase polyprotein 1ab/1a (ORF1ab) (9 epitopes) followed by the spike glycoprotein (S) (5 epitopes) (**Fig. 2D** and **Fig. 12**).

Altogether, our findings identified 24 highly conserved potential human CD8^+^ T cell epitopes from the sequence of SARS-CoV-2 that are highly conserved among 81,963 SARS-CoV-2 strains, the 4 major ‘‘common cold’’ Coronaviruses (i.e. CoV-OC43, CoV-229E, CoV-HKU1 genotype B, and CoV-NL63), and several SL-CoV strains that are isolated from bats and pangolins. Our results also suggested that the replicase polyprotein ORF1ab and the spike glycoprotein could be the most immunodominant antigens, that cross-react between human and animal Coronaviruses, and that is targeted by CD8^+^ T cells from both SARS-CoV-2 and ‘‘common cold’’ Coronaviruses infected humans.

### 3. In silico screening of potential promiscuous SARS-CoV-2 CD4^+^ T cell epitopes that are highly conserved between human and bat/pangolin Coronaviruses

We subsequently identified a total of 9,594 potential HLA-DR-restricted CD4^+^ T cell epitopes from the whole genome sequence of SARS-CoV-2-Wuhan-Hu-1 strain (MN908947.3) using multiple databases and algorithms including the SYFPEITHI, MHC-II Binding Predictions, Tepitool and TEPITOPEpan (**Supplemental Table S2**). These potential promiscuous CD4^+^ T cell epitopes were screened *in silico* against the five most frequent HLA-DR alleles with large coverage in the human population, regardless of race or ethnicity: HLA-DRB1*01:01, HLA-DRB1*11:01, HLA-DRB1*15:01, HLA-DRB1*03:01, HLA-DRB1*04:01 (**Supplemental Figs. S1B** and **S1D**). The number of potential CD4^+^ T cell epitopes was later narrowed down to 16 epitopes based on: (*i*) the epitope sequences that are highly conserved among 81,963 SARS-CoV-2 strains, the 4 major ‘‘common cold’’ and 25 SL-CoV strains isolated from bats, civet cats, pangolins and camels (**Fig. 3**); and (*ii*) their high binding affinity to HLA-DR molecules using *in silico* molecular docking models (**Supplemental Fig. S3**). The sequences of most of the 16 CD4^+^ T cell epitopes are 100% conserved and common among 81,963 SARS-CoV-2 strains currently circulating in 6 continents (**Fig. 3**). High degree of sequence similarities were also identified in the sequences of most 16 CD4^+^ T cell epitopes among the SARS-CoV-2 strains and the six strains of previous human SARS-CoVs (e.g. up to 100 % sequence identity for epitopes ORF1ab_5019-5033_, ORF1ab_6088-6102_, ORF1ab _6420-6434_, E_20-34_, E_26-40_ and M_176-190_). Moreover, high degree of sequence similarities were also identified among the SARS-CoV-2 strains and the SL-CoV strains isolated from bats and pangolins. In contrast, a low sequence similarity, of around 20%-40%, was identified among CD4^+^ T cell epitopes from SARS-CoV-2 strains and the SL-CoV strains isolated from civet cats and the MERS-like CoV strains isolated from camels (**Figs. 3** and **12**).

**Figure 3:**
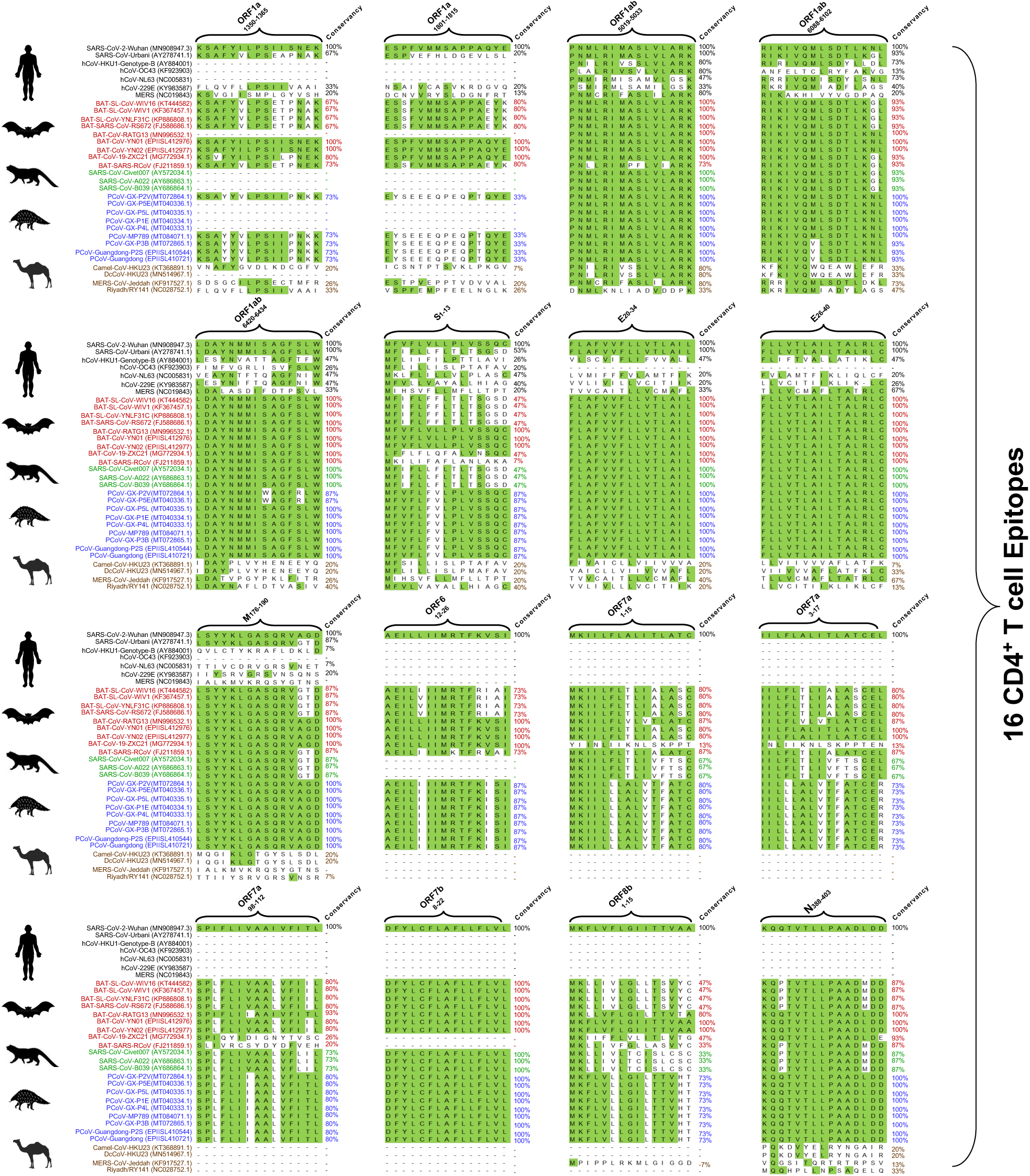
Identification of highly conserved potential SARS-CoV-2-derived human CD4^+^ T cell epitopes that bind with high affinity to HLA-DR molecules: Out of a total of 9,594 potential HLA-DR-restricted CD4^+^ T cell epitopes from the whole genome sequence of SARS-CoV-2-Wuhan-Hu-1 strain (NC-045512.2), 16 epitopes that bind with high affinity to HLA-DRB1 molecules were selected. The conservancy of the 16 CD4^+^ T cell epitopes was analyzed among human and animal Coronaviruses. Shown are the comparison of sequence homology for the 16 CD4^+^ T cell epitopes among 81,963 SARS-CoV-2 strains (that currently circulate in 6 continents), the 4 major ‘‘common cold’’ Coronaviruses that cased previous outbreaks (i.e. CoV-OC43, CoV-229E, CoV-HKU1, and CoV-NL63), and the SL-CoVs that were isolated from bats, civet cats, pangolins and camels. Epitope sequences highlighted in green present high degree of homology among the currently circulating 81,963 SARS-CoV-2 strains and at least a 50% conservancy among two or more humans SARS-CoV strains from previous outbreaks, and the SL-CoV strains isolated from bats, civet cats, pangolins and camels, as described in *Materials* and *Methods*. *Homo Sapiens-black*, bats (*Rhinolophus affinis*, *Rhinolophus malayanus-red*), pangolins (*Manis javanica-blue*), civet cats (*Paguma larvata-green*), and camels (*Camelus dromedaries-brown*).

The 16 highly conserved CD4^+^ T cell epitopes are distributed within 9 out of the 12 structural and non-structural ORFs (i.e. ORF1ab, S, E, M, ORF6, ORF7a, ORF7b, ORF8 and N). The highest numbers of epitopes were localized in the replicase polyprotein ORF1ab/1a (5 epitopes) followed by ORF7a (3 epitopes) (**Figs. 3** and **12**). Unlike the human CD8^+^ T cell epitopes, the human CD4^+^ T cell epitopes are found to be expressed in each of the structural S, E, M, and N proteins. Two epitopes are from the envelope protein (E), 1 epitope from the membrane protein (M), 1 epitope from the nucleoprotein (N) protein, and 1 epitope from the spike protein (S). The remaining CD4^+^ T cell epitopes are distributed among the ORF6, ORF7a, ORF7b and ORF8 proteins (**Figs. 3** and **12**).

Altogether, these results identified 16 potential CD4^+^ T cell epitopes from the whole sequence of SARS-CoV-2 that cross-react and have high sequence similarity among 81,963 SARS-CoV-2 strains, the main 4 major ‘‘common cold’’ Coronaviruses and the SL-CoV strains isolated from bats and pangolins. Similar to CD8^+^ T cell epitopes, the replicase polyprotein ORF1ab appeared to be the most immunodominant antigen with high number of conserved epitopes that maybe targeted by human CD4^+^ T cells.

### 4. Cross-reactive human and animal Coronavirus-derived epitopes recalled memory CD4^+^ and CD8^+^ T cells from SARS-CoV-2 patients and unexposed healthy individuals and were immunogenic in “humanized” HLA transgenic mice

Next, we assessed whether the potential SARS-CoV-2 CD4^+^ and CD8^+^ T cell epitopes, that are highly conserved between human and animal Coronaviruses, would recall memory CD8^+^ T cells from COVID-19 patients as well as from healthy individuals, who have never been exposed to SARS-CoV-2 or to COVID-19 patients (i.e. from healthy individuals blood samples that were collected from 2014 to 2018, **Figs. 4A** and **5A**). Detailed clinical and demographic characteristics of the COVID-19 patients and the unexposed healthy individuals enrolled in the present study, with respect to age, gender, HLA-A*02:01 and HLA-DR distribution, COVID-19 disease severity, comorbidity and biochemical parameters are described in **Table 1** and in *Materials* and *Methods*.

**Figure 4:**
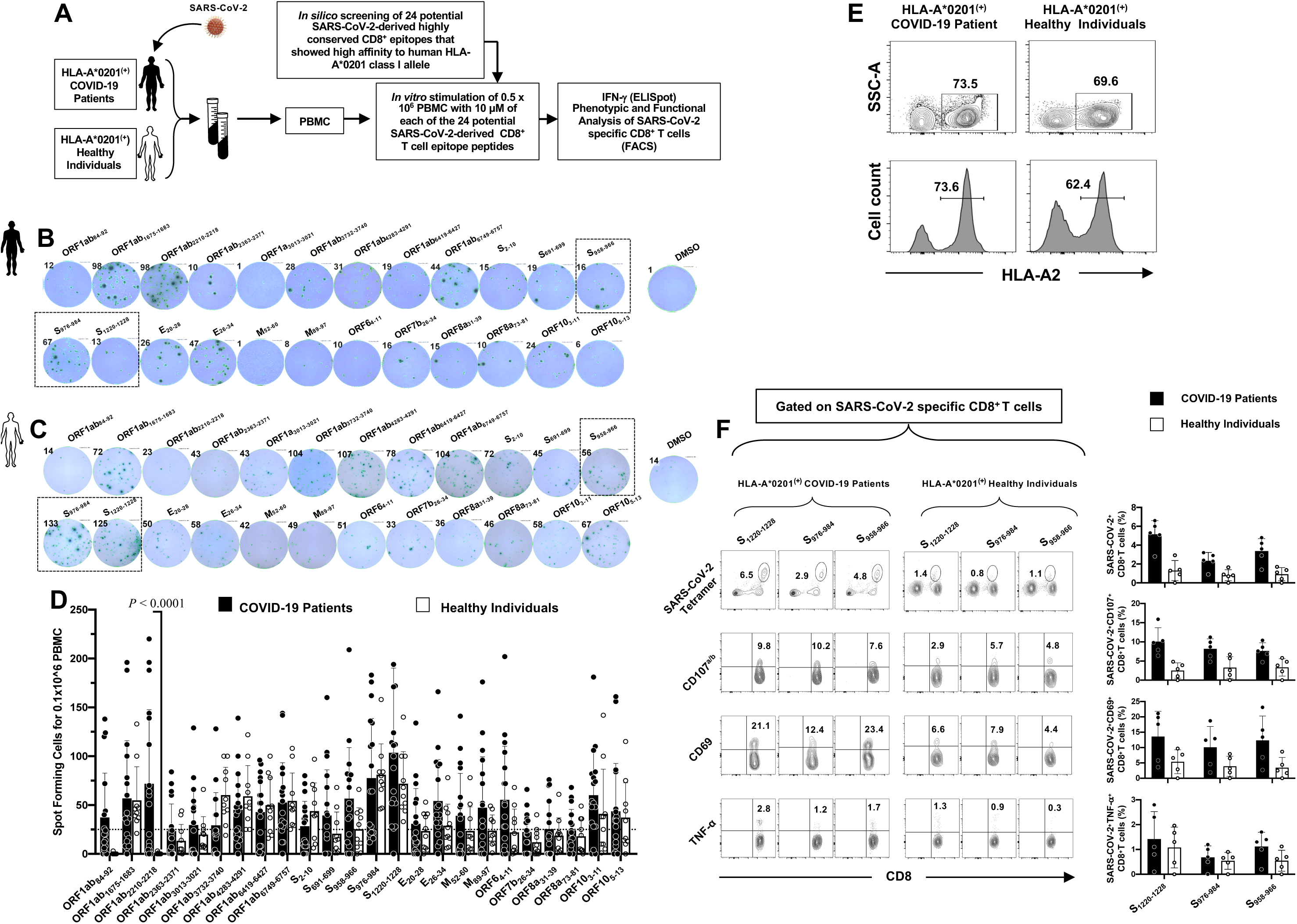
CD8^+^ T cells specific to highly conserved SARS-CoV-2 epitopes detected in COVID-19 patients and unexposed healthy individuals: (**A**) Experimental design: PBMCs from HLA-A2 positive COVID-19 patients (*n* = 62) (**B**) and controls unexposed healthy individuals (*n* = 10) (**C**) were isolated and stimulated overnight with 10 uM of each of the 24 SARS-CoV-2-derived CD8^+^ T cell epitope peptides. The number of IFN-γ-producing cells were quantified using ELISpot assay (**B, C** and **D**). PBMCs from HLA-A0201 positive COVID-19 patients (**E**) were further stimulated for an additional 5 hours in the presence of mAbs specific to CD107^a^ and CD107^b^, and golgi-plug and golgi-stop. Tetramers specific to Spike epitopes, CD107^a/b^ and CD69 and TNF-α expression were then measured by FACS. Representative FACS plot showing the frequencies of Tetramer^+^CD8^+^ T cells, CD107^a/b+^CD8^+^ T cells, CD69^+^CD8^+^ T cells and TNF-α^+^CD8^+^ T cells following priming with a group of 4 Spike CD8^+^ T cell epitope peptides (**F**, *left panel*). Average frequencies of tetramer^+^CD8^+^ T cells, CD107^a/b+^CD8^+^ T cells, CD69^+^CD8^+^ T cells and TNF-α^+^CD8^+^ T cells (**F**, *right panel*).

**Figure 5:**
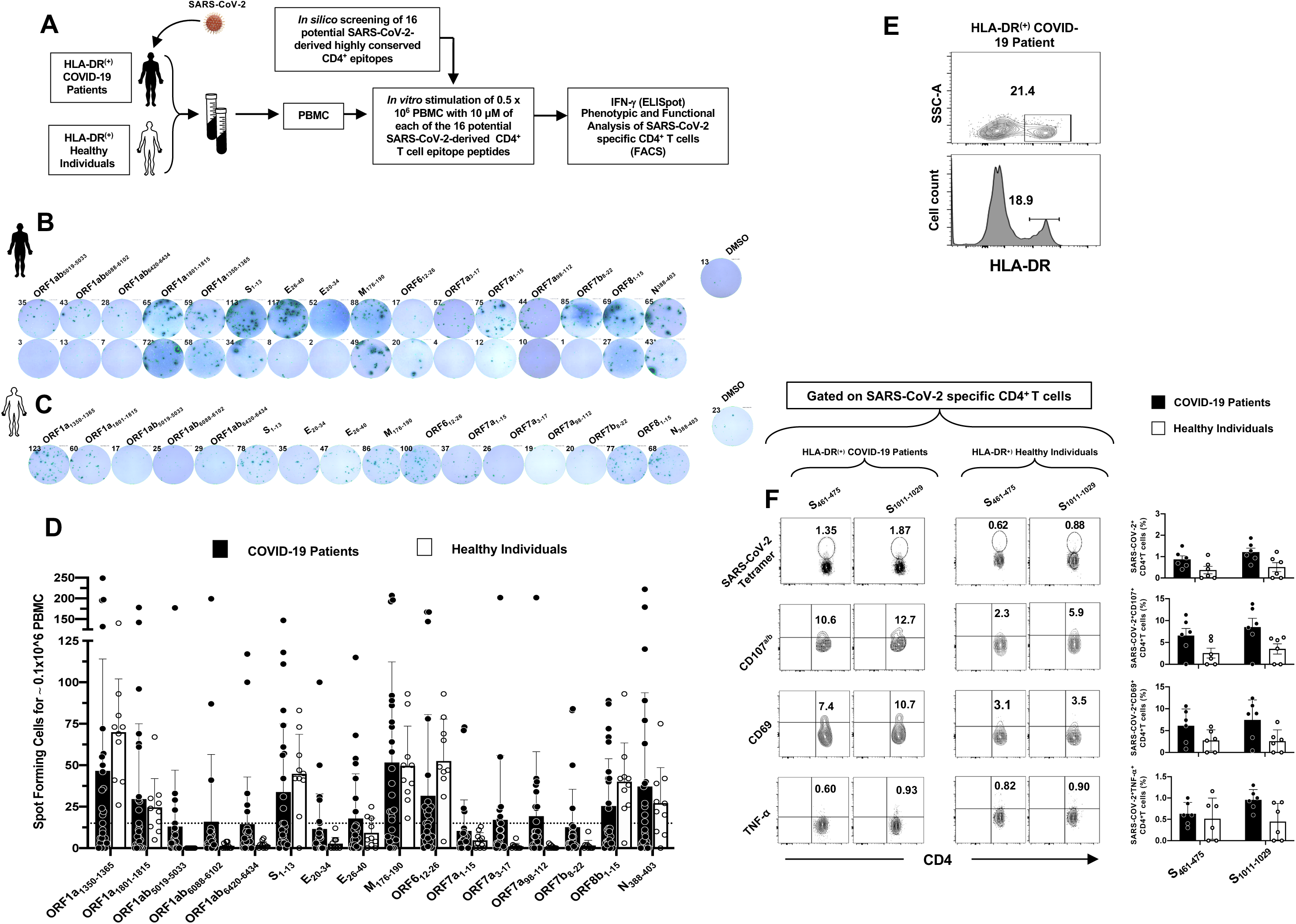
CD4^+^ T cells specific to highly conserved SARS-CoV-2 epitopes detected in COVID-19 patients and unexposed healthy individuals: (**A**) Experimental design: PBMCs from HLA-DR positive COVID-19 patients (*n* = 63) (**B**) and controls unexposed healthy individuals (*n* = 10) (**C**) were isolated and stimulated for 48 hrs. with 10 uM of each of the 24 SARS-CoV-2-derived CD8^+^ T cell epitope peptides. The number of IFN-γ-producing cells were quantified using ELISpot assay (**B, C** and **D**). PBMCs from HLA-DR-positive COVID-19 patients (**E**) were further stimulated for an additional 5 hours in the presence of mAbs specific to CD107^a^ and CD107^b^, and golgi-plug and golgi-stop. Tetramers specific to two Spike epitopes, CD107^a/b^ and CD69 and TNF-α expression were then measured by FACS. Representative FACS plot showing the frequencies of Tetramer^+^CD4^+^ T cells, CD107^a/b+^CD4^+^ T cells, CD69^+^CD4^+^ T cells and TNF-α^+^CD4^+^ T cells following priming with a group of 2 Spike CD4^+^ T cell epitope peptides (**F**, *left panel*). Average frequencies of tetramer^+^CD4^+^ T cells, CD107^a/b+^CD4^+^ T cells, CD69^+^CD4^+^ T cells and TNF-α^+^CD4^+^ T cells (**F**, *right panel*).

Blood-derived PBMCs from COVID-19 patients (*black*, **Fig. 4B**) and healthy individuals (*white,* **Fig. 4C**) were analyzed by ELISpot for frequencies SARS-CoV-2 epitopes-specific IFN-γ-producing CD8^+^ T cells. As shown in **Figs. 4B** and **4D**, significant numbers of SARS-CoV-2 epitopes-specific memory CD8^+^ T cells producing IFN-γ were detected in PBMCs of COVID-19 patients. Out of the 24 highly conserved cross-reactive SARS-CoV-2 CD8^+^ T cell epitopes, significant T cell responses were detected against 15 epitopes derived from structural Spike (S) and Envelope proteins (E) (i.e. S_958-966_, S_976-984_, S_1220-1228_, and E_26-34_) and from non-structural proteins (i.e. ORF1ab_84-92_, ORF1ab_1675-1683_, ORF1ab_2210-2218_, ORF1ab_2363-2371_, ORF1ab_3732-3740_, ORF1ab_4283-4290_, ORF1ab_6419-6427_, ORF1ab_6749-6757_, ORF6_4-11_, ORF10_3-11_ and ORF10_5-13_) (**Figs. 2D, 4B** and **4D**). Moreover, among the 24 SARS-CoV-2 epitopes, 11 epitopes recalled memory CD8^+^ T cells from unexposed healthy individuals (i.e. ORF1ab_1675-1683_, ORF1ab_3732-3740_, ORF1ab_4283-4290_, ORF1ab_6419-6427_, ORF1ab_6749-6757_, S_2-10_, S_958-966_, S_976-984_, S_1220-1228_, ORF10_3-11_, and ORF10_5-13_) (**Figs. 4C** and **Fig. 4D**). However, the unexposed healthy individuals exhibited a different pattern of CD8^+^ T cell immunodominance as compared to COVID-19 patients. We then compared the epitopes-specificity and function of memory CD8^+^ T cells in HLA-*A0201-positive COVID-19 patients and healthy individuals (**Fig. 4E**) using flow cytometry (**Fig. 4F**). For a better comparison, a similar FACS gating strategy was applied to PBMCs-derived T cells from both COVID-19 and healthy donors (*not shown*). The COVID-19 patients appeared to have a higher frequency of CD8^+^ T cells compared to healthy donors (**Fig. 4F**). Tetramer staining showed that many of SARS-CoV-2 epitope-specific CD8^+^ T cells are multifunctional producing IFN-γ, TNF-α and expressing CD69 and CD107^a/b^ markers of activation and cytotoxicity in COVID-19 patients (**Fig. 4F**).

Similar to SARS-CoV-2 memory CD8^+^ T cells, memory CD4^+^ T cells specific to several highly conserved SARS-CoV-2 epitopes were detected in both COVID-19-recovered patients and unexposed healthy individuals (**Figs. 5B-D**). Significant responses were detected in COVID-19 patients against 10 epitopes derived from both structural proteins (i.e. S_1-13_, E_26-40_, M_176-190_ and N_388-403_) and non-structural proteins (i.e. ORF1a_1350-1365_, ORF1a_1801-1815_, ORF6_12-26_, ORF7a_3-17_, ORF7a_98-112_, ORF7b_8-22_ and ORF8b_1-15_) out of the 16 highly conserved cross-reactive SARS-CoV-2 CD4^+^ T cell epitopes (**Figs. 3, 5B** and **5D**). Among the 16 SARS-CoV-2 epitopes, 7 immunodominant epitopes recalled memory CD4^+^ T cells from unexposed healthy individuals with no history of COVID-19 or no contact with COVID-19 patients (i.e. ORF1a_1350-1365_, ORF1a_1801-1815_, S_1-13_, M_176-190_, ORF6_12-26_, ORF8b_8-22_ and N_388-403_) (**Figs. 5C** and **5D**). Unlike for CD8^+^ T cell responses, the unexposed healthy individuals exhibited a similar pattern of CD4^+^ T cell immunodominance as compared to COVID-19 patients. Significant proportions of SARS-CoV-2 epitopes-specific CD4^+^ T cells, expressing CD69, CD107^a/b^ and TNF-α, were detected using specific tetramers in PBMCs of HLA-DR positive COVID-19 patients, as compared to healthy patients (**Figs. 5E** and **5F**).

The immunogenicity of the identified SARS-CoV-2 human CD4^+^ and CD8^+^ T cell epitopes was assessed in “humanized” HLA-DR/HLA-A*02:01 double transgenic mice (**Figs. 6A** and **7A**). A mixture of peptides incorporating CD4^+^ T-cell or CD8^+^ T-cell epitopes were delivered with CpG and Alum, as shown in **Figs. 6A** and **7A** and detailed in the *Materials* and *Methods*. As a negative control, mice received adjuvant alone. The induced SARS-CoV-2 epitope-specific CD4^+^ and CD8^+^ T cell responses were determined in the spleen using multiple immunological assays, including IFN-γ ELISpot, FACS surface markers of activation, markers of cytotoxic degranulation and intracellular cytokine staining. The gating strategy employed in mice is shown in **Figs 6B** and **7B**. Two weeks after the second immunization with the mixture of CD8^+^ T-cell peptides, 11 out of 24 highly conserved SARS-CoV-2 human CD8^+^ T cell epitope peptides were immunogenic in “humanized” HLA-DR/HLA-A*02:01 double transgenic mice (**Figs. 6C** and **6D**). The immunogenic epitopes were derived from both structural Spike protein (S_2-10_, S_958-966_, S_1000-1008_ and S_1220-1228_) and Envelope protein (E_20-28_) and from non-structural proteins (i.e. ORF1ab_2363-2371_, ORF1ab_3732-3740_, ORF1ab_5470-5478_, ORF8_73-81_, and ORF10_5-13_). Moreover, 8 out of 16 SARS-CoV-2 peptides induced significant CD4^+^ T-cell responses in “humanized” HLA-DR/HLA-A*02:01 double transgenic mice (**Figs. 7C** and **7D**). The immunogenic epitopes were derived from both structural Spike protein (S_1-13_) and membrane protein (M_176-190_) and from non-structural proteins (ORF1a_1350-1365_, ORF1a_5019-5033_, ORF1a_6420-6434_, ORF6_12-26_, ORF7b_8-22_ and ORF8b_1-15_).

**Figure 6:**
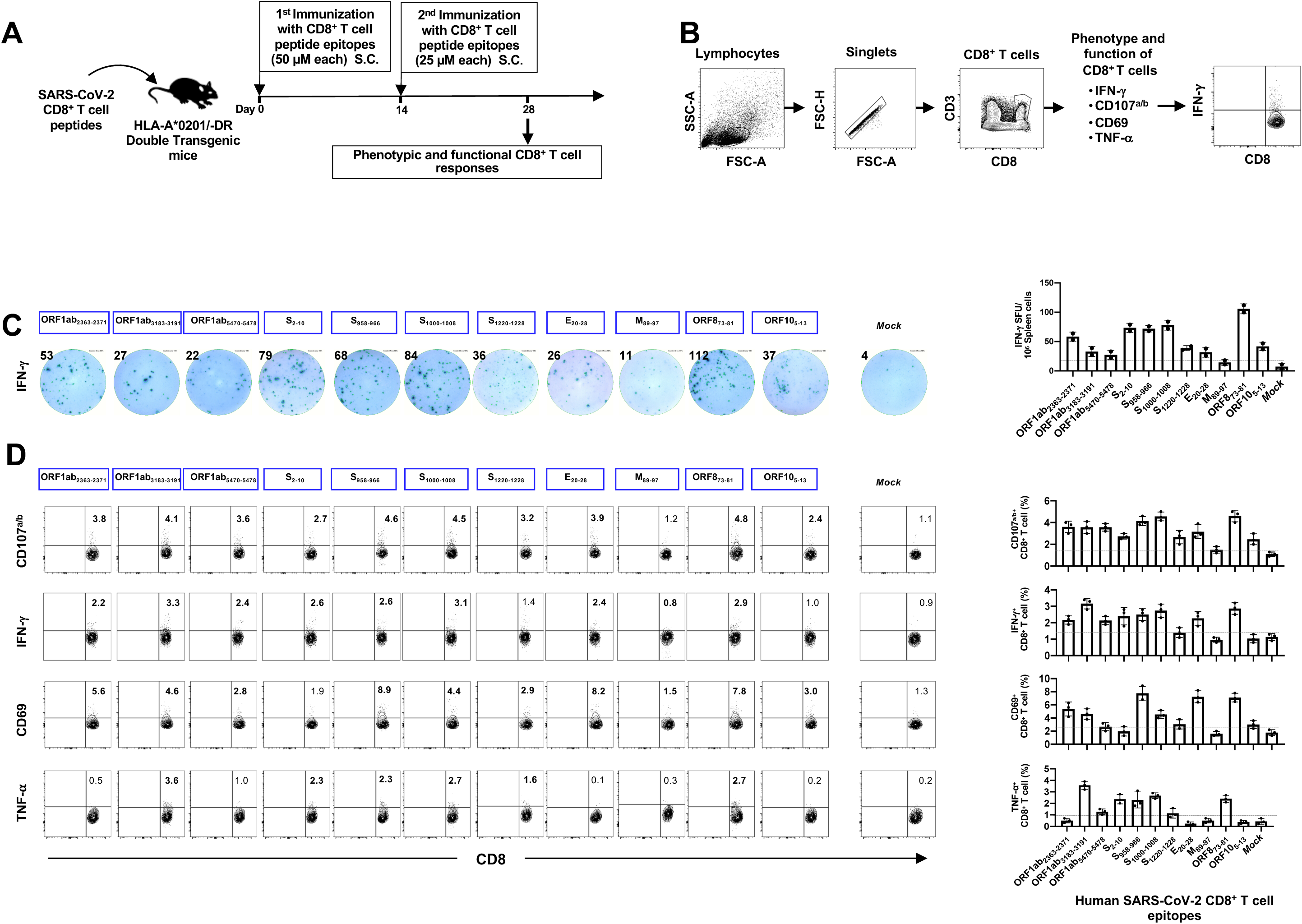
Immunogenicity of genome-wide identified human SARS-CoV-2 CD8^+^ T epitopes in HLA-A*02:01/HLA-DR double transgenic mice. (**A**) Timeline of immunization and immunological analyses. Eight groups of age-matched HLA-A*02:01 transgenic mice (*n* = 3) were immunized subcutaneously, on days 0 and 14, with a mixture of four SARS-CoV-2-derived human CD8^+^ T cell peptide epitopes mixed with PADRE CD4^+^ T helper epitope, delivered in alum and CpG_1826_ adjuvants. As a negative control, mice received adjuvants alone (mock-immunized). (**B**) Gating strategy used to characterize spleen-derived CD8^+^ T cells. Lymphocytes were identified by a low forward scatter (FSC) and low side scatter (SSC) gate. Singlets were selected by plotting forward scatter area (FSC-A) vs. forward scatter height (FSC-H). CD8 positive cells were then gated by the expression of CD8 and CD3 markers. (**C**) Representative ELISpot images (*left panel*) and average frequencies (*right panel*) of IFN-γ-producing cell spots from splenocytes (10^6^ cells/well) stimulated for 48 hours with 10 µM of individual SARS-CoV-2 peptides. The number on the top of each ELISpot image represents the number of IFN-γ-producing spot forming T cells (SFC) per one million splenocytes. (**D**) Representative FACS plot (*left panel*) and average frequencies (*right panel*) of IFN-γ and TNF-α production by, and CD107^a/b^ and CD69 expression on SARS-CoV-2 epitope-primed CD8^+^ T cells determined by FACS. Numbers indicate frequencies of IFN-γ^+^CD8^+^ T cells, CD107^+^CD8^+^ T cells, CD69^+^CD8^+^ T cells and TNF-α^+^CD8^+^ T cells, detected in 3 immunized mice.

**Figure 7:**
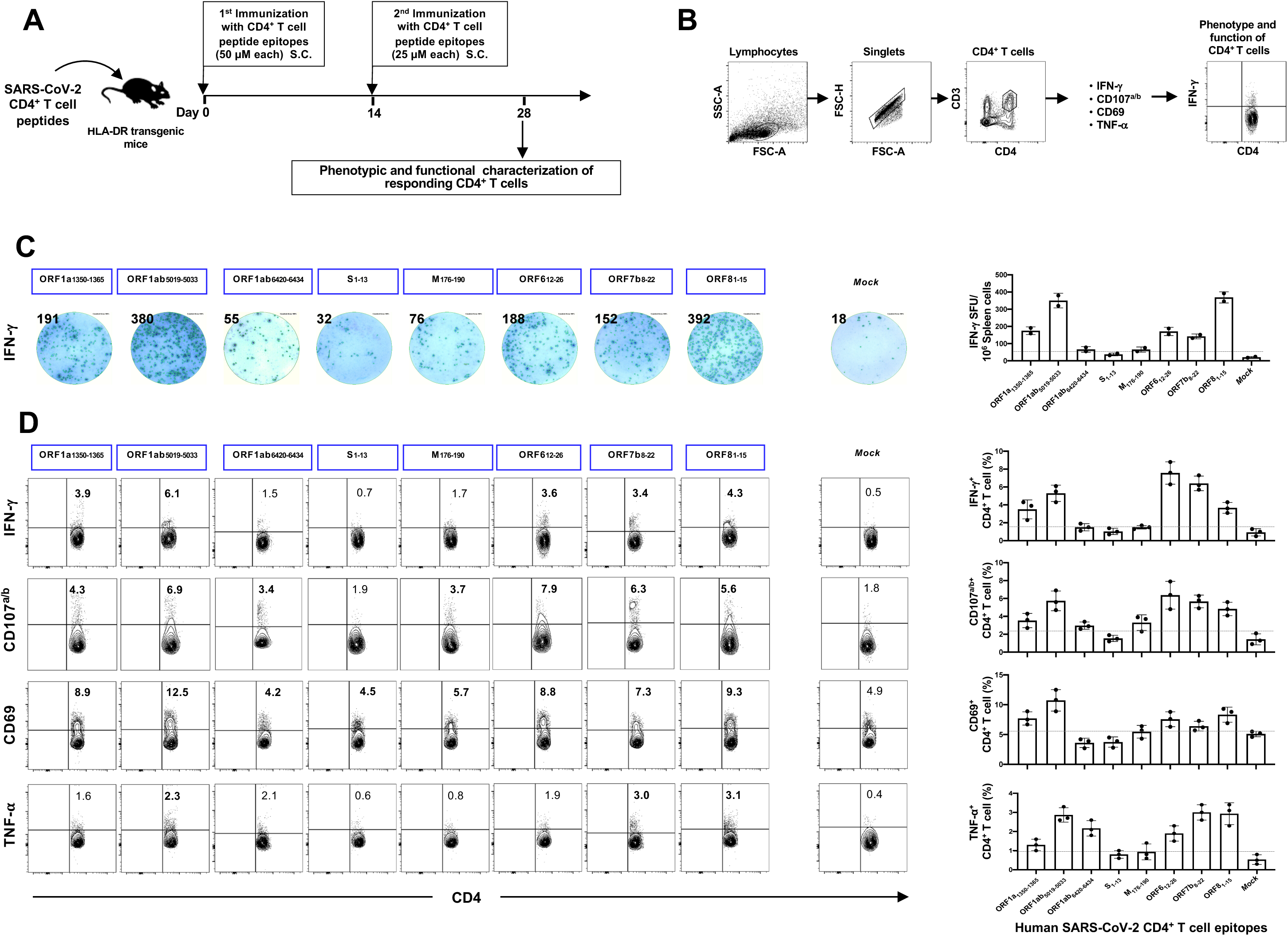
Immunogenicity of genome-wide identified human SARS-CoV-2 CD4^+^ T epitopes in HLA-A*02:01/HLA-DR double transgenic mice. (**A**) Timeline of immunization and immunological analyses. Four groups of age-matched HLA-DR transgenic mice (*n* = 3) were immunized subcutaneously, on days 0 and 14, with a mixture of four SARS-CoV-2-derived human CD4^+^ T cell peptide epitopes delivered in alum and CpG_1826_ adjuvants. As a negative control, mice received adjuvants alone (mock-immunized). (**B**) Gating strategy used to characterize spleen-derived CD4^+^ T cells. CD4 positive cells were gated by the CD4 and CD3 expression markers. (**C**) Representative ELISpot images (*left panel*) and average frequencies (*right panel*) of IFN-γ-producing cell spots from splenocytes (10^6^ cells/well) stimulated for 48 hours with 10 µM of individual SARS-CoV-2 peptides. The number of IFN-γ-producing spot forming T cells (SFC) per one million of total cells is presented on the top of each ELISpot image. (**D**) Representative FACS plot (*left panel*) and average frequencies (*right panel*) show IFN-γ and TNF-α-production by, and CD107^a/b^ and CD69 expression on SARS-CoV-2 epitope-primed CD4^+^ T cells determined by FACS. The numbers indicate percentages of IFN-γ^+^CD4^+^ T cells, CD107^+^CD4^+^ T cells, CD69^+^CD4^+^ T cells and TNF-α^+^CD4^+^ T cells detected in 3 immunized mice.

Altogether, these results indicate that pre-existing memory CD4^+^ T and CD8^+^ T cells specific to both structural and non-structural protein antigens and epitopes are present in COVID-19 patients and unexposed healthy individuals. Most T cell epitopes are concentrated in the non-structural proteins which appeared to be more targets for human CD4^+^ T and CD8^+^ T cells. These memory T cells recognized highly conserved SARS-CoV-2 epitopes that cross-react with the human and animal Coronaviruses. It is likely that infection with “common cold” Coronavirus induced long-lasting memory CD4^+^ and CD8^+^ T cells specific to the structural and non-structural SARS-CoV-2 epitopes in healthy unexposed individuals. Heterologous immunity and heterologous immunopathology orchestrated by these cross-reactive epitope-specific memory CD4^+^ and CD8^+^ T cells, following previous multiple exposures to “common cold” Coronaviruses, may have shaped protection vs. susceptibility to SARS-CoV-2 infection and disease, with a yet-to-be determined mechanism.

### 5. Identification of B-cell epitopes from SARS-CoV-2 Spike protein that are highly conserved between human and animal Coronaviruses, that are antigenic in humans and immunogenic in “humanized” HLA transgenic mice

We next predicted potential linear B-cell (antibody) epitopes on Spike protein sequence of the first SARS-CoV-2-Wuhan-Hu-1 strain (NCBI GeneBank accession number MN908947.3) using BepiPred 2.0, with a cutoff of 0.55 (corresponding to a specificity greater than 0.81 and sensitivity below 0.30) and considering sequences having more than 5 amino acid residues ^54^. This stringent screening process initially resulted in the identification of 28 linear B-cell epitopes (**Supplemental Table S3**). From this pool of 28 potential epitopes, we later selected 10 B-cell epitopes, (19 to 62 amino acids in length), based on: (*i*) their sequences being highly conserved between SARS-CoV-2, the main 4 major ‘‘common cold’’ Coronaviruses (CoV-OC43, CoV-229E, CoV-HKU1, and CoV-NL63 ^55^), and the SARS-like SL-CoVs that are isolated from bats, civet cats, pangolins and camels; and (*ii*) the probability of exposure each linear epitope to the surface of infected target cells (**Fig. 8**). The Spike epitope sequences highlighted in blue indicate a high degree of homology among the currently circulating 81,963 SARS-CoV-2 strains and at least a 50% conservancy among two or more human SARS-CoV strains from previous outbreaks, and the SL-CoV strains isolated from bats, civet cats, pangolins and camels (**Fig. 8**). Two of the 10 B-cell epitopes namely S_369-393_, and S_440-501_ overlap with the Spike’s receptor binding domain (RBD) region that bind to the ACE2 receptor (designated as RBD-1 and RBD-2 in **Supplemental Fig. S4A**). Higher interaction similarity scores were observed for RBD-derived epitopes S_369-393_ and S_471-501_ when molecular docking was performed against the ACE2 receptor (**Supplemental Fig. S4B**).

**Figure 8:**
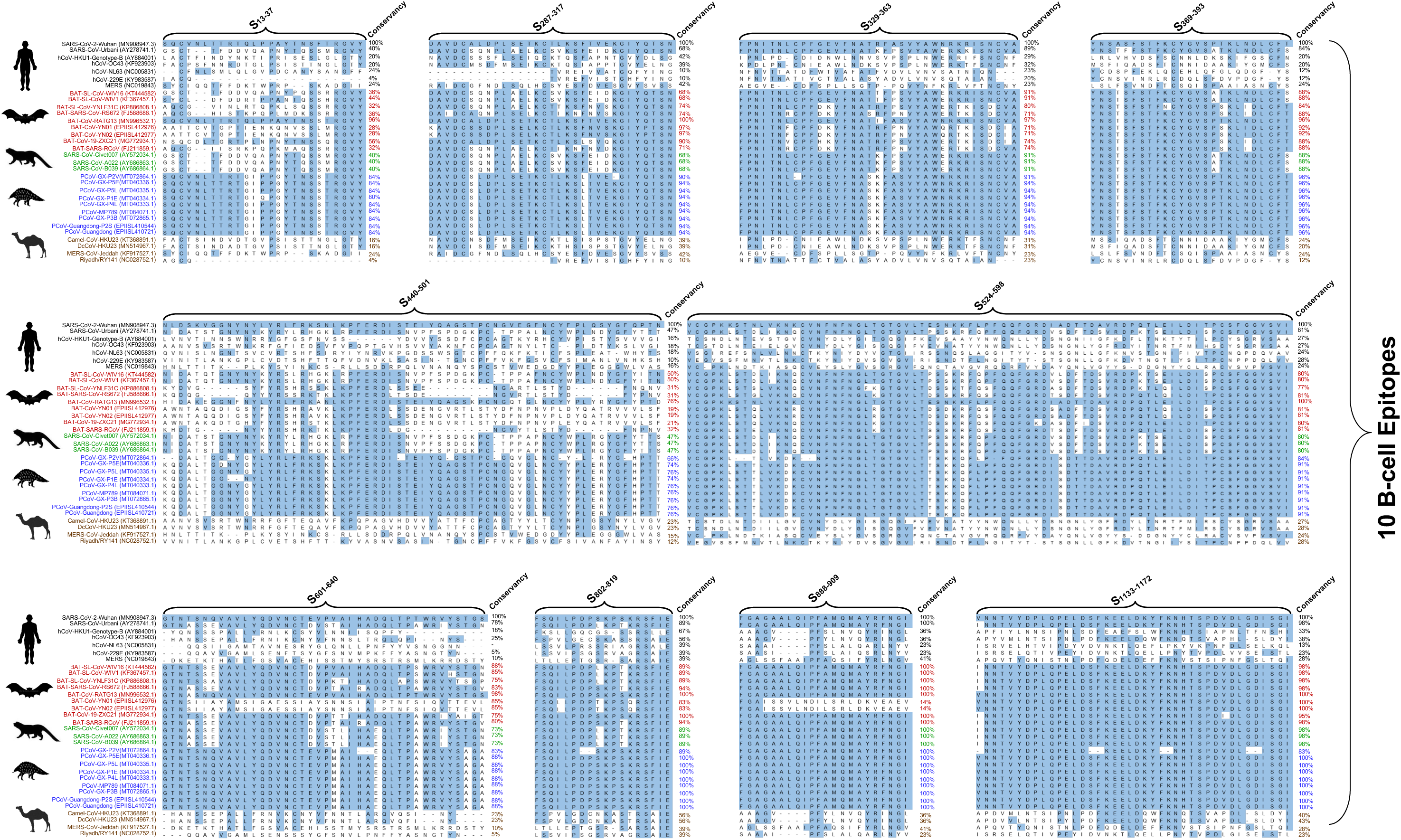
Conservation of Spike-derived B cell epitopes among human, bat, civet cat, pangolin, and camel coronavirus strains: Multiple sequence alignment performed using ClustalW among 29 strains of SARS coronavirus (SARS-CoV) obtained from human, bat, civet, pangolin, and camel. This includes 7 human SARS/MERS-CoV strains (SARS-CoV-2-Wuhan (MN908947.3), SARS-HCoV-Urbani (AY278741.1), CoV-HKU1-Genotype-B (AY884001), CoV-OC43 (KF923903), CoV-NL63 (NC005831), CoV-229E (KY983587), MERS (NC019843)); 8 bat SARS-CoV strains (BAT-SL-CoV-WIV16 (KT444582), BAT-SL-CoV-WIV1 (KF367457.1), BAT-SL-CoV-YNLF31C (KP886808.1), BAT-SARS-CoV-RS672 (FJ588686.1), BAT-CoV-RATG13 (MN996532.1), BAT-CoV-YN01 (EPIISL412976), BAT-CoV-YN02 (EPIISL412977), BAT-CoV-19-ZXC21 (MG772934.1); 3 Civet SARS-CoV strains (SARS-CoV-Civet007 (AY572034.1), SARS-CoV-A022 (AY686863.1), SARS-CoV-B039 (AY686864.1)); 9 pangolin SARS-CoV strains (PCoV-GX-P2V(MT072864.1), PCoV-GX-P5E(MT040336.1), PCoV-GX-P5L (MT040335.1), PCoV-GX-P1E (MT040334.1), PCoV-GX-P4L (MT040333.1), PCoV-MP789 (MT084071.1), PCoV-GX-P3B (MT072865.1), PCoV-Guangdong-P2S (EPIISL410544), PCoV-Guangdong (EPIISL410721)); 4 camel SARS-CoV strains (Camel-CoV-HKU23 (KT368891.1), DcCoV-HKU23 (MN514967.1), MERS-CoV-Jeddah (KF917527.1), Riyadh/RY141 (NC028752.1)) and 1 recombinant strain (FJ211859.1)). Regions highlighted with blue color represent the sequence homology. The B cell epitopes, which showed at least 50% conservancy among two or more strains of the SARS Coronavirus or possess receptor-binding domain (RBD) specific amino acids were selected as candidate epitopes.

We later determined the ability of each of the 10 B-cell epitopes selected from the Spike protein, that showed a high conservancy between human and animal Coronaviruses, to induce in B6 mice SARS-CoV-2 epitope-specific antibody-producing plasma B cells and IgG antibodies (**Fig. 9**). Synthetic peptides corresponding to each linear B cell epitope were produced. Since 4 epitopes were too long to synthetize (e.g. 62 amino acids), they were divided into 2 or 3 short fragments resulting in a total of 17 B-cell epitope peptides (**Supplemental Table S3**). As illustrated in **Fig. 9A**, groups of five B6 mice each received two subcutaneous (s.c.) injections with mixtures of 3 to 4 B-cell epitope peptides, mixed with CpG and Alum adjuvants. Negative control mice received adjuvant alone, without Ags. The frequency of antibody-producing plasma B cells and the level of IgG antibodies specific to each SARS-CoV-2 B cell epitope were determined in the spleen and in the serum using FACS staining of CD138 and B220 surface markers and IgG-ELISpot and ELISA assays, respectively. The gating strategy used to determine the frequencies of plasma B-cells in the spleen is shown in **Fig. 9B**. Out of the 17 Spike B-cell epitopes, 7 epitopes (S_13-37_, S_287-317_, S_524-558_, S_544-578_, S_565-598_, S_601-628_, and S_614-640_) induced high frequencies of CD138^+^B220^+^ plasma B cells in the spleen of B6 mice (**Fig. 9C**). The IgG ELISpot assay confirmed that 7 out of the 17 Spike B-cell epitopes induced significant numbers of IgG-producing B cells in the spleen (**Fig. 9D**). Moreover, significant amount of IgG were detected in the serum of the immunized B6 mice. These IgG antibodies were specific to 5 out of the 17 Spike B-cell peptide epitopes (S_13-37_, S_287-317_, S_565-598_, S_601-628_, and S_614-640_) (**Fig. 9E**). As expected non-immunized animals or those that received adjuvant alone did not develop detectable IgG responses. Of these 5 highly immunogenic B cell peptides, 4 peptides (S_13-37_, S_287-317_, S_601-628_, and S_614-640_) were highly antigenic as they were recognized by serum IgG from COVID-19 patients, confirming the presence of at least one native linear B cell epitope in each peptide (**Fig. 9F**). In summary, we identified four highly conserved immunogenic and antigenic human B-cell target epitopes from the Spike SARS-CoV-2 virus that recall IgG antibodies from COVID-19 patients.

**Figure 9:**
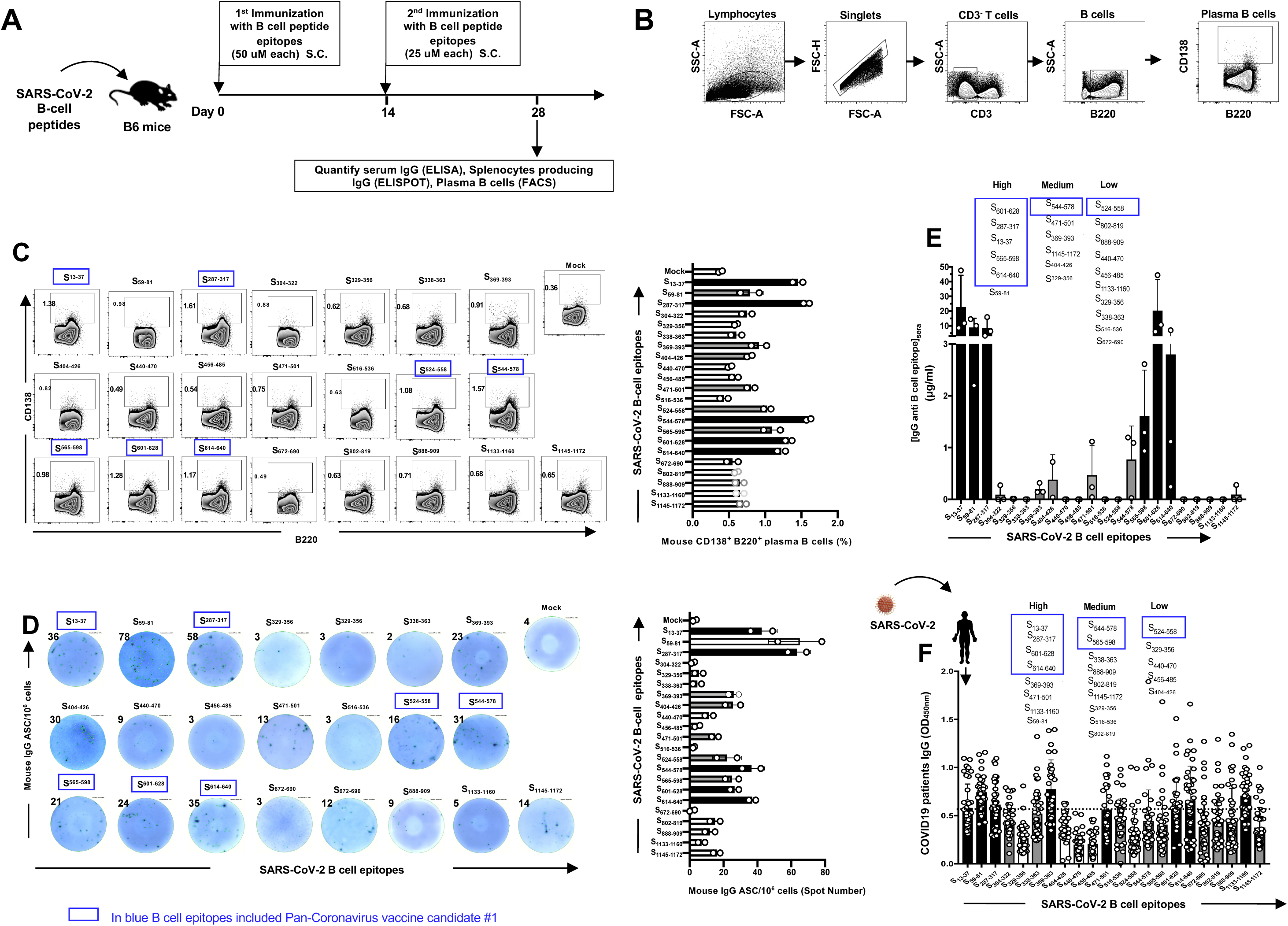
IgG antibodies specific to SARS-CoV-2 Spike protein-derived B-cell epitopes in immunized B6 mice and in convalescent COVID-19 patients: (**A**) Timeline of immunization and immunological analyses. A total of 17 SARS-CoV-2 derived B-cell epitope peptides selected from SARS-CoV-2 Spike protein and tested in B6 mice were able to induce antibody responses. Four groups of age-matched B6 mice (*n* = 3) were immunized subcutaneously, on days 0 and 14, with a mixture of 4 or 5 SARS-CoV-2 derived B-cell peptide epitopes emulsified in alum and CpG_1826_ adjuvants. Alum/CpG_1826_ adjuvants alone were used as negative controls (mock-immunized). (**B** and **C**) The frequencies of IgG-producing CD3^(-)^CD138^(+)^B220^(+)^ plasma B cells were determined in the spleen of immunized mice by flow cytometry. (**B**) The gating strategy was as follows: Lymphocytes were identified by a low forward scatter (FSC) and low side scatter (SSC) gate. Singlets were selected by plotting forward scatter area (FSC-A) versus forward scatter height (FSC-H). B cells were then gated by the expression of CD3^(-)^ and B220^(+)^ cells and CD138 expression on plasma B cells determined. (**C**) Representative FACS plot (*left panels*) and average frequencies (*right panel*) of plasma B cells detected in spleen of immunized mice. The percentages of plasma CD138^(-)^B220^(+)^B cells is indicated in the top left of each dot plot. (**D**) SARS-CoV-2 derived B-cell epitopes-specific IgG responses were quantified in immune serum, 14 days post-second immunization (i.e. day 28), by ELISpot (Number of IgG^(+)^Spots). Representative ELISpot images (*left panels*) and average frequencies (*right panel*) of anti-peptide specific IgG-producing B cell spots (1 x10^6^ splenocytes/well) following 4 days *in vitro* B cell polyclonal stimulation with mouse Poly-S (Immunospot). The top/left of each ELISpot image shows the number of IgG-producing B cells per half a million cells. ELISA plates were coated with each individual immunizing peptide. The B-cell epitopes-specific IgG concentrations (µg/mL) measured by ELISA in: (**E**) Levels of IgG detected in peptide-immunized B6 mice, after subtraction of the background measured from mock-vaccinated mice. The dashed horizontal line indicates the limit of detection; and in (**F**) Level of IgG specific to each of the 17 Spike peptides detected SARS-CoV-2 infected patients, after subtraction of the background measured from healthy non-exposed individuals. *Black bars* and *gray bars* show high and medium immunogenic B cell peptides, respectively. The dashed horizontal line indicates the limit of detection.

### 6. Identification of genome-wide highly conserved SARS-CoV-2 human B, CD4^+^ and CD8^+^ T cell epitopes that are associated with asymptomatic SARS-CoV-2 infection

We next investigated whether antibodies, CD4^+^ and CD8^+^ T cells from symptomatic vs. asymptomatic COVID-19 patients and unexposed healthy individuals had different profile of epitopes specificities.

The COVID-19 patients were divided into 4 groups depending on the severity of the symptoms: Group 1 that comprised SARS-CoV-2 infected patients that never develop any symptoms (i.e. asymptomatic patients, *n* = 11); Group 2 with mild symptoms (i.e. Inpatient only, *n* = 32); Group 3 with moderate symptoms (i.e. ICU admission, *n* = 11) and Group 4 with severe symptoms (i.e. ICU admission +/− Intubation or death, *n* = 9). As expected, compared to the asymptomatic group, all of the 3 symptomatic groups (i.e. mild, moderate and severe) had higher percentages of comorbidities, including diabetes (22% to 64%), hypertension (64% to 78%), cardiovascular disease (11% to 18%) and obesity (9% to 50%) (**Table 1**). The final Group 5 comprised of unexposed healthy individuals (controls), with no history of COVID-19 or contact with COVID-19 patients (*n* = 10).

The COVID-19 patients that present severe symptoms had significantly higher frequencies of ORF1ab_1675-1683_ and ORF1ab_2210-2218_ epitopes-specific CD8^+^ T cells compared to asymptomatic patients (**Fig. 10**). In contrast, a significantly higher frequencies of S_691-699_, ORF6_4-11_, and ORF8a_31-39_ epitope-specific CD8^+^ T cells were detected in asymptomatic patients and unexposed healthy individuals as compared to COVID-19 patients with moderate to severe symptoms (**Fig. 10A**). However, there were no significant correlations between the overall frequencies of SARS-CoV-2-specific CD8^+^ T cells and the severity of the COVID-19 disease (**Fig. 10B**, *P* = 0.6088).

**Figure 10:**
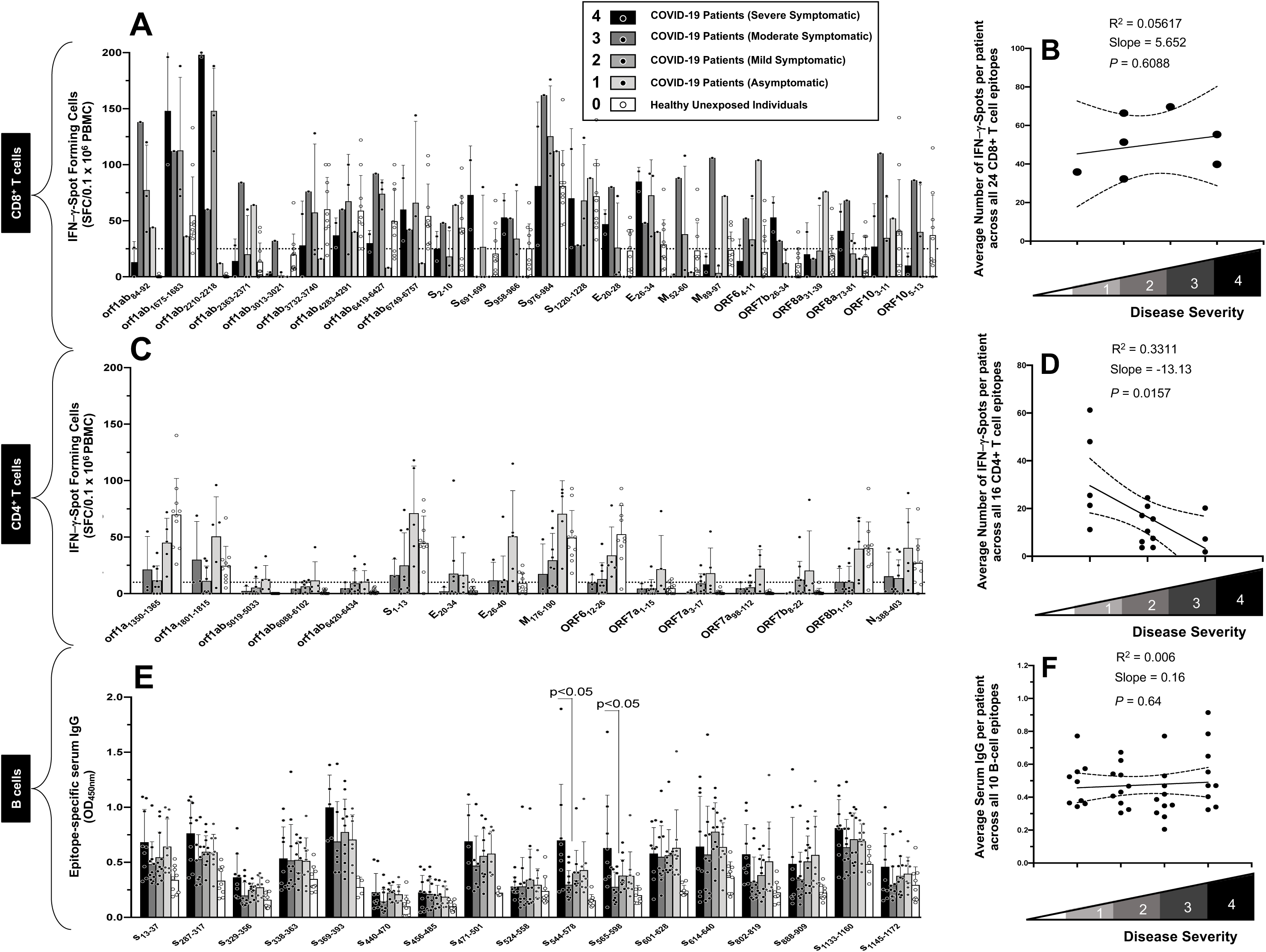
Association of immune responses specific to SARS-CoV-2 B, CD4^+^ and CD8^+^ T cell epitopes with severity of COVID-19 disease. Serum and PBMCs from COVID-19 patients (*n* = 62) and controls unexposed healthy individuals (*n* = 10) were isolated and stimulated overnight with 10 uM of each of the 24 SARS-CoV-2-derived CD8^+^ T cell epitope peptides. The number of IFN-γ-producing cells were quantified using ELISpot assay and Spike epitopes-specific IgG antibodies detected by ELISA. (**A**) Average frequencies of IFN-γ-producing CD8^+^ T cells in COVID-19 patients, with various levels of severity and from unexposed healthy individuals. (**B**) Correlations between the overall frequencies of SARS-CoV-2-specific CD8^+^ T cells and COVID-19 disease severity. (**C**) Average frequencies of IFN-γ-producing CD4^+^ T cells in COVID-19 patients, with various levels 0f severity and from unexposed healthy individuals. (**D**) Correlations between the overall frequencies of SARS-CoV-2-specific CD4^+^ T cells and COVID-19 disease severity. (**E**) Average level of IgG specific to each of the 17 Spike peptides detected SARS-CoV-2 infected patients, after subtraction of the background measured from healthy non-exposed individuals. (**F**) Correlations between the IgG responses detected in COVID-19 patients, with various levels of disease severity.

In contrast to CD8^+^ T cells, we detected a significantly higher frequencies of memory CD4^+^ T cells specific to 8 highly conserved epitopes in asymptomatic COVID-19 patients compared to symptomatic COVID-19 patients, regardless of level of severity of symptoms (**Fig. 10C**, *P* < 0.005). Moreover, significant correlations were detected between the overall high frequencies of SARS-CoV-2-specific CD4^+^ T cells and low severity of symptoms in COVID-19 patients, suggesting a critical role of CD4^+^ T cells specific to highly conserved asymptomatic epitopes in the apparent “protection” from COVID-19 symptoms (**Fig. 10D**, *P* = 0.0157). Finally, we found the symptomatic patients with the most severe symptoms had significantly higher IgG antibodies specific to some linear B-cell epitopes (e.g. S_544-578_ and S_565-598_) compared to asymptomatic patients, suggesting some Spike-specific antibodies may not be protective, but instead maybe immune enhancing (**Fig. 10E**, *P* < 0.05). No significant correlations was found between the overall Spike-specific antibody responses and the low severity of symptoms in COVID-19 patients (**Fig. 10F**, *P* > 0.05).

Circulating memory CD4^+^ and CD8^+^ T_CIRC_ cell populations are categorized into two major phenotypically distinct sub-populations: effector memory T cells (CD62L^low^CD44^high^T_EM_) and central memory T cells (CD62L^high^CD44^high^T_CM_) ^56, 57^. Next, we investigated whether symptomatic vs. asymptomatic COVID-19 patients had different frequencies of SARS-CoV-2-specific memory CD4^+^ and CD8^+^ T_EM_ and T_CM_ sub-populations (**Fig. 11A**). We gated on CD3^+^CD4^+^ T cells and CD3^+^CD8^+^ T cells specific to the immunodominant SARS-CoV-2 epitopes (**Fig. 11B**). As shown in **Figs. 11C** and **D**, regardless of the severity of the symptoms, COVID-19 patients had significantly higher frequencies of CD62L^low^CD44^high^CD8^+^ T_EM_ cells compared to the frequencies of CD62L^high^CD44^high^CD4^+^ T_CM_ cells (*P* < 0.05).

**Figure 11.**
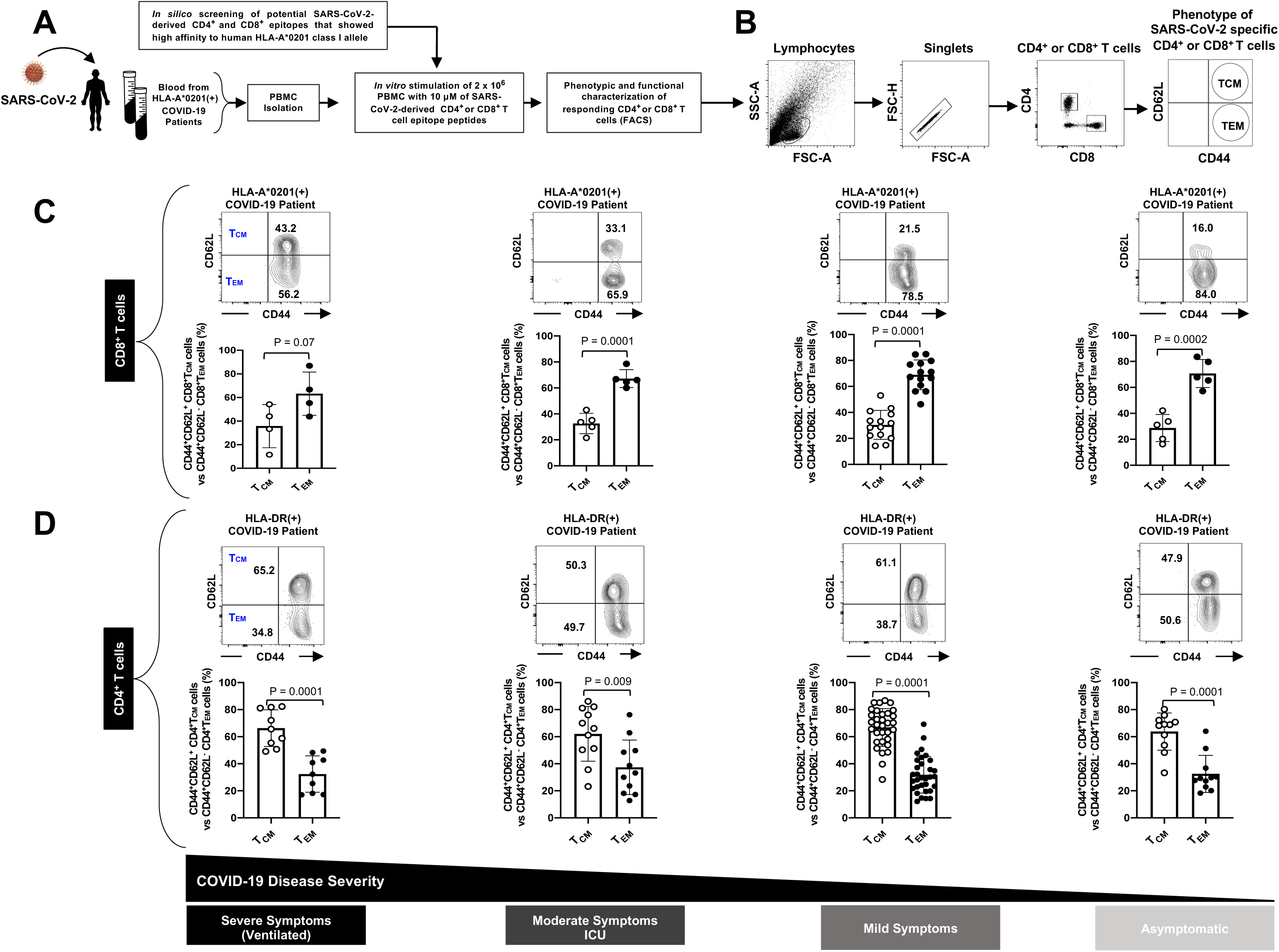
Frequency of CD4^+^ and CD8^+^ T cells with effector memory phenotype (CD8^+^ T_EM_ cells) detected in symptomatic and asymptomatic COVID-19 patients. (**A**) PBMCs from COVID-19 patients (*n* = 63) were isolated. (**B**) The gating strategy used to phenotype CD4^+^ and CD8^+^ T cells in COVID-19 patients. The frequencies of CD8^+^ (**C**) and CD4^+^ (**D**) T cell sub-populations with either effector memory T cells (CD62L^low^CD44^high^T_EM_ cells) and central memory T cells (CD62L^high^CD44^high^T_CM_ cells) detected in COVID-19 patients that were divided into 4 groups depending on the severity of the symptoms: Group 1 that comprised SARS-CoV-2 infected patients that never develop any symptoms or any viral diseases (i.e. asymptomatic patients) (*n* = 11); Group 2 with mild symptoms (i.e. Inpatient only, *n* = 32); Group 3 with moderate symptoms (i.e. ICU admission, *n* = 11); and Group 4 with severe symptoms (i.e. ICU admission +/− Intubation or death, *n* = 9). The results are representative of 2 independent experiments in each individual. The indicated *P* values, calculated using one-way ANOVA Test, show statistical significance between symptomatic and asymptomatic COVID patients.

Altogether, these results suggest a dichotomy in the phenotype of memory CD4^+^ and CD8^+^ T cell sub-populations in COVID-19 patients. More of central memory CD4^+^ T_CM_ cells and more of effector memory CD8^+^ T cells appeared present in the blood of COVID-19 patients, regardless of the severity of symptoms. Moreover, we discovered that, in contrast to SARS-CoV-2 epitopes-specific CD8^+^ T cells and IgG antibodies, high frequencies of IFN-γ-producing CD4^+^ T cells specific to 8 highly conserved Coronavirus epitopes, were associated with asymptomatic SARS-CoV-2 infection. This suggests that the asymptomatic COVID-19 patients that develop high frequencies of functional IFN-γ-producing CD4^+^ T cells specific to cross-reactive Coronavirus epitopes from structural and non-structural proteins (**Fig. 12**), may have been better protected against subsequent severe SARS-CoV-2 infection and/or disease.

**Figure 12:**
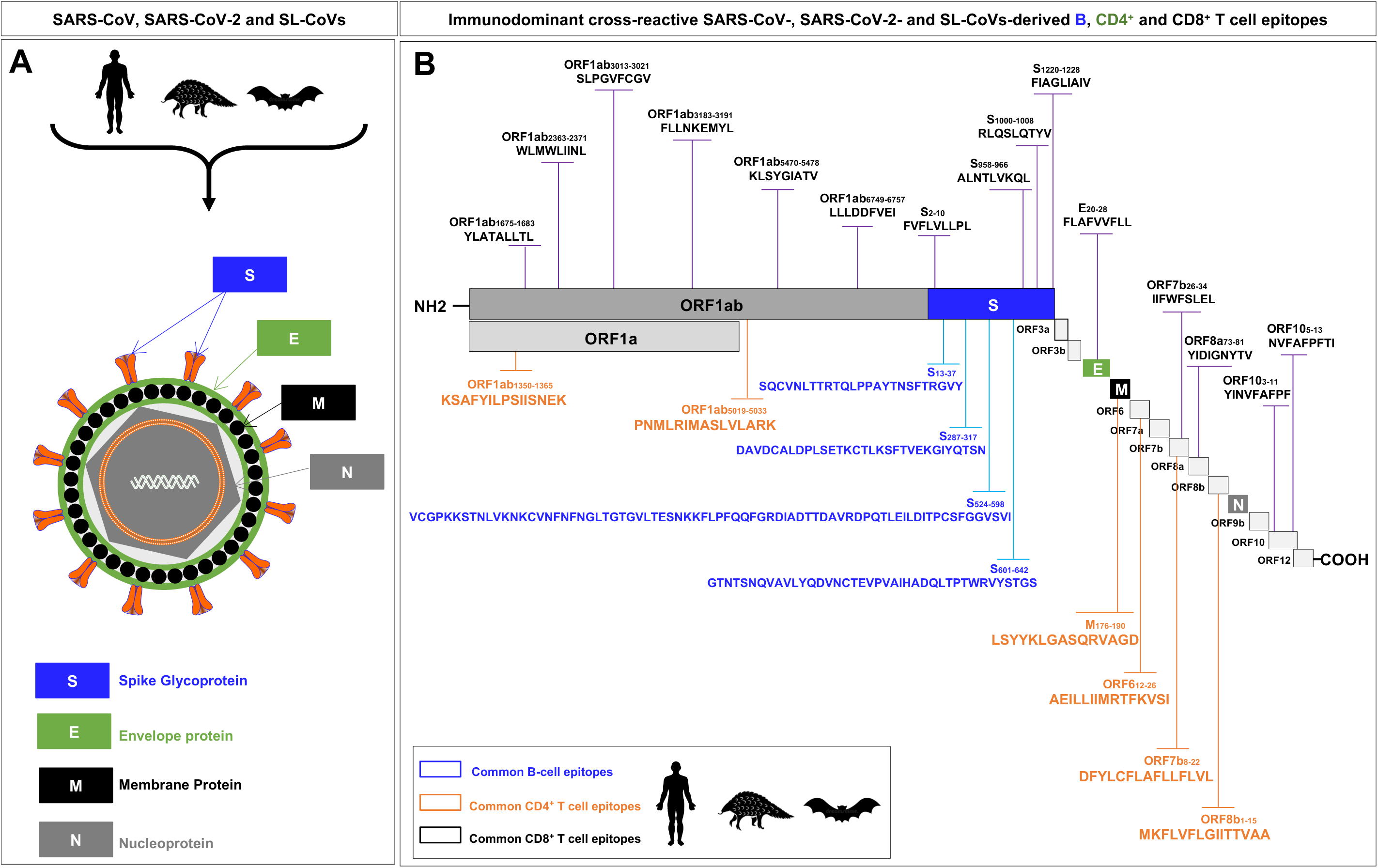
Illustrations of the SARS-CoV/SARS-CoV-2 genome-wide location of the highly conserved, antigenic and immunogenic CD4^+^ T cell, CD8^+^ T cell, and B-cell epitopes. (**A**) Enveloped, spherical, about 120 nm in diameter, SARS-CoV/SARS-CoV-2 genome encodes four structural proteins: spike (S), envelope (E), membrane (M), and nucleocapsid (N), highlighted in blue, green, gray and black, respectively. (**B**) The SARS-CoV/SARS-CoV-2 genome encodes two large non-structural genes ORF1a (green) and ORF1b (gray), encoding 16 non-structural proteins (NSP1–NSP16). The genome encodes at least six accessory proteins (shades of light grey) that are unique to SARS-CoV/SARS-CoV-2 in terms of number, genomic organization, sequence, and function. The common SARS-CoV, SARS-CoV-2 and SL-CoVs-derived human B (*blue*), CD4^+^ (*green*) and CD8^+^ (*black*) T cell epitopes are shown. Structural and non-structural open reading frames utilized in this study were from SARS-CoV-2-Wuhan-Hu-1 strain (NCBI accession number NC-045512.2). The amino acid sequence of the SARS-CoV-2-Wuhan-Hu-1 structural and non-structural proteins was screened for human B, CD4^+^ and CD8^+^ T cell epitopes using different computational algorithms as previously described in *Materials and Methods*. Shown are genome-wide identified SARS-CoV-2 human B cell epitopes (*in blue*), CD4^+^ T cell epitopes (*in green*), CD8^+^ T cell epitopes (*in black*) that are highly conserved between human and animal Coronaviruses.

## DISCUSSION

While the current COVID-19 pandemic will likely disappear through implementation of physical distancing, barriers and mass immunization with a COVID-19 vaccine, it is indispensable that a safe and effective pre-emptive vaccine be developed, and in place ready to protect, against another inevitable COVID pandemic that will emerge in the years to come. More than 169 vaccine candidates against COVID-19 are currently being pre-clinically developed globally ^58^. Many of these vaccines are targeting exclusively the SARS-CoV-2 that causes COVID-19 but no other human and animal Coronaviruses (reviewed in ^58^ and ^59^). Among these, eighteen COVID-19 vaccine candidates, using a variety of different approaches - live attenuated virus, viral vectors and sub-unit mRNA or protein/adjuvant – have already entered Phase I/II/III clinical trials ^59^. While the current vaccine efforts, have focused primarily on COVID-19 and mainly used the Spike protein as target antigen, our strategy has been to develop a pre-emptive pan-Coronavirus vaccine, that incorporates several SARS-CoV-2 epitopes, and designed to target past, present and future Coronavirus outbreaks.

Towards the goal of developing a multi-epitopes pre-emptive pan-Coronavirus human vaccine; in the present study; we identified the asymptomatic and cross-reactive human B and T cell epitopes of SARS-CoV-2 that are highly conserved among the human SARS-CoVs and animal SL-CoVs. While antiviral SARS-CoV-2-specific antibodies and CD4^+^ and CD8^+^ T cell responses appear crucial in protecting asymptomatic COVID-19 patients and convalescent patients, very little information exists with regards to the repertoire of targeted SARS-CoV-2 B and T cell epitopes that are common within a substantial group of human and animal Coronaviruses ^45, 60^. The highly conserved human B and T cell epitopes reported in this study have huge implications for the development of an universal pre-emptive pan-Coronavirus vaccine to induce (or to boost) neutralizing antibodies (Abs), CD4^+^ T helpers (Th1), and antiviral CD8^+^ cytotoxic T-cells (CTL) ^21, 61^. Some of our identified epitopes are similar to those recently reported by Grifoni, *et al*, ^45, 60^ while other epitopes have never been reported. Moreover, in agreement with recent reports ^45, 60, 62^, our study revealed a high degree of similarity among SARS-CoV-2, SARS-CoV and bat-SL-CoVs epitope sequences, but not the MERS-CoV epitope sequences

With no universal pan-Coronavirus vaccine is yet available, five short- and long-term COVID-19 scenarios may occur: (1) the COVID-19 pandemic will be controlled and disappear, with through implementation of physical distancing/barriers or yet-to-be determined mechanism(s) and factor(s). Nevertheless, because small outbreaks SARS-CoV-2 virus will still likely occur around the world, maintaining these small outbreaks in check will still require the implementation of preventive Coronavirus vaccination; (2) COVID-19 pandemic will be controlled by current measures, but there will be localized clusters that may occur in some regions and/or countries of the world, which may lead to another pandemic.; (3) A gradual spread of asymptomatic SARS-CoV-2 infections, that are undetectable, hence may later lead to a resumption of another COVID pandemic; (4) This COVID-19 outbreak returns and transforms into a seasonal “flu-like” global pandemic; and (5) Other global COVID-like pandemics will emerge in the coming years, caused by yet another spillover of an unknown zoonotic bat-derived SARS-like Coronavirus (SL-CoV) into an unvaccinated human population. In all five scenarios; a pre-emptive pan-Coronavirus vaccine that incorporates highly conserved and common epitopes could target not only past and current COVID outbreaks, but may also target future Coronavirus outbreaks that may be caused by yet another spillover of bat SL-CoVs into humans ^21, 61^.

Unlike most protein/adjuvant or vector-based Coronavirus subunit vaccines, that use either whole proteins or whole DNA/mRNA, and whole attenuated or weakened viruses ^63–65^ our multi-epitope Coronavirus vaccine (e.g. the pan-Coronavirus candidate #1 illustrated in **Supplemental Fig. S5**). incorporates multiple human asymptomatic B and CD4^+^/CD8^+^ T cell epitopes that are selected carefully from the whole genome of SARS-CoV-2 for being recognized by antibodies and CD4^+^/CD8^+^ T cells from asymptomatic and convalescent patients that are “naturally protected” from COVID-19. This “asymptomatic” vaccine excludes potentially harmful “symptomatic” epitopes that are highly recognized by antibodies and CD4^+^/CD8^+^ T cells from symptomatic and hence would otherwise increase viral load or exacerbate COVID-19 disease ^50–52^. The present study employed a combinatorial approach for designing an all-in-one multi-epitope pan-Coronavirus vaccine candidate (**Supplemental Fig. S5**) by applying highly conserved genome-wide human B- and T-cell epitopes from 12 genome derived antigenic proteins of SARS-CoV-2. While this study focused on HLA-A*02:01-restricted epitopes, an HLA class I haplotype that is represented similar to 50% of the human population, when epitopes restricted to other HLA-A, HLA-B and HLA-C haplotype including the forecasted population coverage of chosen T cell epitope ensemble (combined HLA class I) is expected to cover 99.8% of the worldwide human population regardless of race and ethnicity. In addition, for a wider vaccine coverage (i.e., close to 99%), our multi-epitope pan-Coronavirus vaccine platform would be easily adapted to include CD8^+^ T cell epitopes for other HLA supertypes that are distributed in the various human populations. The polymorphic HLA molecules can be clustered into a handful of HLA-A supertypes that bind largely overlapping peptide repertoires ^66^. Moreover, such a multi-epitope vaccine would be easily adapted to exclude undesirable epitopes that are restricted to HLA-B*44 and HLA-C*01 alleles which appear to correlate with SARS-CoV-2 virus spreading across certain countries ^67^ and HLA-B*35 allele which appear to be associated with severe pneumonia developed by SARS-CoV-2 in young patients ^68^.

Control of the SARS-CoV-2 viral infection and disease in asymptomatic individuals requires collaborations among helper CD4^+^ T cells, neutralizing antibodies, IFN-γ-producing effector CD4^+^ and cytotoxic CD8^+^ T cells ^45^. While SARS-CoV-2-induced antibody and CD4^+^ and CD8^+^ T cell responses are critical to reducing viral infection ^69^, in a majority of asymptomatic individuals, an excessive pro-inflammatory cytokine storm appears to lead to acute respiratory distress syndrome in many symptomatic individuals ^33–39^. This cytokine storm contributes to the immunopathology seen in severe COVID-19 patients, mandating caution in selecting a potentially protective “asymptomatic” CD4^+^ and CD8^+^ T cell epitopes to be incorporated in future pan-Coronavirus vaccines ^33–36, 70–72^. The “asymptomatic” epitopes recognized by CD4^+^ and CD8^+^ T cells associated with “natural protection” seen in asymptomatic individuals would be highly informative in the development of a safe and efficient pan-Coronavirus vaccine. Besides, these epitopes can also be used in diagnostics of human and animal Coronaviruses.

While SARS-CoV-2 appeared to originate from the natural reservoirs horseshoe bats (*Rhinolophus* spp.) there was a lack of clarity around how the virus was passed onto humans ^73^. Questions remain as to whether the highly contagious and deadly SARS-CoV-2 was: (1) transmitted naturally to humans through a yet-to-be identified intermediate animal reservoir; or (2) accidentally transmitted to humans as a “natural” but recombinant bat virus, or even as a man-made “artificial” recombinant bat virus ^18–20, 74^. Our study revealed that two SL-CoVs strains, called RATG13 and RmYN02, found in horseshoe bats *Rhinolophus affinis* that live in caves of Yunnan province, shared 96% of their genetic sequence with the sequences of over 81,000 SARS-CoV-2 strains ^19^ and were phylogenetically the closest to the first human SARS-CoV-2 strain reported from Wuhan (**Figs. 1B** and **1C**). Evidence suggests that the previous SARS-CoV passed on to humans from bats through civets, as intermediate animal reservoirs, while the previous MERS-CoV passed to humans from bats through camels as intermediate animal reservoir ^7, 75^. Sequence alignments in study identified the pangolin as the likely intermediate animal reservoir of SARS-CoV-2. Regardless of the intermediate animal reservoir of SARS-CoV-2, it appears that all the three deadly Coronavirus outbreaks in the 21^st^ century have originated from bats as natural hosts. The present study highlights the utility to identify highly conserved immune target epitopes that are common among the pre-pandemic SL-CoVs strains currently harbored in *Rhinolophus* spp., horseshoes bats and the post-pandemic human SARS-CoVs^14^. The conserved epitopes between human and animal Coronaviruses would be excellent immune targets to be incorporated in pre-emptive pan-Coronavirus vaccines.

It is inevitable that other COVID-like outbreaks, caused by yet another spillover of a bat SL-CoV, could lead to another COVID-like pandemic with global health, social and economic disasters in the years to come. However, because it is almost impossible to predict which viral strain might cause the next Coronavirus pandemic, it is urgent to develop a pan-Coronavirus vaccine that targets a wide range of human and animal Coronavirus strains. Unlike conventional monovalent vaccines made from epitopes selected from a single virus strain, our pre-emptive multi-epitope pan-Coronavirus vaccine (**Supplemental Fig. S5**) includes several highly conserved human B-, CD4^+^ and CD8^+^ T cell epitopes identified from the entire genome sequences of human SARS-CoVs that cross-react and shared with bat and pangolin SL-CoVs ^19, 76–79^. Unfortunately, for the past two decades, on one hand the governments decided that developing pre-emptive pan-Coronavirus vaccines is costly. On the other hand, private pharmaceutical companies, which mainly operate for profit, won’t invest in potentially unprofitable pre-emptive pan-Coronavirus vaccines for a “Disease X” that doesn’t exist yet, or that has yet to emerge. Thus, development of pre-emptive pan-vaccines against potential zoonotic viruses with a higher probability to emerge and spillover into humans should be funded by WHO and/or non-governmental nonprofit health organizations. The current ongoing collaborative research efforts should not only focus on developing a vaccine for COVID-19, but should also be oriented towards developing pre-emptive pan-Coronavirus vaccines. Such a proactive vaccine strategy would help fight and contain future local outbreaks and epicenters of highly contagious and deadly zoonotic Coronaviruses, globally, before they become a next deadly pandemic worldwide ^21, 61^. Moreover, because it is impossible to predict the time and location of next deadly global pandemic, it is essential to have ready, at least pre-clinically, pan-Coronavirus vaccine candidates that would be quickly implemented in a clinical trial against a substantial group of Coronaviruses before an outbreak spreads into global pandemics.

Our pre-emptive multi-epitope pan-Coronavirus vaccine is highly adaptable to new Coronavirus strains that may appear in the future. If an epitope from a human or an animal Coronavirus mutates, then that epitope can be easily adjusted and replaced in the multi-epitope vaccine with the new mutated epitope ^80^. Thus, it is not excluded that the highly conserved B-cell, CD4^+^ and CD8^+^ T cell epitopes identified in this study from bat’s Coronavirus variants will mutate, following recombination that often occurs for zoonotic events before an animal SL-CoV spills over into humans. Thus, our pre-emptive multi-epitope pan-Coronavirus vaccine strategy could be easily adapted, not only to any zoonotic bat SL-CoVs that may spill over into humans, but also to any mutations and shifts of SARS-CoV-2 variants that may emerge in the future. This high adaptability is expected to speed up the implementation of a future pre-emptive multi-epitope vaccine before an outbreak spreads into global pandemics. Besides, the multi-epitope vaccine can also be adapted to any antigen delivery system that will deliver the identified highly conserved SARS-CoV-2 epitopes ^81^.

In the present study, CD4^+^ and CD8^+^ T cells specific to highly conserved SARS-CoV2 epitopes were detected in healthy adults, recruited between 2014 and 2018, who have never been exposed to the SARS-CoV-2 virus. These findings suggest cross-reactive T cell responses between SARS-CoV-2 and circulating ‘‘common cold’’ Coronaviruses as confirmed by recent reports ^60, 82, 83^. However, since we do not have available history of whether the healthy adults used in our study were exposed to any ‘‘common cold’’ Coronaviruses, such an assertion may not be conclusive. Among the many circulating ‘‘common cold’’ Coronaviruses known to infect humans, four serotypes that cause severe respiratory infections are highly seasonal: CoV-OC43, CoV-229E, CoV-HKU1, and CoV-NL63 ^55^ and appear to have a similar transmission potential to influenza A (H3N2) but their seasonality was more predictable as their outbreaks often emerged in December, peaked in January/February, and began to decrease in March of each year ^55^. The human SARS-CoV-2 CD4^+^ and CD8^+^ T cell epitopes identified in this study are highly conserved between 81,963 strains of SARS-CoV-2 and CoV-OC43, CoV-229E, CoV-HKU1, and CoV-NL63 (**Figs. 2D** and **3**). Whether these apparent cross-reactive CD4^+^ and CD8^+^ T cells play a protective or a harmful role or an entirely negligible role SARS-CoV-2 infection and disease remains to be determined ^82, 83^. Nevertheless, since ‘‘common cold’’ Coronaviruses are usual in children, it will be interesting to determine whether children who appeared more resistant to COVID-19 compared to adults will have robust antiviral memory T cell responses to some asymptomatic epitopes. A stronger CD4^+^ and CD8^+^ T cell responses to common Coronavirus epitopes in children would shed some light on the unique situation currently seen in COVID-19 where immune children tend to be more resistant to SARS-CoV-2 infection and disease, as compared to adults ^84, 85^. Such a result would also imply that a pan-Coronavirus vaccine incorporating cross-reactive highly conserved SARS-CoV-2 human CD4^+^ and CD8^+^ T cell epitopes in the next pan-Coronavirus vaccine would boost protective T cell immunity that is previously induced by a ‘‘common cold’’ Coronavirus, thus protecting not only from seasonal circulating ‘‘common cold’’ Coronaviruses, but also from SARS-CoV-2 infection and disease. Currently, we are in the process of determining whether the individuals that were exposed to ‘‘common cold’’ Coronaviruses will develop frequent cross-reactive tissue-resident memory CD4^+^ and CD8^+^ T cells (T_RM_ cells) that would better protect them from SARS-CoV-2, compared to individuals who have never been exposed to ‘‘common cold’’ Coronaviruses. The results from those studies will be the subject of a future report. It is also likely that these cross-reactive SARS-CoV-2 human CD4^+^ and CD8^+^ T cells might be the result of a heterologous immunity with yet-to-be determined pathogen(s), within or outside the Coronavirus family, initiated by the development of these cross-reactive CD4^+^ and CD8^+^ T cells.

It was recently suggested that a majority of recovered patients produce antibodies against SARS-CoV-2 that would protect against re-infections ^86^. This concept leads to a so called “immunity passport” or “risk-free certificate” that would enable immune recovered individuals to return to work during the COVID-19 pandemic ^87^. However, the quality, the quantity and the epitope specificities of protective antibodies developed by immune recovered individuals remains to be yet determined ^88^. In a disagreement with a recent reports ^89, 90^, in this study we observed differences in the levels of IgG specific to SARS-CoV-2-Spike epitopes, with some spike-specific humoral responses were enriched among COVID-19 patients with severe symptoms, whereas asymptomatic COVID-19 patients develop rather lower levels of these IgG antibodies. This suggest that high titer IgG antibodies specific to some Spike epitopes might be involved in enhancing immunity while IgG specific to different Spike epitopes may be involved in protection. Accordingly, one cannot rule out that vaccination with the whole attenuated virus or even with whole proteins (e.g. Spike protein) can induce both protective and pathogenic immune responses. Such a vaccine may induce antibodies and T cells specific to “asymptomatic” epitopes that are protective while, at the same time, may induce antibodies and T cells specific to other “symptomatic” epitopes from the same protein that may actually accelerate the infection exacerbate the disease. Therefore, a multi-epitope vaccine that selectively incorporates selected “asymptomatic” B and T cell epitopes, while excluding “symptomatic” epitopes would be expected to protect from SARS-CoV-2 infection, while avoiding exacerbation of the infection and/or disease. Besides, unfortunately, the concept of “immunity passport” was mainly based on antibodies while ignoring the quality, the quantity and the epitope specificities of T cells developed by immune recovered individuals that may also be involved in protection from a second infection. The present report found that, in contrast to CD8^+^ T cells, we detected a significantly higher frequencies of memory CD4^+^ T cells specific to 8 highly conserved epitopes in asymptomatic COVID-19 patients compared to symptomatic COVID-19 patients, regardless of level of severity of symptoms. This. it is likely that, besides antibodies, the SARS-CoV-2 infection may simultaneously induce protective and pathogenic T cells specific to asymptomatic and symptomatic epitopes, respectively.

The highly contagious and transmissibility characteristics of SARS-CoV-2, compared to previous Coronavirus outbreaks, is likely due to its high ability to mutate ^91^. Dynamic tracking of variant frequencies revealed that the originally SARS-CoV-2 variant that appeared in January 2020 in Wuhan, China has a D amino acid in position 614 of the spike protein (D614) mutated into a G amino acid in the same position 614 of the spike protein (D614) while traveling to Europe and then to the Americas^41^. The resulting new G614 SARS-CoV-2 variant may have a fitness advantage and appears to be the dominant pandemic variant in Europe and in the United States, leading to a higher magnitude of infection, a higher upper respiratory tract viral loads (although not with increased disease severity), compared to the original Wuhan SARS-CoV-2 variant ^41^. The SARS-CoV-2 variant carrying the Spike protein amino acid change D614G has become the most prevalent form in the global pandemic ^40, 41^. In this context, our anti-viral multi-epitope pan-Coronavirus vaccine candidate #1, which includes the Spike B8 epitope with D amino acid in position 14 (**Fig. 9** and **Supplemental Fig. S5**), could be adapted to target the new G614 SARS-CoV-2 variant by replacing the mutated B8 epitope with a G amino acid in the position 614. Thus, our multi-epitope pan-Coronavirus vaccine strategy could be easily adapted, not only to any zoonotic bat SL-CoVs that may spill over into humans in the future, but also to any mutations and shifts of future SARS-CoV-2 variants. Evidence of strong purifying selection around the receptor binding domains (RBD) in the Spike gene and in other genes among bat, pangolin and human Coronaviruses, indicating similarly strong evolutionary constraints in different host species has been reported ^40, 41^.

Even though the highly conserved Coronavirus human CD4^+^ and CD8^+^ T cell epitopes identified in this report can be enlightening for a pan-Coronavirus vaccine, humans are not immunologically naive, and they often have memory CD4^+^ and CD8^+^ T cell populations that can cross-react with, and respond to, other infectious agents, a phenomenon termed heterologous immunity. Therefore, we cannot exclude that some SARS-CoV-2-specific CD4^+^ and CD8^+^ T cell epitopes identified in this study are cross-reactive with other viral pathogens-derived epitopes, such as epitopes from circulating seasonal influenza or ‘‘common cold’’ Coronaviruses ^92, 93^. This may explain, in part, the high proportion of asymptomatic infections with SARS-CoV-2 in the current pandemic. The latter is supported by a recent elegant study that detected SARS-CoV-2-reactive CD8^+^ and CD4^+^ T cells in healthy individuals that were never exposed to SARS-CoV-2 ^60^. SARS-CoV-2-specific, but cross-reactive, CD4^+^ and CD8^+^ T cells can become activated and modulate the immune responses and clinical outcome of subsequent heterologous SARS-CoV-2 infections. Therefore, T cell cross-reactivity may be crucial in protective heterologous immunity instead of damaging heterologous immunopathology, as has been reported in other systems ^94^. To confirm SARS-CoV-2 heterologous CD4^+^ and CD8^+^ T cell epitopes that may potentially cross-react with other pathogenic (non-Coronaviruses) epitopes, we are currently comparing the CD4^+^ and CD8^+^ T-cell response to those highly conserved SARS-CoV-2 epitopes identified using humans CD4^+^ and CD8^+^ T-cell responses to those of “pathogen-free” SARS-CoV-2-infected transgenic mice.

In summary, we report several human “universal” B, and CD4^+^ and CD8^+^ T cell target epitopes identified from the whole SARS-CoV-2 genome that are highly conserved and common between SARS-CoV-2 Wuhan strain and: (1) circulating ‘‘common cold’’ human Coronaviruses that caused previous human SARS and MERS outbreaks ^61^; (2) 81,963 strains of human SARS-CoV-2 that now circulate in six continents; (3) bat-derived SARS-like strains ^14, 15^; and (4) SL-CoV strains isolated from pangolins ^95^. While the current COVID-19 pandemic will likely disappear through implementation physical distancing and mass vaccination, another COVID pandemic will likely emerge in coming years (the question is not “if” the question is “when”). This work paves the way for the design and evaluation of “pre-emptive” pan-Coronavirus vaccine candidates that will target not only the current human SARS-CoV-2, but also possible future bat-derived SARS-like Coronavirus strains, that might transition and spill over into humans, thus potentially causing future global outbreaks.

## MATERIALS & METHODS

### Human study population

Sixty-three COVID-19 patients and ten unexposed healthy individuals, who had never been exposed to SARS-CoV-2 or COVID-19 patients, were enrolled in this study (**Table 1**). Seventy-eight percent were non-White (African, Asian, Hispanic and others) and 22% were white. Forty-four percent were females, and 56% were males with an age range of 26-95 (median 62)

Detailed clinical and demographic characteristics of the symptomatic vs. asymptomatic COVID-19 patients and the unexposed healthy individuals with respect to age, gender, HLA-A*02:01 and HLA-DR distribution, COVID-19 disease severity, comorbidity and biochemical parameters are detailed in **Table 1**. None of the symptomatic patients were on anti-viral or anti-inflammatory drug treatments at the time of blood sample collections. The symptomatic COVID-19 patients were divided into 4 groups depending on the severity of the symptoms: Group 1 that comprised of SARS-CoV-2 infected patients that never developed any symptoms or any viral diseases (i.e. asymptomatic patients) (*n* = 11); Group 2 with mild symptoms (i.e. Inpatient only, *n* = 32); Group 3 with moderate symptoms (i.e. ICU admission, *n* = 11) and Group 4 with severe symptoms (i.e. ICU admission +/− Intubation or death, *n* = 9). As expected, compared to the asymptomatic group, all of the 3 symptomatic groups (i.e. mild, moderate and severe) had higher percentages of comorbidities, including diabetes (22% to 64%), hypertension (64% to 78%), cardiovascular disease (11% to 18%) and obesity (9% to 50%) (**Table 1**). The final Group 5 was comprised of unexposed healthy individuals (controls), with no history of COVID-19 or contact with COVID-19 patients (*n* = 10). All subjects were enrolled at the University of California Irvine under Institutional Review Board-approved protocols (IRB # 2020-5779). A written informed consent was received from all participants prior to inclusion in the study.

### Sequence comparison among SARS-CoV-2 and previous Coronavirus strains

We retrieved 81,963 human SARS-CoV-2 genome sequences from GISAID database representing countries from North America, South America, Central America, Europe, Asia, Oceania, and Africa (**Fig. 1**). Furthermore, the full length sequences of SARS-CoV strains (SARS-CoV-2-Wuhan-Hu-1 (MN908947.3), SARS-CoV-Urbani (AY278741.1), HKU1-Genotype B (AY884001), CoV-OC43 (KF923903), CoV-NL63 (NC_005831), CoV-229E (KY983587)) and MERS (NC_019843)) found in the human host were obtained from the NCBI GeneBank. SARS-CoV-2 genome sequences from bat (RATG13, ZXC21, YN01. YN02), and pangolin (GX-P2V, GX-P5E, GX-P5L, GX-P1E, GX-P4L, GX-P3B, MP789, Guangdong-P2S) were obtained from NCBI and GSAID. More-so, SARS-CoV strains from bat (WIV16, WIV1, YNLF_31C, Rs672, recombinant strain FJ211859.1), camel (KT368891.1, MN514967.1, KF917527.1, NC_028752.1), and civet (Civet007, A022, B039)) were also retrieved from the NCBI GeneBank. The sequences were aligned using ClustalW algorithm in MEGA X.

### Sequence conservation analysis of SARS-CoV-2

The SARS-CoV-2-Wuhan-Hu-1 (MN908947.3) protein sequence was compared with SARS-CoV and MERS-CoV specific protein sequences obtained from human, bat, pangolin, civet and camel. The Sequence Variation Analysis was performed on the consensus aligned protein sequences from each virus strain. This Sequence Homology Analysis identified consensus protein sequences from the SARS-CoV and MERS-CoV and predicted the Epitope Sequence Analysis.

### SARS-CoV-2 CD8 and CD4 T Cell Epitope Prediction

Epitope prediction was carried out using the twelve proteins predicted for the reference SARS-CoV-2 isolate, Wuhan-Hu-1. The corresponding SARS-CoV-2 protein accession identification numbers are: YP_009724389.1 (ORF1ab), YP_009725295.1 (ORF1a), YP_009724390.1 (surface glycoprotein), YP_009724391.1 (ORF3a), YP_009724392.1 (envelope protein), YP_009724393.1 (membrane glycoprotein), YP_009724394.1 (ORF6), YP_009724395.1 (ORF7a), YP_009725318.1 (ORF7b), YP_009724396.1 (ORF8), YP_009724397.2 (nucleocapsid phosphoprotein), YP_009725255.1 (ORF10). The tools used for CD8^+^ T cell-based epitope prediction were SYFPEITHI, MHC-I binding predictions, and Class I Immunogenicity. Of these, the latter two were hosted on the IEDB platform. For the prediction of CD4^+^ T cell epitopes, we used multiple databases and algorithms, namely SYFPEITHI, MHC-II Binding Predictions, Tepitool, and TEPITOPEpan. For CD8^+^ T cell epitope prediction, we selected the 5 most frequent HLA-A class I alleles (HLA-A*01:01, HLA-A*02:01, HLA-A*03:01, HLA-A*11:01, HLA-A*23:01) with large coverage of the world population, regardless of race and ethnicity (**Supplemental Figs. S1A** and **S1C**) (Middleton et al., 2003), using a phenotypic frequency cutoff ≥ 6%. Similarly, for CD4 T cell epitope prediction, selected HLA-DRB1*01:01, HLA-DRB1*11:01, HLA-DRB1*15:01, HLA-DRB1*03:01, HLA-DRB1*04:01 alleles with large population coverage (**Supplemental Figs. S1B** and **S1D**). Subsequently, using NetMHC we analyzed the SARS-CoV-2 protein sequence against all the aforementioned MHC-I and MHC-II alleles. Epitopes with 9mer length for MHC-I and 15mer length for MHC-II were predicted. Subsequently, the peptides were analyzed for binding stability to the respective HLA allotype. Our stringent epitope selection criteria was based on picking the top 1% epitopes focused on prediction percentile scores.

### SARS-CoV-2 B Cell Epitope Prediction

Linear B cell epitope predictions were carried out on the surface glycoprotein (S), the primary target of B cell immune responses for SARS-CoV. We used the BepiPred 2.0 algorithm embedded in the B cell prediction analysis tool hosted on IEDB platform. For each protein, the epitope probability score for each amino acid and the probability of exposure was retrieved. Potential B cell epitopes were predicted using a cutoff of 0.55 (corresponding to a specificity greater than 0.81 and sensitivity below 0.3) and considering sequences having more than 5 amino acid residues. This screening process resulted in 28 B-cell peptides (**Supplemental Table S3**). From this pool, we selected 10 B-cell epitopes with 19 to 62 amino acid lengths in our current study. Three B-cell epitopes were observed to possess receptor binding domain (RBD) region specific amino acids. Structure-based antibody prediction was performed by using Discotope 2.0, and a positivity cutoff greater than −2.5 was applied (corresponding to specificity greater than or equal to 0.80 and sensitivity below 0.39), using the SARS-CoV-2 spike glycoprotein structure (PDB ID: 6M1D).

### Protein-peptide molecular docking

Computational peptide docking of B cell peptides into the ACE2 Complex (binding protein) was performed using the GalaxyPepDock under GalaxyWEB. To retrieve the ACE2 structure, we used the X-ray crystallographic structure ACE2-B0AT1 complex-6M1D available on the Protein Data Bank. The 6M1D with a structural weight of 334.09 kDa, possesses 2 unique protein chains, 2,706 residues, and 21,776 atoms. In this study, flexible target docking based on an energy-optimization algorithm was carried out on the ligand-binding domain containing ACE2 within the 4GBX structure. Similarity scores were calculated for protein-peptide interaction pairs for each residue. The prediction accuracy is estimated from a linear model as the relationship between the fraction of correctly predicted binding site residues and the template-target similarity measured by the protein structure similarity score and interaction similarity score (S_Inter_) obtained by linear regression. S_Inter_ shows the similarity of amino acids of the B-cell peptides aligned to the contacting residues in the amino acids of the ACE2 template structure. Higher S_Inter_ score represents a more significant binding affinity among the ACE2 molecule and B-cell peptides. Subsequently, molecular docking models were built based on distance restraints for protein-peptide pairs using GalaxyPepDock. Based on optimized energy scores docking models were ranked.

While performing the protein-peptide docking analysis for CD8^+^ T cell epitope peptides, we used the X-ray Crystal structure of HLA-A*02:01 in complex-4UQ3 available on the Protein Data Bank and for CD4 peptides X-ray crystallographic structure HLA-DM-HLA-DRB1 Complex-4GBX.

### Epitope conservancy analysis

The Epitope Conservancy Analysis tool was used to compute the degree of conservancy of CD8^+^ T cell, CD4^+^ T cell, and B-cell epitopes within a given protein sequence of SARS-CoV-2 set at 100% identity level. The fraction of protein sequences that contain the epitope, and identity was defined as the degree of similarity or correspondence among two sequences. The CD8^+^ T cell, and CD4^+^ T cell epitopes were screened against all the twelve structural and non-structural proteins of SARS-CoV-2 namely YP_009724389.1 (ORF1ab), YP_009725295.1 (ORF1a), YP_009724390.1 (surface glycoprotein), YP_009724391.1 (ORF3a), YP_009724392.1 (envelope protein), YP_009724393.1 (membrane glycoprotein), YP_009724394.1 (ORF6), YP_009724395.1 (ORF7a), YP_009725318.1 (ORF7b), YP_009724396.1 (ORF8), YP_009724397.2 (nucleocapsid phosphoprotein), YP_009725255.1 (ORF10). Whereas, the B-cell epitopes were screened for their conservancy against surface glycoprotein (YP_009724390.1) of SARS-CoV-2. Epitope linear sequence conservancy approach was used for linear epitope sequences with a sequence identity threshold set at ≥ 50%. This analysis resulted in (*i*) the calculated degree of conservancy (percent of protein sequence matches a specified identity level) and (*ii*) the matching minimum/maximum identity levels within the protein sequence set. The CD8^+^ T cell epitopes, which showed ≥ 50% conservancy in at-least two human SARS-CoV strains and two SARS-CoV strains (from bat/civet/pangolin/camel) were selected as candidate epitopes.

### Population-Coverage-Based T Cell Epitope Selection

For a robust epitope screening, we evaluated the conservancy of CD8^+^ T cell, CD4^+^ T cell, and B cell epitopes with Human-SARS-CoV-2 genome sequences representing North America, South America, Africa, Europe, Asia, and Australia. As of August 27^th^, 2020, the NextStrain database recorded 81,963 human-SARS-CoV-2 genome sequences and the number of genome sequences continues growing daily. In the present analysis, 81,963 human-SARS-CoV-2 genome sequences were extrapolated from the GISAID and NCBI GeneBank databases. We therefore considered all the 81,963 SARS-CoV-2 genome sequences representing six continents for subsequent conservancy analysis. We set a threshold for a candidate CD8^+^ T cell, CD4^+^ T cell, and B-cell epitope if the epitope showed 100% sequence conservancy in ≥ 95 human-SARS-CoV-2 genome sequences. Further, population coverage calculation (PPC) was carried out using the Population Coverage software hosted on IEDB platform ^96^ PPC was performed to evaluate the distribution of screened CD8^+^ and CD4^+^ T cell epitopes in world population at large in combination with HLA-I (HLA-A*01:01,HLA-A*02:01,HLA-A*03:01,HLA-A*11:01,HLA-A*23:01), and HLA-II (HLA-DRB1*01:01, HLA-DRB1*11:01, HLA-DRB1*15:01, HLA-DRB1*03:01, HLA-DRB1*04:01) alleles.

### Peptide synthesis

Potential peptide epitopes (9-mer long for CD8^+^ T cell epitopes and 15-mer long for CD4^+^ T cell epitopes) identified from twelve human-SARS-CoV-2 proteins namely orf1ab, orf1a, surface glycoprotein, Orf3a, envelope protein, membrane glycoprotein, ORF6, ORF7a, ORF7b, ORF8, nucleocapsid phosphoprotein, and ORF10 were synthesized using solid-phase peptide synthesis and standard 9-fluorenylmethoxycarbonyl technology (21^st^ Century Biochemicals, Inc). The purity of peptides was over 90%, as determined by reversed-phase high-performance liquid chromatography (Vydac C18) and mass spectroscopy (VOYAGER MALDI-TOF System). Stock solutions were made at 1 mg/mL in 10% DMSO in PBS. Similar method of synthesis was used for B cell peptide epitopes from Spike protein of SARS-CoV-2.

### Cell Lines

T_2_ (174 × CEM.T2) mutant hybrid cell line derived from the T-lymphoblast cell line CEM was obtained from the ATCC (www.atcc.org). The T_2_ cell line was maintained in IMDM (ATCC) supplemented with 10% heat-inactivated fetal calf serum (FCS) and 100 U of penicillin/mL, 100 U of streptomycin/mL (Sigma-Aldrich, St. Louis, MO).T_2_ cells lack the functional transporter associated with antigen processing (TAP) heterodimer and failed to express normal amounts of HLA-A*02:01 on the cell surface. HLA-A*02:01 surface expression is stabilized following binding of exogenous peptides to these MHC class I molecules.

### Stabilization of HLA-A*0201 on class-I-HLA-transfected B x T hybrid cell lines

To determine whether synthetic peptides could stabilize HLA-A*0201 molecule expression on the T_2_ cell surface, peptide-inducing HLA-A*0201 up-regulation on T_2_ cells was examined according to a previously described protocol ^97, 98^. T_2_ cells (3 × 10^5^/well) were incubated with different concentrations (30, 10 and 3µM) of 91 individual CD8^+^ T cell specific peptides in 48-well plates for 18 hours at 26°C. Cells were then incubated at 37°C for 3 hours in the presence of 0.7 μL/mL BD GolgiStop^™^ to block cell surface expression of newly synthesized HLA-A*0201 molecules, and human β-2 microglobulin (1μg/mL). The cells were subsequently washed with FACS buffer (1% BSA and 0.1% sodium azide in phosphate-buffered saline) and stained with anti-HLA-A2 specific monoclonal antibody (clone BB7.2) (BD-Pharmingen, San Diego, CA) at 4°C for 30 minutes. After incubation, the cells were washed with FACS buffer, fixed with 2% paraformaldehyde in phosphate-buffered saline, and analyzed by flow cytometry using a Fortessa (Becton Dickinson) flow cytometer equipped with a BD High Throughput Sampler for rapid analysis of samples prepared in plate format. The acquired data were analyzed with FlowJo software (BD Biosciences, San Jose, CA) and expression was measured by mean fluorescence intensity (MFI). Percent MFI increase was calculated as follows: Percent MFI increase = (MFI with the given peptide - MFI without peptide) / (MFI without peptide) x 100. Each experiment was performed 3 times, and means ± SD values were calculated.

### HLA-A*02:01 and HLA-DR1 double transgenic mice

A colony of human leukocyte antigens (HLA) class I and class II double transgenic (Tg) mice is maintained at UC Irvine ^99^ and were housed and treated in accordance the AAALAC (Association for Assessment and Accreditation of Laboratory Animal Care), and NIH (National Institutes of Health) guidelines. The HLA Tg mice retain their endogenous mouse major histocompatibility complex (MHC) locus and express human HLA-A*02:01 and HLA-DRB:01 under the control of its normal promoter ^100, 101^. Prior to this study, the expression of HLA-A*02:01 molecules on the PBMCs of each HLA-Tg mouse was confirmed by fluorescence-activated cell sorting (FACS). All animal experiments were conducted following NIH guidelines for housing and care of laboratory animals and were conducted in facilities approved by the University of California Irvine Association for Assessment and Accreditation of Laboratory Animal Care and according to Institutional Animal Care and Use Committee-approved animal protocols (IACUC # 2020-19-111).

### Immunization of mice

Groups of age-matched HLA transgenic mice/B6 mice (*n* = 3) were immunized subcutaneously, on days 0 and 14, with a mixture of four SARS-CoV-2-derived human CD4^+^ T/ CD8^+^T /B cell peptide epitopes delivered in alum and CpG_1826_ adjuvants. As a negative control, mice received adjuvants alone (mock-immunized).

### Splenocytes isolation

Spleens were harvested from mice two weeks post second immunization. Spleens were placed in 10 ml of cold PBS with 10% fetal bovine serum (FBS) and 2X antibiotic–antimycotic (Life Technologies, Carlsbad, CA). Spleens were minced finely and sequentially passed through a 100 µm screen and a 70 µm screen (BD Biosciences, San Jose, CA). Cells were then pelleted via centrifugation at 400 × *g* for 10 minutes at 4°C. Red blood cells were lysed using a lysis buffer (ammonium chloride) and washed again. Isolated splenocytes were diluted to 1 × 10^6^ viable cells per ml in RPMI media with 10% (v/v) FBS and 2 × antibiotic–antimycotic. Viability was determined by trypan blue staining.

### Flow cytometry analysis

*PBMCs**/***Splenocytes were analyzed by flow cytometry. The following antibodies were used: CD8, CD4, CD62L, CD107^a/b^, CD44, CD69, TNF-α and IFN-γ). For surface staining, mAbs against various cell markers were added to a total of 1X10^6^ cells in phosphate-buffered saline containing 1% FBS and 0.1% sodium azide (fluorescence-activated cell sorter [FACS] buffer) and left for 45 minutes at 4°C. At the end of the incubation period, the cells were washed twice with FACS buffer. A total of 100,000 events were acquired by LSRII (Becton Dickinson, Mountain View, CA) followed by analysis using FlowJo software (TreeStar, Ashland, OR).

### ELISpot assay

All reagents used were filtered through a 0.22 µm filter. Wells of 96-well Multiscreen HTS Plates (Millipore, Billerica, MA) were pre-wet with 70% methanol and then coated with 100 µl primary anti-IFN-γ antibody solution (10 µg/ml of 1-D1K coating antibody from Mabtech in PBS, pH 7.4, V-E4) OVN at 4°C. After washing, nonspecific binding was blocked with 200 µl of RPMI media with 10% (v/v) FBS for 2 hours at room temperature. Following the blockade, 0.5 × 10^6^ cells from patients PBMCs (or from mouse splenocytes) in 100 µl of RPMI were mixed with 10ug individual peptides (with DMSO for no stimulation or with individual peptide at a final concentration of 10 µg/ml). After incubation in humidified 5% CO_2_ at 37°C for 72 hours, cells were removed by washing (using PBS and PBS-Tween 0.02% solution) and 100 µl of biotinylated secondary anti-IFN-γ antibody (clone 7-B6-1, Mabtech) in blocking buffer (PBS 0.5% BSA) was added to each well. Following a 2 hour incubation and washing, HRP-conjugated streptavidin was diluted 1:1000 and wells were incubated with 100 µl for 1 hour at room temperature. Following washing, wells were incubated for 1 hour at room temperature with 100 µl of TMB detection reagent and spots counted with an automated EliSpot Reader System (ImmunoSpot reader, Cellular Technology).

### Efficacy of receptor binding domain region towards inducing specific antibodies against B-cell epitopes in HLA-A2 treated mice

ELISA plates (Cat. M5785, Sigma Aldrich) were first coated overnight at 4°C with 10μg/ml of each B cell peptide epitope. Subsequently plates were washed five times with PBS-Tween 0.01% before to start the blocking by adding PBS 1% BSA for 3 h at room temperature, followed by a second wash. Sera of C57BL/6 mice immunized either with pool B cell peptides alum/CpG or adjuvant alone (control) were added into the wells at various dilutions (1/5, 1/25, 1/125 and 1/625 or PBS only, in triplicate). Plates were incubated at 4°C overnight with the sera, then washed with PBS-Tween 0.01% before to add anti-mouse IgG antibody (Mabtech – 1/500 dilution). After the last washing, Streptavidin-HRP (Mabtech – 1/1000 dilution) was added for 30 minutes at room temperature. Finally, we added 100μl of filtered TMB substrate for 15 minutes and blocked the reaction with H_2_S0_4_ before the read-out (OD measurement was done at 450nm on the Bio-Rad iMark microplate reader).

### Constructing the Phylogenetic Tree

Phylogenetic analyses were conducted in MEGA X. The evolutionary history was performed, and phylogenetic tree was constructed by using the Maximum Likelihood method and Tamura-Nei model. The Maximum Likelihood method assumes that each locus evolves independently by pure genetic drift. The tree with the highest log likelihood was selected. Initial tree(s) for the heuristic search were obtained by applying Neighbor-Join and BioNJ algorithms to a matrix of pairwise genetic distances estimated using the Tamura-Nei model, and then selecting the topology with superior log likelihood value. This analysis involved available nucleotide sequences of SARS-CoV-2 from human (*Homo Sapiens*), bat (*Rhinolophus affinis*, *Rhinolophus malayanus*), and pangolin (*Manis javanica*). In addition, genome sequences from previous outbreaks of SARS-CoV in human, bat, civet, and camel were taken into consideration while performing the evolutionary analyses.

### Data and Code Availability

The human specific SARS-CoV-2 complete genome sequences were retrieved from the GISAID database, whereas the SARS-CoV-2 sequences for pangolin (*Manis javanica*), and bat (*Rhinolophus affinis*, *Rhinolophus malayanus*) were retrieved from NCBI. Genome sequences of previous strains of SARS-CoV for human, bat, civet, and camel were retrieved from the NCBI GeneBank.

### Statistical analyses

Data for each differentially expressed gene among blockade-treated, and mock-treated groups of HLA Tg mice were compared by analysis of variance (ANOVA) and Student’s *t*-test using GraphPad Prism version 6 (La Jolla, CA). Statistical differences observed in the measured CD8-, CD4-T cells and antibody responses between heatlhy donors and COVID-19 patients or across the different group of disease severity were calculated using ANOVA and multiple *t*-test comparison procedures in GraphPad Prism. Data are expressed as the mean ± SD. Results were considered statistically significant at *P* ≤ 0.05.

## ACKNOWLEDGMENTS

This work is supported by the Fast-Grant PR12501 from Emergent Ventures, by a Gavin Herbert Eye Institute internal grant, by Public Health Service Research grants AI158060, AI150091, AI143348, AI147499, AI143326, AI138764, AI124911 and AI110902 from the National Institutes of Allergy and Infectious Diseases (NIAID) to LBM.

The authors would like to thank Dr. Dale Long from the NIH Tetramer Facility (Emory University, Atlanta, GA) for providing the Tetramers used in this study. We thank UC Irvine Center for Clinical Research (CCR) and Institute for Clinical & Translational Science (ICTS) for providing human blood samples used in this study. A special thanks to Dr. Delia F. Tifrea for her continuous efforts and dedication in providing COVID-19 samples that are crucial for this clinical research. We also thank those who contributed directly or indirectly to this COVID-19 vaccine project: Dr. Steven A. Goldstein, Dr. Michael J. Stamos, Dr. Suzanne B. Sandmeyer, Jim Mazzo, Dr. Daniela Bota, Dr. Beverly L. Alger, Dr. Dan Forthal, Dr. Tahseen Muzaffar, Dr. Ilhem Messaoudi, Anju Subba, Janice Briggs, Marge Brannon, Beverley Alberola, Jessica Sheldon, Rosie Magallon and Andria Pontello.

---

**Supplemental Figure S1: Population coverage calculation for all the high binding CD8^+^ and CD4^+^ T cell epitopes screened in the current study:** (A) The SARS-CoV-2-derived CD8^+^ T cell epitopes are screened based on their presence among most frequently observed HLA-A alleles in global population (HLA-A*01:01, HLA-A*02:01, HLA-A*03:01, HLA-A*11:01, HLA-A*23:01). MHC-I binding affinity and high degree of immunogenicity showed high PPC value of 75.66%, average number of epitope hits / HLA combinations recognized by the population is 22.16, and minimum number of epitope hits / HLA combinations recognized by 90% of the population (pc90) is 9.86 (B) The SARS-CoV-2-derived CD4^+^ T cell epitopes are screened based on their presence among most frequently observed HLA-DRB1 alleles in global population (HLA-DRB1*01:01, HLA-DRB1*11:01, HLA-DRB1*15:01, HLA-DRB1*03:01, HLA-DRB1*04:01). MHC-II binding affinity and high degree of immunogenicity showed high PPC values at 59.25%, average number of epitope hits / HLA combinations recognized by the population is 11.13, and minimum number of epitope hits / HLA combinations recognized by 90% of the population (pc90) is 3.96. The PPC plot shows percentage of individuals possessing combination of a number of epitopes/HLA alleles of interest and represents the cumulative percentage of population coverage. The line (-o-) shows the cumulative percentage of population coverage of epitope/HLA combination while the bars depict the population coverage for individual epitope. The analysis also generates the average number of epitope hits / HLA combinations recognized by the world population, and minimum number of epitope hits / HLA combinations recognized by 90% of the world population (pc90) as shown below the PPC plot. (C) The SARS-CoV-2-derived CD8^+^ T cell epitopes are screened based on their presence among most frequently observed HLA-A alleles in global population (HLA-A*01:01, HLA-A*02:01, HLA-A*03:01, HLA-A*11:01, HLA-A*23:01). MHC-I binding affinity and high degree of immunogenicity showed high PPC value of 75.66%, average number of epitope hits / HLA combinations recognized by the population is 14.13, and minimum number of epitope hits / HLA combinations recognized by 90% of the population (pc90) is 6.16. (D) The SARS-CoV-2-derived CD4^+^ T cell epitopes are screened based on their presence among most frequently observed HLA-DRB1 alleles in global population (HLA-DRB1*01:01, HLA-DRB1*11:01, HLA-DRB1*15:01, HLA-DRB1*03:01, HLA-DRB1*04:01). MHC-II binding affinity and high degree of immunogenicity showed high PPC value at 59.25%, average number of epitope hits / HLA combinations recognized by the population is 4.17, and minimum number of epitope hits / HLA combinations recognized by 90% of the population (pc90) is 1.47. The PPC plot shows percentage of individuals possessing combination of a number of epitopes/HLA alleles of interest and represents the cumulative percentage of population coverage. The line (-o-) shows the cumulative percentage of population coverage of epitope/HLA combination while the bars depict the population coverage for individual epitope. The analysis also generates the average number of epitope hits / HLA combinations recognized by the world population, and minimum number of epitope hits / HLA combinations recognized by 90% of the world population (pc90) as shown below the PPC plot.

**Supplemental Figure S2: Docking of highly conserved SARS-CoV-2-derived human CD8^+^ T cell epitopes to HLA-A*02:01 molecules:** (A) Docking of the 24 high-affinity CD8^+^ T cell binder peptides to the groove of HLA-A*02:01 molecules. (B) Summary of the interaction similarity scores of the 24 high-affinity CD8^+^ T cell epitope peptides to HLA-A*02:01 molecules determined by protein-peptide molecular docking analysis. Black columns depict CD8^+^ T cell epitope peptides with high interaction similarity scores.

**Supplemental Figure S3: Molecular docking of highly conserved SARS-CoV-2 CD4^+^ T cell epitopes to HLA-DRB1 molecules:** (A) Molecular docking of 16 CD4^+^ T cell epitopes, conserved among human SARS-CoV-2 strains, previous humans SARS/MERS-CoV and bat SL-CoVs into the groove of the HLA-DRB1 protein crystal structure (PDB accession no: 4UQ3) was determined using the GalaxyPepDock server. The 16 CD4^+^ T cell epitopes are promiscuous restricted to HLA-DRB1*01:01, HLA-DRB1*11:01, HLA-DRB1*15:01, HLA-DRB1*03:01 and HLA-DRB1*04:01 alleles. The CD4^+^ T cell peptides are shown in ball and stick structures, and the HLA-DRB1 protein crystal structure is shown as a template. The prediction accuracy is estimated from a linear model as the relationship between the fraction of correctly predicted binding site residues and the template-target similarity measured by the protein structure similarity score (TM score) and interaction similarity score (S_Inter_) obtained by linear regression. SInter shows the similarity of the amino acids of the CD8^+^ T cell peptides aligned to the contacting residues in the amino acids of the HLA-DRB1 template structure. (B) Histograms representing interaction similarity score of CD4^+^ T cells specific epitopes observed from the protein-peptide molecular docking analysis.

**Supplemental Figure S4: Docking of SARS-CoV-2 Spike glycoprotein-derived B cell epitopes to human ACE2 receptor:** (A) Molecular docking of 17 B-cell epitopes, identified from the SARS-CoV-2 Spike glycoprotein, with ACE2 receptors. B cell epitope peptides are shown in ball and stick structures whereas the ACE2 receptor protein is shown as a template. S_471-501_ and S_369-393_ peptide epitopes possess receptor binding domain region specific amino acid residues. The prediction accuracy is estimated from a linear model as the relationship between the fraction of correctly predicted binding site residues and the template-target similarity measured by the protein structure similarity score and interaction similarity score (S_Inter_) obtained by linear regression. S_Inter_ shows the similarity of amino acids of the B-cell peptides aligned to the contacting residues in the amino acids of the ACE2 template structure. Higher S_Inter_ score represents a more significant binding affinity among the ACE2 molecule and B-cell peptides. (B) Summary of the interaction similarity score of 17 B cells specific epitopes observed from the protein-peptide molecular docking analysis. B cell epitopes with high interaction similarity scores are indicated in black.

**Supplemental Table S1: Genome-wide screening of CD8^+^ T cell epitopes in human SARS-CoV-2 strain:** MHC-I binding prediction tool was used to screen 9,660 CD8^+^ T cell epitopes from 12 SARS-CoV-2 proteins YP_009724389.1 (ORF1ab), YP_009725295.1 (ORF1a), YP_009724390.1 (surface glycoprotein), YP_009724391.1 (ORF3a), YP_009724392.1 (envelope protein), YP_009724393.1 (membrane glycoprotein), YP_009724394.1 (ORF6), YP_009724395.1 (ORF7a), YP_009725318.1 (ORF7b), YP_009724396.1 (ORF8), YP_009724397.2 (nucleocapsid phosphoprotein), YP_009725255.1 (ORF10). The CD8+ T cell epitopes were screened against HLA-A*01:01, HLA-A*02:01, HLA-A*03:01, HLA-A*11:01, HLA-A*23:01 alleles. For each epitope corresponding MHC-I Allele, source of SARS-CoV-2 protein, start and end sequence position of the epitope, peptide score, and percentile rank have been provided. A small numbered percentile rank indicates high affinity.

**Supplemental Table S2: Genome-wide screening of CD4^+^ T cell epitopes in human SARS-CoV-2 strain:** MHC-II binding prediction tool was used to screen 9,594 CD4^+^ T cell epitopes from 12 SARS-CoV-2 proteins YP_009724389.1 (ORF1ab), YP_009725295.1 (ORF1a), YP_009724390.1 (surface glycoprotein), YP_009724391.1 (ORF3a), YP_009724392.1 (envelope protein), YP_009724393.1 (membrane glycoprotein), YP_009724394.1 (ORF6), YP_009724395.1 (ORF7a), YP_009725318.1 (ORF7b), YP_009724396.1 (ORF8), YP_009724397.2 (nucleocapsid phosphoprotein), YP_009725255.1 (ORF10). The CD4^+^ T cell epitopes were screened against HLA-DRB1*01:01, HLA-DRB1*03:01, HLA-DRB1*04:01, HLA-DRB1*11:01, and HLA-DRB1*15:01 alleles. For each epitope corresponding MHC-II Allele, source of SARS-CoV-2 protein, start and end sequence position of the epitope, peptide score, and percentile rank have been provided. A small numbered percentile rank indicates high affinity.

**Supplemental Table S3: Screening of B-cell epitopes on surface glycoprotein in human SARS-CoV-2 strain:** Antibody Epitope Prediction platform hosted on IEDB was used to screen linear B-cell peptides based on sequence characteristics of the antigen using amino acid scales. 28 B-cell epitopes with ≥ 5 bp amino acid length are screened through this process. The corresponding start and end positions in the surface glycoprotein (YP_009724390.1) for the epitopes have been provided. The tables provide values of calculated scores for each residue. The larger score for the residues might be interpreted as that the residue might have a higher probability to be part of epitope. The residues with scores above the threshold ≥ 0.5 are predicted to be part of an epitope marked with “E” in the output table.

## REFERENCES

1. Jeyanathan, M. et al. Immunological considerations for COVID-19 vaccine strategies. Nat Rev Immunol 20, 615–632 (2020).

2. Carroll, D. et al. Building a global atlas of zoonotic viruses. Bull World Health Organ 96, 292–294 (2018).

3. Allen, T. et al. Global hotspots and correlates of emerging zoonotic diseases. Nat Commun 8, 1124 (2017).

4. Valitutto, M.T. et al. Detection of novel coronaviruses in bats in Myanmar. PloS one 15, e0230802 (2020).

5. Vijaykrishna, D. et al. Evolutionary insights into the ecology of coronaviruses. J Virol 81, 4012–4020 (2007).

6. Masters, P.S. The molecular biology of coronaviruses. Adv Virus Res 66, 193–292 (2006).

7. Guan, Y. et al. Isolation and characterization of viruses related to the SARS coronavirus from animals in southern China. Science 302, 276–278 (2003).

8. Pyrc, K., Jebbink, M.F., Berkhout, B. & van der Hoek, L. Genome structure and transcriptional regulation of human coronavirus NL63. Virol J 1, 7 (2004).

9. Farsani, S.M. et al. The first complete genome sequences of clinical isolates of human coronavirus 229E. Virus Genes 45, 433–439 (2012).

10. St-Jean, J.R. et al. Human respiratory coronavirus OC43: genetic stability and neuroinvasion. J Virol 78, 8824–8834 (2004).

11. Lu, G., Wang, Q. & Gao, G.F. Bat-to-human: spike features determining ‘host jump’ of coronaviruses SARS-CoV, MERS-CoV, and beyond. Trends Microbiol 23, 468–478 (2015).

12. Wurzer, W.J., Obojes, K. & Vlasak, R. The sialate-4-O-acetylesterases of coronaviruses related to mouse hepatitis virus: a proposal to reorganize group 2 Coronaviridae. J Gen Virol 83, 395–402 (2002).

13. Agnihothram, S. et al. A mouse model for Betacoronavirus subgroup 2c using a bat coronavirus strain HKU5 variant. mBio 5, e00047–00014 (2014).

14. Menachery, V.D. et al. SARS-like WIV1-CoV poised for human emergence. Proc Natl Acad Sci U S A 113, 3048–3053 (2016).

15. Menachery, V.D. et al. Corrigendum: A SARS-like cluster of circulating bat coronaviruses shows potential for human emergence. Nat Med 22, 446 (2016).

16. Menachery, V.D. et al. A SARS-like cluster of circulating bat coronaviruses shows potential for human emergence. Nat Med 21, 1508–1513 (2015).

17. Liu, J. et al. Overlapping and discrete aspects of the pathology and pathogenesis of the emerging human pathogenic coronaviruses SARS-CoV, MERS-CoV, and 2019-nCoV. J Med Virol 92, 491–494 (2020).

18. Menachery, V.D., Graham, R.L. & Baric, R.S. Jumping species-a mechanism for coronavirus persistence and survival. Current opinion in virology 23, 1–7 (2017).

19. Zhou, P. et al. A pneumonia outbreak associated with a new coronavirus of probable bat origin. Nature 579, 270–273 (2020).

20. Koyama, T., Weeraratne, D., Snowdon, J.L. & Parida, L. Emergence of Drift Variants That May Affect COVID-19 Vaccine Development and Antibody Treatment. Pathogens 9 (2020).

21. Lun, Z.R. & Qu, L.H. Animal-to-human SARS-associated coronavirus transmission? Emerg Infect Dis 10, 959 (2004).

22. Layne, S.P., Hyman, J.M., Morens, D.M. & Taubenberger, J.K. New coronavirus outbreak: Framing questions for pandemic prevention. Science translational medicine 12 (2020).

23. Chowell, G. & Mizumoto, K. The COVID-19 pandemic in the USA: what might we expect? Lancet 395, 1093–1094 (2020).

24. Breslin, N. et al. COVID-19 infection among asymptomatic and symptomatic pregnant women: Two weeks of confirmed presentations to an affiliated pair of New York City hospitals. Am J Obstet Gynecol MFM, 100118 (2020).

25. An, P., Song, P., Wang, Y. & Liu, B. Asymptomatic Patients with Novel Coronavirus Disease (COVID-19). Balkan Med J (2020).

26. Chen, H. et al. Response of memory CD8+ T cells to severe acute respiratory syndrome (SARS) coronavirus in recovered SARS patients and healthy individuals. J Immunol 175, 591–598 (2005).

27. Rajendran, K. et al. Convalescent plasma transfusion for the treatment of COVID-19: Systematic review. J Med Virol (2020).

28. Pinto, D. et al. Cross-neutralization of SARS-CoV-2 by a human monoclonal SARS-CoV antibody. Nature (2020).

29. Ni, L. et al. Detection of SARS-CoV-2-Specific Humoral and Cellular Immunity in COVID-19 Convalescent Individuals. Immunity (2020).

30. Lee, C.Y., Lin, R.T.P., Renia, L. & Ng, L.F.P. Serological Approaches for COVID-19: Epidemiologic Perspective on Surveillance and Control. Front Immunol 11, 879 (2020).

31. Azkur, A.K. et al. Immune response to SARS-CoV-2 and mechanisms of immunopathological changes in COVID-19. Allergy (2020).

32. Ni, L. et al. Detection of SARS-CoV-2-Specific Humoral and Cellular Immunity in COVID-19 Convalescent Individuals. Immunity 52, 971–977 e973 (2020).

33. Mehta, P. et al. COVID-19: consider cytokine storm syndromes and immunosuppression. Lancet 395, 1033–1034 (2020).

34. Vaninov, N. In the eye of the COVID-19 cytokine storm. Nat Rev Immunol (2020).

35. Ma, J. et al. Potential effect of blood purification therapy in reducing cytokine storm as a late complication of critically ill COVID-19. Clin Immunol 214, 108408 (2020).

36. Henderson, L.A. et al. On the alert for cytokine storm: Immunopathology in COVID-19. Arthritis & rheumatology (2020).

37. Walsh, K.B. et al. Animal model of respiratory syncytial virus: CD8+ T cells cause a cytokine storm that is chemically tractable by sphingosine-1-phosphate 1 receptor agonist therapy. J Virol 88, 6281–6293 (2014).

38. Matheu, M.P. et al. Three phases of CD8 T cell response in the lung following H1N1 influenza infection and sphingosine 1 phosphate agonist therapy. PloS one 8, e58033 (2013).

39. Walsh, K.B. et al. Suppression of cytokine storm with a sphingosine analog provides protection against pathogenic influenza virus. Proc Natl Acad Sci U S A 108, 12018–12023 (2011).

40. Korber, B. et al. Tracking Changes in SARS-CoV-2 Spike: Evidence that D614G Increases Infectivity of the COVID-19 Virus. Cell (2020).

41. Li, X. et al. Emergence of SARS-CoV-2 through Recombination and Strong Purifying Selection. bioRxiv (2020).

42. Zheng, M. & Song, L. Novel antibody epitopes dominate the antigenicity of spike glycoprotein in SARS-CoV-2 compared to SARS-CoV. Cell Mol Immunol (2020).

43. Walls, A.C. et al. Structure, Function, and Antigenicity of the SARS-CoV-2 Spike Glycoprotein. Cell (2020).

44. Tilocca, B. et al. Molecular basis of COVID-19 relationships in different species: a one health perspective. Microbes Infect (2020).

45. Grifoni, A. et al. A Sequence Homology and Bioinformatic Approach Can Predict Candidate Targets for Immune Responses to SARS-CoV-2. Cell Host Microbe 27, 671–680 e672 (2020).

46. Bhattacharya, M., et al. Development of epitope-based peptide vaccine against novel coronavirus 2019 (SARS-COV-2): Immunoinformatics approach. J Med Virol (2020).

47. Baruah, V. & Bose, S. Immunoinformatics-aided identification of T cell and B cell epitopes in the surface glycoprotein of 2019-nCoV. J Med Virol 92, 495–500 (2020).

48. Ahmed, S.F., Quadeer, A.A. & McKay, M.R. Preliminary Identification of Potential Vaccine Targets for the COVID-19 Coronavirus (SARS-CoV-2) Based on SARS-CoV Immunological Studies. Viruses 12 (2020).

49. Vita, R. et al. The Immune Epitope Database (IEDB): 2018 update. Nucleic Acids Res 47, D339–D343 (2019).

50. Sette, A. & Sidney, J. Nine major HLA class I supertypes account for the vast preponderance of HLA-A and -B polymorphism. Immunogenetics 50, 201–212 (1999).

51. Sette, A. & Sidney, J. HLA supertypes and supermotifs: a functional perspective on HLA polymorphism. Curr Opin Immunol 10, 478–482 (1998).

52. Hertz, T. & Yanover, C. Identifying HLA supertypes by learning distance functions. Bioinformatics 23, e148–155 (2007).

53. Chentoufi, A.A. et al. HLA-A*0201-restricted CD8+ cytotoxic T lymphocyte epitopes identified from herpes simplex virus glycoprotein D. J Immunol 180, 426–437 (2008).

54. Jespersen, M.C., Peters, B., Nielsen, M. & Marcatili, P. BepiPred-2.0: improving sequence-based B-cell epitope prediction using conformational epitopes. Nucleic Acids Res 45, W24–W29 (2017).

55. Monto, A.S. et al. Coronavirus Occurrence and Transmission Over 8 Years in the HIVE Cohort of Households in Michigan. The Journal of infectious diseases 222, 9–16 (2020).

56. Samandary, S. et al. Associations of HLA-A, HLA-B and HLA-C alleles frequency with prevalence of herpes simplex virus infections and diseases across global populations: implication for the development of an universal CD8+ T-cell epitope-based vaccine. Hum Immunol 75, 715–729 (2014).

57. Sallusto, F., Lenig, D., Forster, R., Lipp, M. & Lanzavecchia, A. Two subsets of memory T lymphocytes with distinct homing potentials and effector functions. Nature 401, 708–712 (1999).

58. Sempowski, G.D., Saunders, K.O., Acharya, P., Wiehe, K.J. & Haynes, B.F. Pandemic Preparedness: Developing Vaccines and Therapeutic Antibodies For COVID-19. Cell (2020).

59. Lurie, N., Saville, M., Hatchett, R. & Halton, J. Developing Covid-19 Vaccines at Pandemic Speed. The New England journal of medicine 382, 1969–1973 (2020).

60. Grifoni, A. et al. Targets of T Cell Responses to SARS-CoV-2 Coronavirus in Humans with COVID-19 Disease and Unexposed Individuals. Cell (2020).

61. Ng, O.W. & Tan, Y.J. Understanding bat SARS-like coronaviruses for the preparation of future coronavirus outbreaks - Implications for coronavirus vaccine development. Hum Vaccin Immunother 13, 186–189 (2017).

62. Wu, A. et al. Genome Composition and Divergence of the Novel Coronavirus (2019-nCoV) Originating in China. Cell Host Microbe 27, 325–328 (2020).

63. Martin, J.E. et al. A SARS DNA vaccine induces neutralizing antibody and cellular immune responses in healthy adults in a Phase I clinical trial. Vaccine 26, 6338–6343 (2008).

64. He, Y. & Jiang, S. Vaccine design for severe acute respiratory syndrome coronavirus. Viral Immunol 18, 327–332 (2005).

65. He, Y., Zhou, Y., Siddiqui, P. & Jiang, S. Inactivated SARS-CoV vaccine elicits high titers of spike protein-specific antibodies that block receptor binding and virus entry. Biochem Biophys Res Commun 325, 445–452 (2004).

66. Sidney, J., Peters, B., Frahm, N., Brander, C. & Sette, A. HLA class I supertypes: a revised and updated classification. BMC Immunol 9, 1 (2008).

67. Correale, P. et al. HLA-B*44 and C*01 Prevalence Correlates with Covid19 Spreading across Italy. Int J Mol Sci 21 (2020).

68. Correale, P. et al. HLA Expression Correlates to the Risk of Immune Checkpoint Inhibitor-Induced Pneumonitis. Cells 9 (2020).

69. Peng, Y. et al. Broad and strong memory CD4(+) and CD8(+) T cells induced by SARS-CoV-2 in UK convalescent individuals following COVID-19. Nat Immunol (2020).

70. Pedersen, S.F. & Ho, Y.C. SARS-CoV-2: A Storm is Raging. J Clin Invest (2020).

71. Fung, S.Y., Yuen, K.S., Ye, Z.W., Chan, C.P. & Jin, D.Y. A tug-of-war between severe acute respiratory syndrome coronavirus 2 and host antiviral defence: lessons from other pathogenic viruses. Emerg Microbes Infect 9, 558–570 (2020).

72. Akhmerov, A. & Marban, E. COVID-19 and the Heart. Circ Res (2020).

73. Cyranoski, D. The biggest mystery: what it will take to trace the coronavirus source. Nature (2020).

74. Dallavilla, T. et al. Bioinformatic analysis indicates that SARS-CoV-2 is unrelated to known artificial coronaviruses. Eur Rev Med Pharmacol Sci 24, 4558–4564 (2020).

75. Haagmans, B.L. et al. Middle East respiratory syndrome coronavirus in dromedary camels: an outbreak investigation. Lancet Infect Dis 14, 140–145 (2014).

76. Wang, L.F., Anderson, D.E., Mackenzie, J.S. & Merson, M.H. From Hendra to Wuhan: what has been learned in responding to emerging zoonotic viruses. Lancet 395, e33–e34 (2020).

77. Plowright, R.K. et al. Transmission or Within-Host Dynamics Driving Pulses of Zoonotic Viruses in Reservoir-Host Populations. PLoS Negl Trop Dis 10, e0004796 (2016).

78. Eng, C.L., Tong, J.C. & Tan, T.W. Distinct Host Tropism Protein Signatures to Identify Possible Zoonotic Influenza A Viruses. PloS one 11, e0150173 (2016).

79. Simons, R.R., Gale, P., Horigan, V., Snary, E.L. & Breed, A.C. Potential for introduction of bat-borne zoonotic viruses into the EU: a review. Viruses 6, 2084–2121 (2014).

80. Crooke, S.N., Ovsyannikova, I.G., Kennedy, R.B. & Poland, G.A. Immunoinformatic identification of B cell and T cell epitopes in the SARS-CoV-2 proteome. Scientific reports 10, 14179 (2020).

81. Kar, T. et al. A candidate multi-epitope vaccine against SARS-CoV-2. Scientific reports 10, 10895 (2020).

82. Mateus, J. et al. Selective and cross-reactive SARS-CoV-2 T cell epitopes in unexposed humans. Science (2020).

83. Grifoni, A. et al. Targets of T Cell Responses to SARS-CoV-2 Coronavirus in Humans with COVID-19 Disease and Unexposed Individuals. Cell 181, 1489–1501 e1415 (2020).

84. Baggio, S. et al. SARS-CoV-2 viral load in the upper respiratory tract of children and adults with early acute COVID-19. Clin Infect Dis (2020).

85. Moratto, D. et al. Immune response in children with COVID-19 is characterized by lower levels of T-cell activation than infected adults. Eur J Immunol (2020).

86. Mansour, M. et al. Prevalence of SARS-CoV-2 Antibodies Among Healthcare Workers at a Tertiary Academic Hospital in New York City. J Gen Intern Med (2020).

87. Voo, T.C., Clapham, H. & Tam, C.C. Ethical implementation of ‘immunity passports’ during the COVID-19 pandemic. The Journal of infectious diseases (2020).

88. Kofler, N. & Baylis, F. Ten reasons why immunity passports are a bad idea. Nature 581, 379–381 (2020).

89. Atyeo, C. et al. Distinct Early Serological Signatures Track with SARS-CoV-2 Survival. Immunity 53, 524–532 e524 (2020).

90. Wang, B. et al. Bivalent binding of a fully human IgG to the SARS-CoV-2 spike proteins reveals mechanisms of potent neutralization. bioRxiv (2020).

91. Benvenuto, D. et al. Evidence for mutations in SARS-CoV-2 Italian isolates potentially affecting virus transmission. J Med Virol (2020).

92. Ong, E.Z. et al. A Dynamic Immune Response Shapes COVID-19 Progression. Cell Host Microbe 27, 879–882 e872 (2020).

93. Le Bert, N. et al. SARS-CoV-2-specific T cell immunity in cases of COVID-19 and SARS, and uninfected controls. Nature 584, 457–462 (2020).

94. Welsh, R.M. & Selin, L.K. No one is naive: the significance of heterologous T-cell immunity. Nat Rev Immunol 2, 417–426 (2002).

95. Becker, M.M. et al. Synthetic recombinant bat SARS-like coronavirus is infectious in cultured cells and in mice. Proc Natl Acad Sci U S A 105, 19944–19949 (2008).

96. Bui, H.H. et al. Predicting population coverage of T-cell epitope-based diagnostics and vaccines. BMC bioinformatics 7, 153 (2006).

97. Oh, S. et al. Human CTLs to wild-type and enhanced epitopes of a novel prostate and breast tumor-associated protein, TARP, lyse human breast cancer cells. Cancer Res 64, 2610–2618 (2004).

98. Stuber, G. et al. Identification of wild-type and mutant p53 peptides binding to HLA-A2 assessed by a peptide loading-deficient cell line assay and a novel major histocompatibility complex class I peptide binding assay. Eur J Immunol 24, 765–768 (1994).

99. Khan, A.A. et al. Phenotypic and functional characterization of herpes simplex virus glycoprotein B epitope-specific effector and memory CD8+ T cells from symptomatic and asymptomatic individuals with ocular herpes. J Virol 89, 3776–3792 (2015).

100. Srivastava, R. et al. The Herpes Simplex Virus Latency-Associated Transcript Gene Is Associated with a Broader Repertoire of Virus-Specific Exhausted CD8+ T Cells Retained within the Trigeminal Ganglia of Latently Infected HLA Transgenic Rabbits. J Virol 90, 3913–3928 (2016).

101. Khan, A.A. et al. Human Asymptomatic Epitope Peptide/CXCL10-Based Prime/Pull Vaccine Induces Herpes Simplex Virus-Specific Gamma Interferon-Positive CD107(+) CD8(+) T Cells That Infiltrate the Corneas and Trigeminal Ganglia of Humanized HLA Transgenic Rabbits and Protect against Ocular Herpes Challenge. J Virol 92 (2018).

