## Supplement Figures 1 to 5 for "Genome-Wide Asymptomatic B-Cell, CD4^+^ and CD8^+^ T-Cell Epitopes, that are Highly Conserved Between Human and Animal Coronaviruses, Identified from SARS-CoV-2 as Immune Targets for Pre-Emptive Pan-Coronavirus Vaccines"

### A Predicted population coverage (PPC) value of human CD8⁺ T cell epitopes

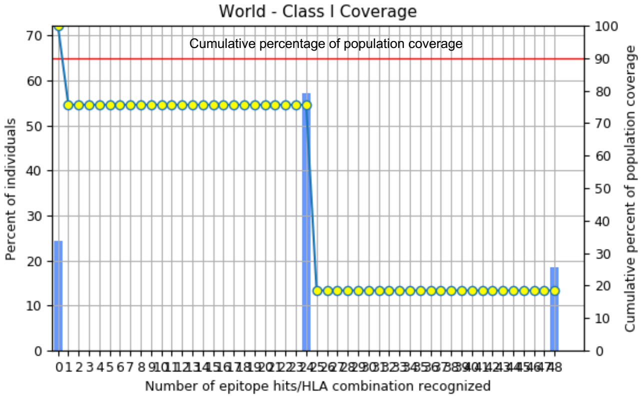


| Population/Ethnicity | Class I | | |
|---|---|---|---|
| | Cumulative Coverage | Average hit | pc90 |
| World | **75.66%** | 22.6 | 9.86 |

| Epitope | SARS-CoV-2 derived CD8⁺ Epitopes sequence | HLA binding alleles | Cumulative PPC value (%) |
|---|---|---|---|
| ORF1ab$_{84-92}$ | VMVELVAEL | HLA-A*01:01,HLA-A*02:01,HLA-A*03:01,HLA-A*11:01,HLA-A*23:01 | 75.66 |
| ORF1ab$_{1675-1683}$ | YLATALLTL | HLA-A*01:01,HLA-A*02:01,HLA-A*03:01,HLA-A*11:01,HLA-A*23:01 | 75.66 |
| ORF1ab$_{2210-2218}$ | CLEASFNYL | HLA-A*01:01,HLA-A*02:01,HLA-A*03:01,HLA-A*11:01,HLA-A*23:01 | 75.66 |
| ORF1ab$_{2363-2371}$ | WLMWLIINL | HLA-A*01:01,HLA-A*02:01,HLA-A*03:01,HLA-A*11:01,HLA-A*23:01 | 75.66 |
| ORF1ab$_{3013-3021}$ | SLPGVFCGV | HLA-A*01:01,HLA-A*02:01,HLA-A*03:01,HLA-A*11:01,HLA-A*23:01 | 75.66 |
| ORF1ab$_{3732-3740}$ | SMWALIISV | HLA-A*01:01,HLA-A*02:01,HLA-A*03:01,HLA-A*11:01,HLA-A*23:01 | 75.66 |
| ORF1ab$_{4283-4291}$ | YLASGGQPI | HLA-A*01:01,HLA-A*02:01,HLA-A*03:01,HLA-A*11:01,HLA-A*23:01 | 75.66 |
| ORF1ab$_{6419-6427}$ | YLDAYNMMI | HLA-A*01:01,HLA-A*02:01,HLA-A*03:01,HLA-A*11:01,HLA-A*23:01 | 75.66 |
| ORF1ab$_{6749-6757}$ | LLLDDFVEI | HLA-A*01:01,HLA-A*02:01,HLA-A*03:01,HLA-A*11:01,HLA-A*23:01 | 75.66 |
| S$_{2-10}$ | FVFLVLLPL | HLA-A*01:01,HLA-A*02:01,HLA-A*03:01,HLA-A*11:01,HLA-A*23:01 | 75.66 |
| S$_{691-699}$ | SIIAYTMSL | HLA-A*01:01,HLA-A*02:01,HLA-A*03:01,HLA-A*11:01,HLA-A*23:01 | 75.66 |
| S$_{958-966}$ | ALNTLVKQL | HLA-A*01:01,HLA-A*02:01,HLA-A*03:01,HLA-A*11:01,HLA-A*23:01 | 75.66 |
| S$_{976-984}$ | VLNDILSRL | HLA-A*01:01,HLA-A*02:01,HLA-A*03:01,HLA-A*11:01,HLA-A*23:01 | 75.66 |
| S$_{1220-1228}$ | FIAGLIAIV | HLA-A*01:01,HLA-A*02:01,HLA-A*03:01,HLA-A*11:01,HLA-A*23:01 | 75.66 |
| E$_{20-28}$ | FLAFVVFLL | HLA-A*01:01,HLA-A*02:01,HLA-A*03:01,HLA-A*11:01,HLA-A*23:01 | 75.66 |
| E$_{26-34}$ | FLLVTLAIL | HLA-A*01:01,HLA-A*02:01,HLA-A*03:01,HLA-A*11:01,HLA-A*23:01 | 75.66 |
| M$_{52-60}$ | IFLWLLWPV | HLA-A*01:01,HLA-A*02:01,HLA-A*03:01,HLA-A*11:01,HLA-A*23:01 | 75.66 |
| M$_{89-97}$ | GLMWLSYFI | HLA-A*01:01,HLA-A*02:01,HLA-A*03:01,HLA-A*11:01,HLA-A*23:01 | 75.66 |
| ORF6$_{4-11}$ | HLVDFQVTI | HLA-A*01:01,HLA-A*02:01,HLA-A*03:01,HLA-A*11:01,HLA-A*23:01 | 75.66 |
| ORF7b$_{26-34}$ | IIFWFSLEL | HLA-A*01:01,HLA-A*02:01,HLA-A*03:01,HLA-A*11:01,HLA-A*23:01 | 75.66 |
| ORF8a$_{31-39}$ | YVVDDPCPI | HLA-A*01:01,HLA-A*02:01,HLA-A*03:01,HLA-A*11:01,HLA-A*23:01 | 75.66 |
| ORF8a$_{73-81}$ | YIDIGNYTV | HLA-A*01:01,HLA-A*02:01,HLA-A*03:01,HLA-A*11:01,HLA-A*23:01 | 75.66 |
| ORF10$_{3-11}$ | YINVFAFPF | HLA-A*01:01,HLA-A*02:01,HLA-A*03:01,HLA-A*11:01,HLA-A*23:01 | 75.66 |
| ORF10$_{5-13}$ | NVFAFPFTI | HLA-A*01:01,HLA-A*02:01,HLA-A*03:01,HLA-A*11:01,HLA-A*23:01 | 75.66 |

### B Predicted population coverage (PPC) value of human CD4⁺ T cell epitopes

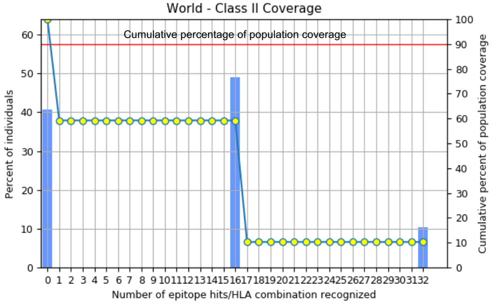


| Population/ Ethnicity | Class I | | |
|---|---|---|---|
| | Cumulative Coverage | Average hit | pc90 |
| World | **59.25%** | 11.13 | 3.96 |

| Epitope | SARS-CoV-2 derived CD4⁺ Epitopes sequence | HLA binding alleles | Cumulative PPC (%) |
|---|---|---|---|
| ORF1a$_{1350-1365}$ | KSAFYILPSIISNEK | HLA-DRB1*01:01, HLA-DRB1*11:01, HLA-DRB1*15:01, HLA-DRB1*03:01, HLA-DRB1*04:01 | 59.25 |
| ORF1a$_{1801-1815}$ | ESPFVMMSAPPAQYE | HLA-DRB1*01:01, HLA-DRB1*11:01, HLA-DRB1*15:01, HLA-DRB1*03:01, HLA-DRB1*04:01 | 59.25 |
| ORF1ab$_{5019-5033}$ | PNMLRIMASLVLARK | HLA-DRB1*01:01, HLA-DRB1*11:01, HLA-DRB1*15:01, HLA-DRB1*03:01, HLA-DRB1*04:01 | 59.25 |
| ORF1ab$_{6088-6102}$ | RIKVQMLSDTLKNL | HLA-DRB1*01:01, HLA-DRB1*11:01, HLA-DRB1*15:01, HLA-DRB1*03:01, HLA-DRB1*04:01 | 59.25 |
| ORF1ab$_{6420-6434}$ | LDAYNMMISAGFSLW | HLA-DRB1*01:01, HLA-DRB1*11:01, HLA-DRB1*15:01, HLA-DRB1*03:01, HLA-DRB1*04:01 | 59.25 |
| S$_{1-13}$ | MFVFLVLLPLVSS | HLA-DRB1*01:01, HLA-DRB1*11:01, HLA-DRB1*15:01, HLA-DRB1*03:01, HLA-DRB1*04:01 | 59.25 |
| E$_{20-34}$ | FLAFVVFLLVTLAIL | HLA-DRB1*01:01, HLA-DRB1*11:01, HLA-DRB1*15:01, HLA-DRB1*03:01, HLA-DRB1*04:01 | 59.25 |
| E$_{26-40}$ | FLLVTLAILTALRLC | HLA-DRB1*01:01, HLA-DRB1*11:01, HLA-DRB1*15:01, HLA-DRB1*03:01, HLA-DRB1*04:01 | 59.25 |
| M$_{176-190}$ | LSYYKLGASQRVAGD | HLA-DRB1*01:01, HLA-DRB1*11:01, HLA-DRB1*15:01, HLA-DRB1*03:01, HLA-DRB1*04:01 | 59.25 |
| ORF6$_{12-26}$ | AEILLIIMRTFKVSI | HLA-DRB1*01:01, HLA-DRB1*11:01, HLA-DRB1*15:01, HLA-DRB1*03:01, HLA-DRB1*04:01 | 59.25 |
| ORF7a$_{1-15}$ | MKIILFLALITLATC | HLA-DRB1*01:01, HLA-DRB1*11:01, HLA-DRB1*15:01, HLA-DRB1*03:01, HLA-DRB1*04:01 | 59.25 |
| ORF7a$_{3-17}$ | IIFLALITLATCEL | HLA-DRB1*01:01, HLA-DRB1*11:01, HLA-DRB1*15:01, HLA-DRB1*03:01, HLA-DRB1*04:01 | 59.25 |
| ORF7a$_{98-112}$ | SPIFLIVAAIVFITL | HLA-DRB1*01:01, HLA-DRB1*11:01, HLA-DRB1*15:01, HLA-DRB1*03:01, HLA-DRB1*04:01 | 59.25 |
| ORF7b$_{8-22}$ | DFYLCFLAFLLFLVL | HLA-DRB1*01:01, HLA-DRB1*11:01, HLA-DRB1*15:01, HLA-DRB1*03:01, HLA-DRB1*04:01 | 59.25 |
| ORF$_{8b1-15}$ | MKFLVFLGIITTVAA | HLA-DRB1*01:01, HLA-DRB1*11:01, HLA-DRB1*15:01, HLA-DRB1*03:01, HLA-DRB1*04:01 | 59.25 |
| N$_{388-4031}$ | KQQTVTLLPAADLDDF | HLA-DRB1*01:01, HLA-DRB1*11:01, HLA-DRB1*15:01, HLA-DRB1*03:01, HLA-DRB1*04:01 | 59.25 |

### C Predicted population coverage (PPC) value of human $CD8^+$ T cell epitopes in Pan-Coronavirus Vaccine candidate # 1

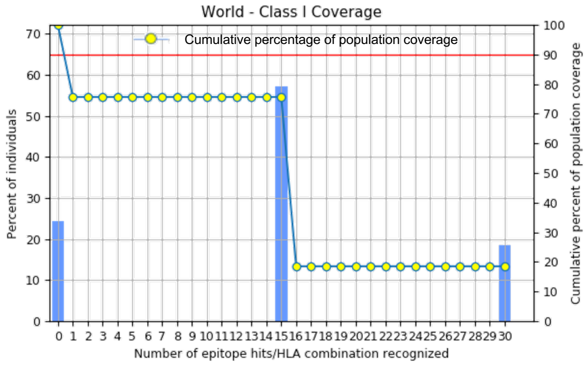


| Population/Ethnicity | Class I | | |
|---|---|---|---|
| | Cumulative Coverage | Average hit | pc90 |
| World | **75.66%** | 14.13 | 6.16 |

| Epitope | SARS-CoV-2 derived $CD8^+$ Epitopes sequence | HLA binding alleles | Cumulative PPC (%) |
|---|---|---|---|
| ORF1ab$_{3013-3021}$ | SLPGVFCGV | HLA-A*01:01,HLA-A*02:01,HLA-A*03:01,HLA-A*11:01,HLA-A*23:01 | 75.66 |
| ORF1ab$_{1675-1683}$ | YLATALLTL | HLA-A*01:01,HLA-A*02:01,HLA-A*03:01,HLA-A*11:01,HLA-A*23:01 | 75.66 |
| ORF10$_{3-11}$ | YINVFAFPF | HLA-A*01:01,HLA-A*02:01,HLA-A*03:01,HLA-A*11:01,HLA-A*23:01 | 75.66 |
| E$_{26-34}$ | FLLNKEMYL | HLA-A*01:01,HLA-A*02:01,HLA-A*03:01,HLA-A*11:01,HLA-A*23:01 | 75.66 |
| S$_{2-10}$ | FLAFVVFLL | HLA-A*01:01,HLA-A*02:01,HLA-A*03:01,HLA-A*11:01,HLA-A*23:01 | 75.66 |
| S$_{1220-1228}$ | FIAGLIAIV | HLA-A*01:01,HLA-A*02:01,HLA-A*03:01,HLA-A*11:01,HLA-A*23:01 | 75.66 |
| ORF1ab$_{2363-2371}$ | WLMWLIINL | HLA-A*01:01,HLA-A*02:01,HLA-A*03:01,HLA-A*11:01,HLA-A*23:01 | 75.66 |
| ORF8a$_{73-81}$ | YIDIGNYTV | HLA-A*01:01,HLA-A*02:01,HLA-A*03:01,HLA-A*11:01,HLA-A*23:01 | 75.66 |
| ORF7b$_{26-34}$ | IIFWFSLEL | HLA-A*01:01,HLA-A*02:01,HLA-A*03:01,HLA-A*11:01,HLA-A*23:01 | 75.66 |
| ORF10$_{5-13}$ | NVFAFPFTI | HLA-A*01:01,HLA-A*02:01,HLA-A*03:01,HLA-A*11:01,HLA-A*23:01 | 75.66 |
| S$_{2-10}$ | FVFLVLLPL | HLA-A*01:01,HLA-A*02:01,HLA-A*03:01,HLA-A*11:01,HLA-A*23:01 | 75.66 |
| S$_{958-966}$ | ALNTLVKQL | HLA-A*01:01,HLA-A*02:01,HLA-A*03:01,HLA-A*11:01,HLA-A*23:01 | 75.66 |
| ORF1ab$_{5470-5478}$ | KLSYGIATV | HLA-A*01:01,HLA-A*02:01,HLA-A*03:01,HLA-A*11:01,HLA-A*23:01 | 75.66 |
| S$_{1000-1008}$ | RLQSLQTYV | HLA-A*01:01,HLA-A*02:01,HLA-A*03:01,HLA-A*11:01,HLA-A*23:01 | 75.66 |
| ORF1ab$_{6749-6757}$ | LLLDDFVEI | HLA-A*01:01,HLA-A*02:01,HLA-A*03:01,HLA-A*11:01,HLA-A*23:01 | 75.66 |

### D Predicted population coverage (PPC) value of human $CD4^+$ T cell epitopes in Pan-Coronavirus Vaccine candidate # 1

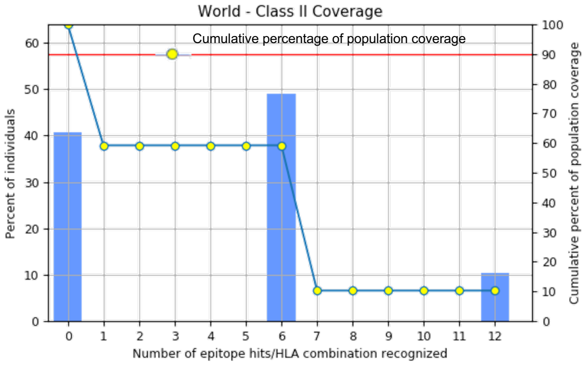


| Population/Ethnicity | Class II | | |
|---|---|---|---|
| | Cumulative Coverage | Average hit | pc90 |
| World | **59.25%** | 4.17 | 1.47 |

| Epitope | SARS-CoV-2 derived $CD4^+$ Epitopes sequence | HLA binding alleles | Cumulative PPC (%) |
|---|---|---|---|
| ORF1a$_{1350-1365}$ | KSAFYILPSIISNEK | HLA-DRB1*01:01, HLA-DRB1*11:01, HLA-DRB1*15:01, HLA-DRB1*03:01, HLA-DRB1*04:01 | 59.25 |
| ORF1ab$_{5019-5033}$ | PNMLRIMASLVLARK | HLA-DRB1*01:01, HLA-DRB1*11:01, HLA-DRB1*15:01, HLA-DRB1*03:01, HLA-DRB1*04:01 | 59.25 |
| M$_{176-190}$ | LSYYKLGASQRVAGD | HLA-DRB1*01:01, HLA-DRB1*11:01, HLA-DRB1*15:01, HLA-DRB1*03:01, HLA-DRB1*04:01 | 59.25 |
| ORF6$_{12-26}$ | AEILLIIMRTFKVSI | HLA-DRB1*01:01, HLA-DRB1*11:01, HLA-DRB1*15:01, HLA-DRB1*03:01, HLA-DRB1*04:01 | 59.25 |
| ORF7b$_{8-22}$ | DFYLCFLAFLLFLVL | HLA-DRB1*01:01, HLA-DRB1*11:01, HLA-DRB1*15:01, HLA-DRB1*03:01, HLA-DRB1*04:01 | 59.25 |
| ORF$_{8b1-15}$ | MKFLVFLGIITTVAA | HLA-DRB1*01:01, HLA-DRB1*11:01, HLA-DRB1*15:01, HLA-DRB1*03:01, HLA-DRB1*04:01 | 59.25 |

Prakash *et al.* Supplemental Fig. S2

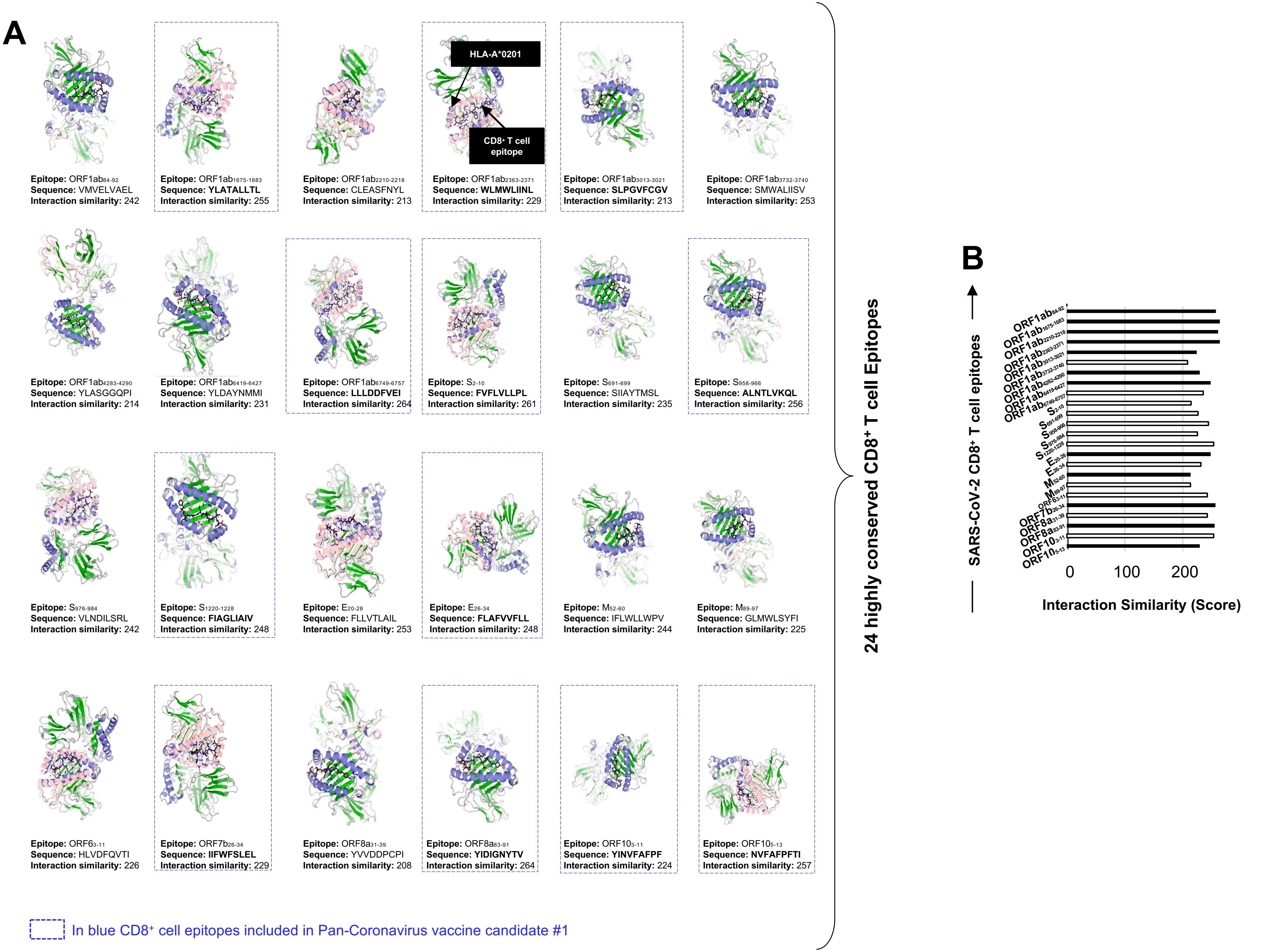

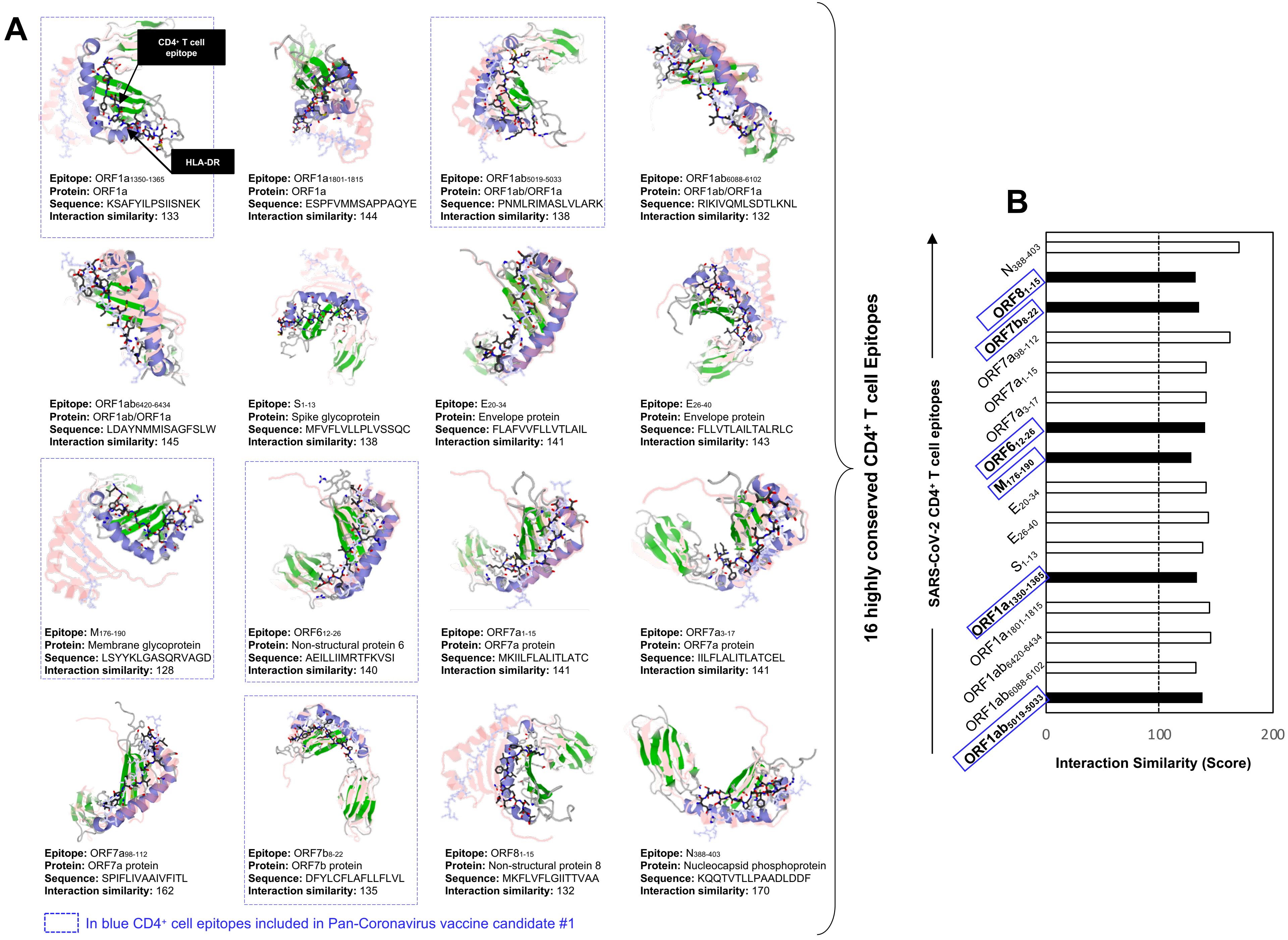
A
CD4+ T cell epitope
HLA-DR
Epitope: ORF1a1350-1365
Protein: ORF1a
Sequence: KSAFYILPSIISNEK
Interaction similarity: 133
Epitope: ORF1a1801-1815
Protein: ORF1a
Sequence: ESPFVMMSAPPAQYE
Interaction similarity: 144
Epitope: ORF1ab5019-5033
Protein: ORF1ab/ORF1a
Sequence: PNMLRIMASLVLARK
Interaction similarity: 138
Epitope: ORF1ab6088-6102
Protein: ORF1ab/ORF1a
Sequence: RIKIVQMLSDTLKNL
Interaction similarity: 132
Epitope: ORF1ab6420-6434
Protein: ORF1ab/ORF1a
Sequence: LDAYNMMISAGFSLW
Interaction similarity: 145
Epitope: S1-13
Protein: Spike glycoprotein
Sequence: MFVFLVLLPLVSSQC
Interaction similarity: 138
Epitope: E20-34
Protein: Envelope protein
Sequence: FLAFVVFLLVTLAIL
Interaction similarity: 141
Epitope: E26-40
Protein: Envelope protein
Sequence: FLLVTLAILTALRLC
Interaction similarity: 143
Epitope: M176-190
Protein: Membrane glycoprotein
Sequence: LSYYKLGASQRVAGD
Interaction similarity: 128
Epitope: ORF612-26
Protein: Non-structural protein 6
Sequence: AEILLIIMRTFKVSI
Interaction similarity: 140
Epitope: ORF7a1-15
Protein: ORF7a protein
Sequence: MKIILFLALITLATC
Interaction similarity: 141
Epitope: ORF7a3-17
Protein: ORF7a protein
Sequence: IILFLALITLATCEL
Interaction similarity: 141
Epitope: ORF7a98-112
Protein: ORF7a protein
Sequence: SPIFLIVAAIVFITL
Interaction similarity: 162
Epitope: ORF7b8-22
Protein: ORF7b protein
Sequence: DFYLCFLAFLLFLVL
Interaction similarity: 135
Epitope: ORF81-15
Protein: Non-structural protein 8
Sequence: MKFLVFLGIITTVAA
Interaction similarity: 132
Epitope: N388-403
Protein: Nucleocapsid phosphoprotein
Sequence: KQQTVTLLPAADLDDF
Interaction similarity: 170
In blue CD4+ cell epitopes included in Pan-Coronavirus vaccine candidate #1
16 highly conserved CD4+ T cell Epitopes
B
SARS-CoV-2 CD4+ T cell epitopes
N388-403
ORF81-15
ORF7b8-22
ORF7a98-112
ORF7a1-15
ORF7a3-17
ORF612-26
M176-190
E20-34
E26-40
S1-13
ORF1a1350-1365
ORF1a1801-1815
ORF1ab6420-6434
ORF1ab6088-6102
ORF1ab5019-5033
0
100
200
Interaction Similarity (Score)

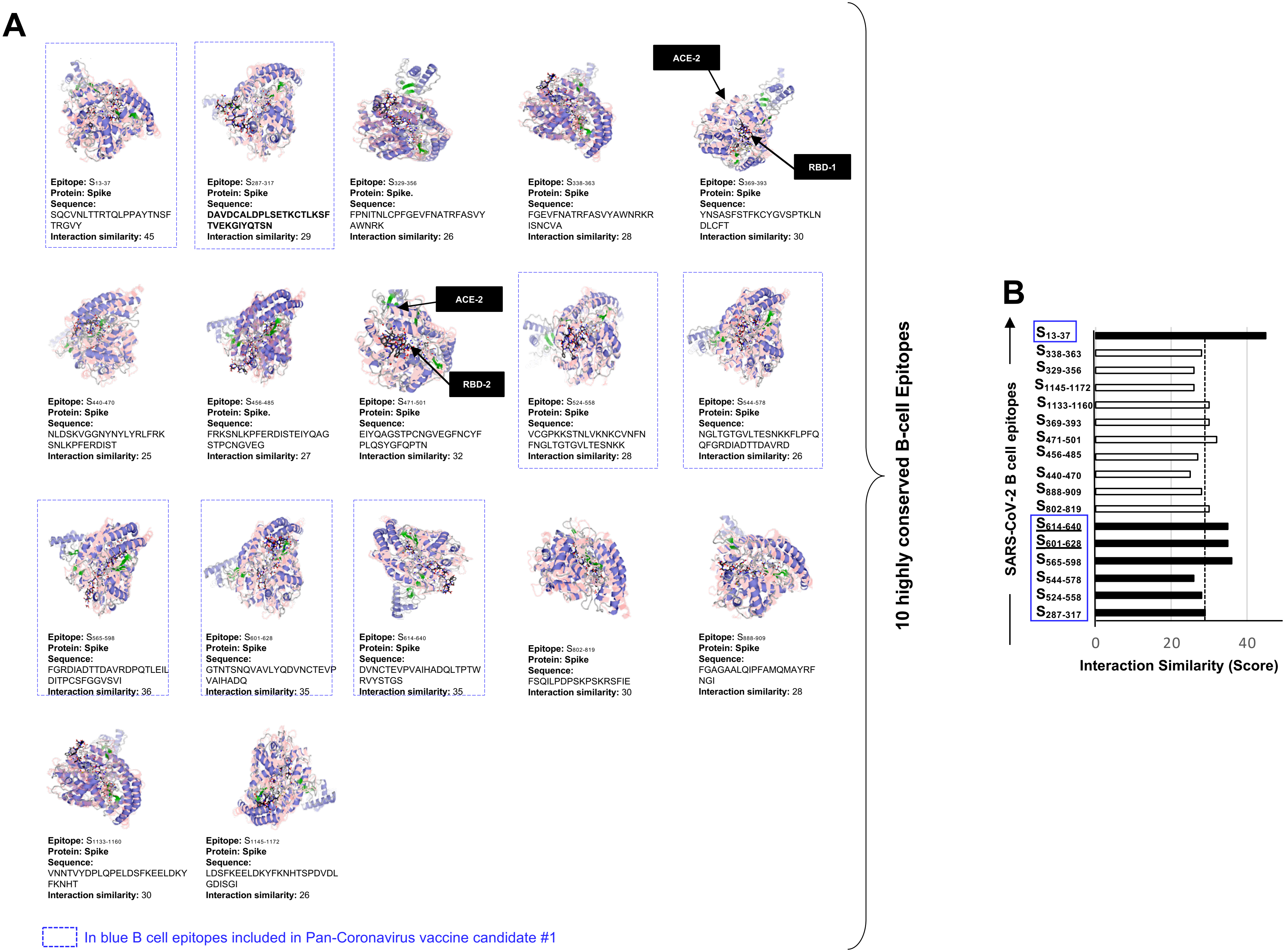
A
ACE-2
RBD-1
Epitope: S13-37
Protein: Spike
Sequence:
SQCVNLTTRTQLPPAYTNSF
TRGVY
Interaction similarity: 45
Epitope: S287-317
Protein: Spike
Sequence:
DAVDCALDPLSETKCTLKSF
TVEKGIYQTSN
Interaction similarity: 29
Epitope: S329-356
Protein: Spike.
Sequence:
FPNITNLCPFGEVFNATRFASVY
AWNRK
Interaction similarity: 26
Epitope: S338-363
Protein: Spike
Sequence:
FGEVFNATRFASVYAWNRKR
ISNCVA
Interaction similarity: 28
Epitope: S369-393
Protein: Spike
Sequence:
YNSASFSTFKCYGVSPTKLN
DLCFT
Interaction similarity: 30
ACE-2
RBD-2
Epitope: S440-470
Protein: Spike
Sequence:
NLDSKVGGNYNYLYRLFRK
SNLKPFERDIST
Interaction similarity: 25
Epitope: S456-485
Protein: Spike.
Sequence:
FRKSNLKPFERDISTEIYQAG
STPCNGVEG
Interaction similarity: 27
Epitope: S471-501
Protein: Spike
Sequence:
EIYQAGSTPCNGVEGFNCYF
PLQSYGFQPTN
Interaction similarity: 32
Epitope: S524-558
Protein: Spike
Sequence:
VCGPKKSTNLVKNKCVNFN
FNGLTGTGVLTESNKK
Interaction similarity: 28
Epitope: S544-578
Protein: Spike
Sequence:
NGLTGTGVLTESNKKFLPFQ
QFGRDIADTTDAVRD
Interaction similarity: 26
Epitope: S565-598
Protein: Spike
Sequence:
FGRDIADTTDAVRDPQTLEIL
DITPCSFGGVSVI
Interaction similarity: 36
Epitope: S601-628
Protein: Spike
Sequence:
GTNTSNQVAVLYQDVNCTEVP
VAIHADQ
Interaction similarity: 35
Epitope: S614-640
Protein: Spike
Sequence:
DVNCTEVPVAIHADQLTPTW
RVYSTGS
Interaction similarity: 35
Epitope: S802-819
Protein: Spike
Sequence:
FSQILPDPSKPSKRSFIE
Interaction similarity: 30
Epitope: S888-909
Protein: Spike
Sequence:
FGAGAALQIPFAMQMAYRF
NGI
Interaction similarity: 28
Epitope: S1133-1160
Protein: Spike
Sequence:
VNNTVYDPLQPELDSFKEELDKY
FKNHT
Interaction similarity: 30
Epitope: S1145-1172
Protein: Spike
Sequence:
LDSFKEELDKYFKNHTSPDVDL
GDISGI
Interaction similarity: 26
In blue B cell epitopes included in Pan-Coronavirus vaccine candidate #1
10 highly conserved B-cell Epitopes
B
SARS-CoV-2 B cell epitopes
S13-37
S338-363
S329-356
S1145-1172
S1133-1160
S369-393
S471-501
S456-485
S440-470
S888-909
S802-819
S614-640
S601-628
S565-598
S544-578
S524-558
S287-317
0
20
40
Interaction Similarity (Score)

Prakash *et al.* Supplemental Fig. S5

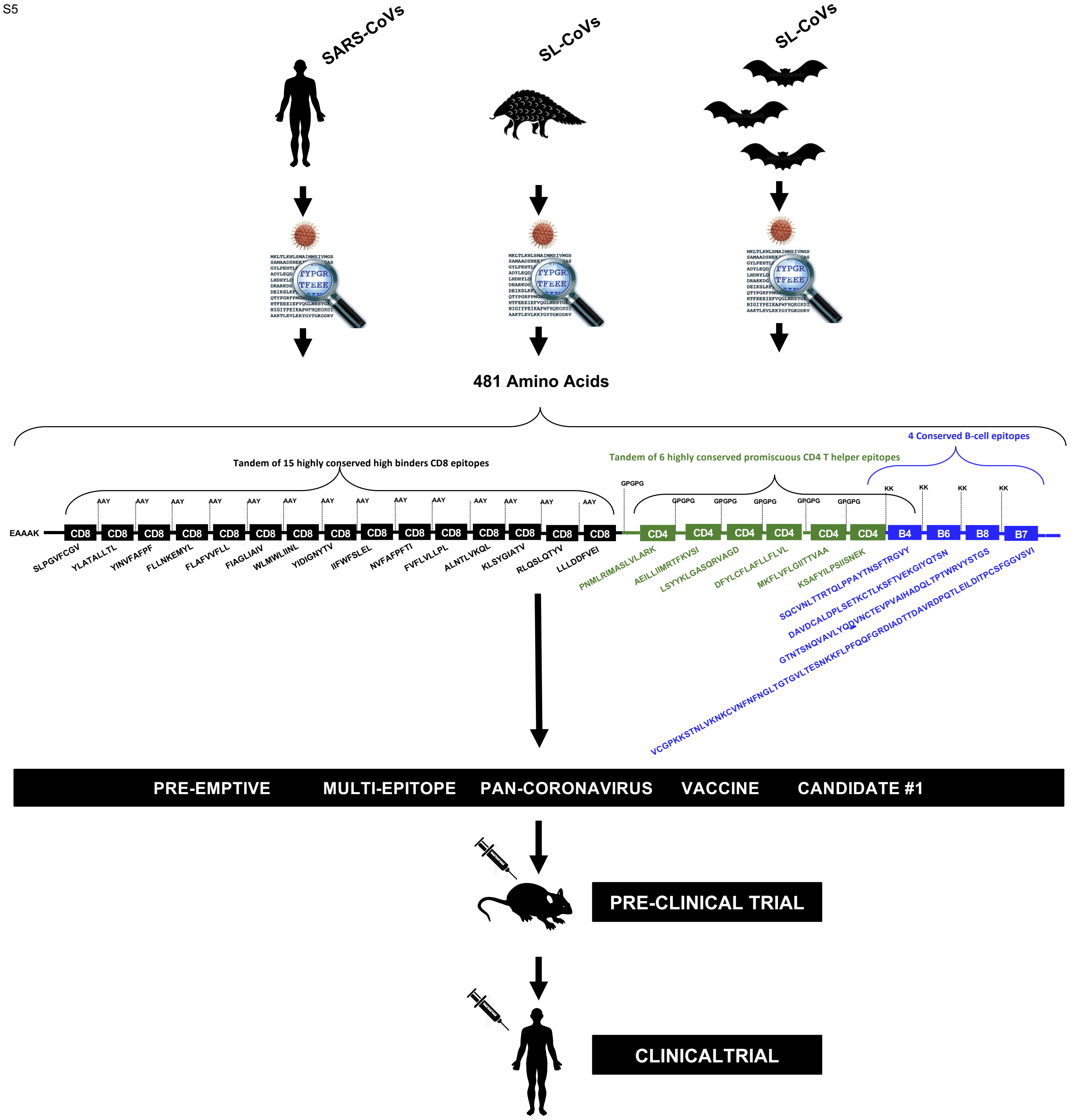
